# Phase separation by ssDNA binding protein controlled *via* protein-protein and protein-DNA interactions

**DOI:** 10.1101/797431

**Authors:** Gábor M. Harami, Zoltán J. Kovács, Rita Pancsa, János Pálinkás, Veronika Baráth, Krisztián Tárnok, András Málnási-Csizmadia, Mihály Kovács

## Abstract

Bacterial single stranded (ss) DNA-binding proteins (SSB) are essential for the replication and maintenance of the genome. SSBs share a conserved ssDNA-binding domain, a less conserved intrinsically disordered linker (IDL) and a highly conserved C-terminal peptide (CTP) motif that mediates a wide array of protein-protein interactions with DNA-metabolizing proteins. Here we show that the *E. coli* SSB protein forms liquid-liquid phase separated condensates in cellular-like conditions through multifaceted interactions involving all structural regions of the protein. SSB, ssDNA and SSB-interacting molecules are highly concentrated within the condensates, whereas phase separation is overall regulated by the stoichiometry of SSB and ssDNA. Together with recent results on subcellular SSB localization patterns, our results point to a conserved mechanism by which bacterial cells store a pool of SSB and SSB-interacting proteins. Dynamic phase separation enables rapid mobilization of this protein pool to protect exposed ssDNA and repair genomic loci affected by DNA damage.

## INTRODUCTION

Single-stranded DNA-binding proteins (SSB) are essential, ubiquitous and abundant proteins that are involved in all aspects of genome maintenance in all kingdoms of life (Shereda et al., 2008). Prokaryotes possess a structurally well conserved SSB protein which forms nucleoprotein filaments with single-stranded (ss) DNA to protect it from reannealing and harmful chemical alterations (Antony and Lohman, 2019; Shereda et al., 2008). In addition, SSBs recruit essential DNA metabolic enzymes to sites of action via protein-protein interactions (Shereda et al., 2008). In *E. coli* and most bacteria, SSBs function as homotetramers, but dimeric SSBs are also known (Shereda et al., 2008). Regardless of quaternary structure, functional units of SSBs contain 4 OB folds that assemble into a conserved 3D structure. Importantly, the vast majority of bacteria encode at least one SSB variant in which the OB fold is followed by a “tail” region which can be subdivided to the intrinsically disordered linker (IDL) and C-terminal peptide regions (CTP, C-terminal nine amino acids) (**Fig. 1A**) (Bianco, 2017; Raghunathan et al., 1997; Savvides et al., 2004). The IDL of SSB is one of the few known protein regions with a universally conserved propensity for structural disorder (Pancsa and Tompa, 2016); however, its length and amino acid composition are not conserved (Bianco, 2017; Paradžik et al., 2017). In contrast, the sequence of the CTP is highly conserved among bacteria and it mediates essential interactions with at least 18 proteins involved in nucleic acid metabolism (Antony and Lohman, 2019; Shereda et al., 2008). Accordingly, CTP mutations weakening protein-protein interactions are lethal (Antony et al., 2013). The CTP of *E. coli* SSB is also known to interact with the OB fold, possibly within the tetramer or between adjacent SSB tetramers (Kozlov et al., 2010; Shishmarev et al., 2014; Su et al., 2014).

**Figure 1.**
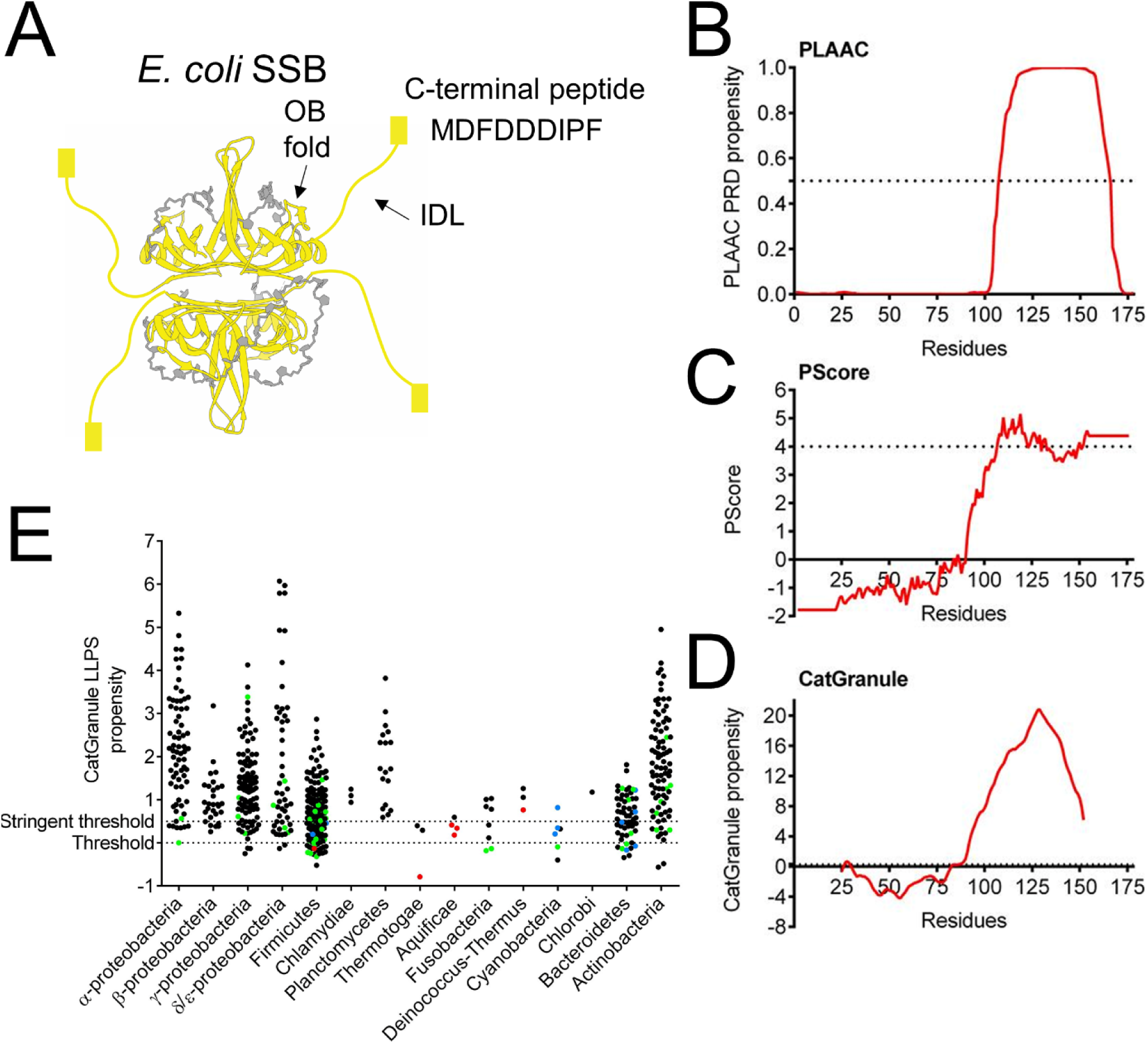
Bioinformatics analysis reveals conserved role of the IDL region of bacterial SSB proteins in phase separation. **A**, Crystal structure of the *E. coli* SSB homotetramer (yellow) bound to two 35mer ssDNA molecules (grey) (PDB ID: 1EYG). Each SSB monomer comprises an ssDNA-binding OB fold (oligonucleotide/oligosaccharide binding domain), an intrinsically disordered linker region (IDL; yellow line) and the conserved C-terminal peptide (CTP; yellow box; last 9 residues indicated). **B-D**, The **B**, PLAAC (prion-like amino acid composition; PRD: prion-like domain), **C**, PScore and **D**, CatGranule algorithms indicate prion-like features (PLAAC) and a propensity for liquid-liquid phase separation (LLPS) for the C-terminal region of *E. coli* SSB (residues 113-177). Predicted LLPS propensities are shown as red lines, whereas the threshold values of the methods are indicated as black dashed lines on each graph. For each method, the residues with predicted values above the corresponding threshold value are the ones predicted with a positive LLPS propensity. For the CatGranule method (**panel C**) the threshold value is at *y* = 0; the dashed line is slightly shifted to enhance its visibility. Note that CatGranule does not provide predicted values for protein termini, resulting in its prediction curve being shorter than the SSB sequence. The algorithms used are validated predictors of LLPS propensity (Vernon and Forman-Kay, 2019). **E**, CatGranule-predicted LLPS propensity scores for SSB proteins of representative bacterial strains from 15 major phylogenetic groups of Bacteria indicate broad conservation of the LLPS propensity of SSB across the bacterial kingdom. CatGranule provides a profile and a single propensity value for each protein, having a basic minimum threshold of 0.0 and a more stringent threshold of 0.5 for LLPS propensity (see Methods). For Thermotogae, Aquificae and Cyanobacteria, less than 50 % of SSBs scored positively (0 %, 25 % and 16.7 %, respectively), while in the other 12 groups this fraction was above 50 %. Overall, LLPS propensity is highly conserved (502 positive hits of 717 SSBs using the stringent threshold). We identified three possible reasons for the low scores of certain SSBs. (*i*) Thermotogae and Aquificae abound in hyperthermophilic species, wherein structurally disordered regions are generally heavily shortened and affected by adaptive sequence composition changes (Pancsa et al., 2019b). To assess if this could be the reason for their low predicted LLPS propensity, we identified hyperthermophilic species (optimum growth temperature ≥ 75°C based on the Genomes Online (Mukherjee et al., 2017) and/or BacDive databases (Reimer et al., 2019)) in the whole dataset (respective SSBs highlighted in red). (*ii*) We also identified SSB sequences that appeared to be fragments based on the absence of the CTP motif (and often the IDL too) but could not be excluded due to the lack of orthologs (highlighted in blue). (*iii*) As we do not expect all SSB variants of a strain to drive LLPS, SSBs having an ortholog with higher predicted LLPS propensity retained in the clean dataset were identified and highlighted in green. As expected, the highlighted SSBs tend to have low scores. Cases (*i*-*iii*) together explain ∼35 % of CatGranule predictions below the basic threshold, and ∼21.4 % of predictions below the stringent threshold.

In the past few years, many eukaryotic and viral proteins were shown to form dynamic liquid condensates termed as membrane-less organelles that fulfil crucial functions in normal cellular physiology as well as stress tolerance (Alberti et al., 2019; Banani et al., 2017; Shin and Brangwynne, 2017). Such condensates include, among others, the nucleolus, stress granules, P-bodies, neuronal granules, post-synaptic densities, the heterochromatin and super enhancers (Banani et al., 2017). These entities form via liquid-liquid phase separation (LLPS) that is usually driven by multivalent weak protein-protein or protein-nucleic acid interactions, mostly mediated by intrinsically disordered regions (IDRs) of proteins (Shin and Brangwynne, 2017). The proteins driving phase separation and their specific interaction partners are highly concentrated in the resulting condensates, while the contents of the condensates can also undergo dynamic exchange with the surroundings through the phase boundary (Banani et al., 2016; Shin and Brangwynne, 2017). Such phase-separated protein condensates provide diverse functional advantages for cells. For instance, they can serve as activators of reactions due to the high local concentrations of components, as biomolecular filters due to their selectivity, as stress sensors due to their responsiveness and as reservoirs due to their ability to store macromolecules in an inactive, condensed state (Alberti et al., 2019; Pancsa et al., 2019a). Accordingly, mutations affecting the phase separation propensity of proteins are often linked to neurodegeneration and other pathological conditions (Alberti and Dormann, 2019). Whereas LLPS seems to be a heavily exploited process in eukaryotes and viruses (Heinrich et al., 2018; Nikolic et al., 2017, 2018; Zhou et al., 2019), up until today only a few bacterial proteins were reported to undergo LLPS. These include *Caulobacter crescentus* Ribonuclease E (RNase E) that orchestrates RNA degradosomes (Al-Husini et al., 2018), *Escherichia coli* (*E. coli*) FtsZ that forms the division ring during cell divisions (Monterroso et al., 2019) and *Mycobacterium tuberculosis* Rv1747 that is a membrane transporter of cell wall biosynthesis intermediates (Heinkel et al., 2019). All in all, the hitherto accumulated observations imply that LLPS is a fundamental mechanism for the effective spatio-temporal organization of cellular space, with emerging but hardly explored functions in bacteria.

LLPS is typically driven by multivalent weak interactions, which ensure the dynamic, liquid-like properties of the resulting condensates. Diverse protein sequence compositional features, structural modules, overall architectural properties and modes of protein-protein interactions are known to favor multivalent weak interactions and thus to promote LLPS. Disordered regions of low sequence complexity, prion-like regions, repeated short linear motifs and their cognate binding domains, nucleic acid-binding domains and oligomerization domains can all be hallmarks of LLPS propensity based on the properties of the hitherto described cases (Mittag and Parker, 2018).

The modular architecture of SSB, its ability to homooligomerize and to interact with multiple binding partners, its weakly conserved but compositionally biased IDL region, its conserved short linear CTP motif and central integrator role in genome maintenance processes all prompted us to raise the hypothesis that SSB drives the formation of liquid condensates through LLPS. Here we show that SSB is indeed capable of LLPS among physiological conditions, mediated via multifaceted inter-tetramer interactions involving all SSB protein regions. SSB, ssDNA and SSB-interacting partners are highly concentrated within the phase-separated droplets, whereas LLPS is overall regulated by the stoichiometry of SSB and ssDNA. Our bioinformatics analysis indicates that the LLPS-forming propensity of the SSB IDL is broadly conserved across all major phylogenetic groups of Eubacteria. Together with the recently observed dynamic spot-like subcellular localization patterns of SSB (Zhao et al., 2019), our results suggest that bacterial cells store an abundant pool of SSB and SSB-interacting proteins in phase-separated condensates via a conserved mechanism. The discovered features enable rapid mobilization of SSB to ssDNA regions exposed upon DNA damage or metabolic processes to serve efficient repair, replication and recombination.

## RESULTS

### SSB forms dynamic LLPS condensates in physiologically relevant conditions in vitro

Due to the fact that SSB shares several features with hitherto described LLPS drivers, we sought to determine if LLPS could be a yet undiscovered capability of SSB facilitating its multifaceted roles in genome maintenance. We performed a bioinformatics analysis that revealed that *E. coli* SSB, in particular its IDL region, shows high propensity for LLPS, according to multiple dedicated sequence-based prediction methods (Bolognesi et al., 2016; Lancaster et al., 2014; Vernon et al., 2018) (**Fig. 1B-D**). This feature of SSB is universally conserved among bacteria, as the majority of analyzed SSBs (72.1 %) harbored by representative bacterial species from 15 large phylogenetic groups show similar LLPS propensities (**Tables S1-2; Fig. 1E**).

In line with its predicted LLPS propensity, purified *E. coli* SSB (30 µM; SSB concentrations are expressed as those of tetramer molecules throughout the paper) forms an opaque, turbid solution at low NaCl concentration (50 mM) *in vitro* at room temperature and also at the native temperature of *E. coli* cells (37°C) (**Fig. 2A-B**). The process is reversible as elevation of NaCl concentration to 200 mM leads to immediate loss of turbidity. The Cl^−^ concentration in *E. coli* cells depends on their environment (Schultz et al., 1962); however, while the exact value is not known, it can be assumed to be in a similar range (5-100 mM) as that measured for the cells of mammalian *E. coli* hosts, whereas the major metabolic anion in bacterial cells is glutamate (Glu; 100 mM) (Bennett et al., 2009). Strikingly, SSB forms a turbid solution even in 200 mM NaGlu (**Fig. 2A-B**). (It must be noted that experiments using NaGlu always contained a fixed amount of 20 mM NaCl, for technical reasons.) Investigation of the protein solution by differential interference contrast (DIC) microscopy after diluting SSB into low-salt buffer revealed the formation of regular spherical droplets with diameters in the µm range (**Fig. 2C**). Epifluorescence microscopic experiments obtained using a total internal reflection fluorescence microscope in which solutions of SSB mixed with a fluorescently-labeled SSB variant (SSB^G26C^, allowing for selective labeling with a cysteine-reactive fluorescein dye (Dillingham et al., 2008)) were visualized also support the presence of SSB droplets up to 100 mM NaCl or 500 mM NaGlu (**Fig. 2D**). Magnesium-acetate (MgOAc) had no apparent effect on droplet formation in the investigated concentration range (0 – 10 mM) (**Fig. S1A**). The apparent diameter of droplets shows a non-normal (and non-lognormal) distribution (**Fig. S1B-D**) with the median of the apparent droplet diameter slightly, but significantly, decreasing with increasing monovalent salt concentration (**Fig. 2E**), but remaining constant in the investigated MgOAc concentration regime (**Fig. 2F**). Turbidity measurements also confirmed that for NaCl a sharp cutoff concentration around 100 mM exists above which no droplets form, while NaGlu and MgOAc do not influence droplet formation (**Fig. 2G-H**). These observations indicate that SSB is able to undergo LLPS under physiological ionic conditions, even though Cl^−^ ions at higher concentrations inhibit the process, a property typical of LLPS systems (Alberti et al., 2019; Lin et al., 2018).

**Figure 2.**
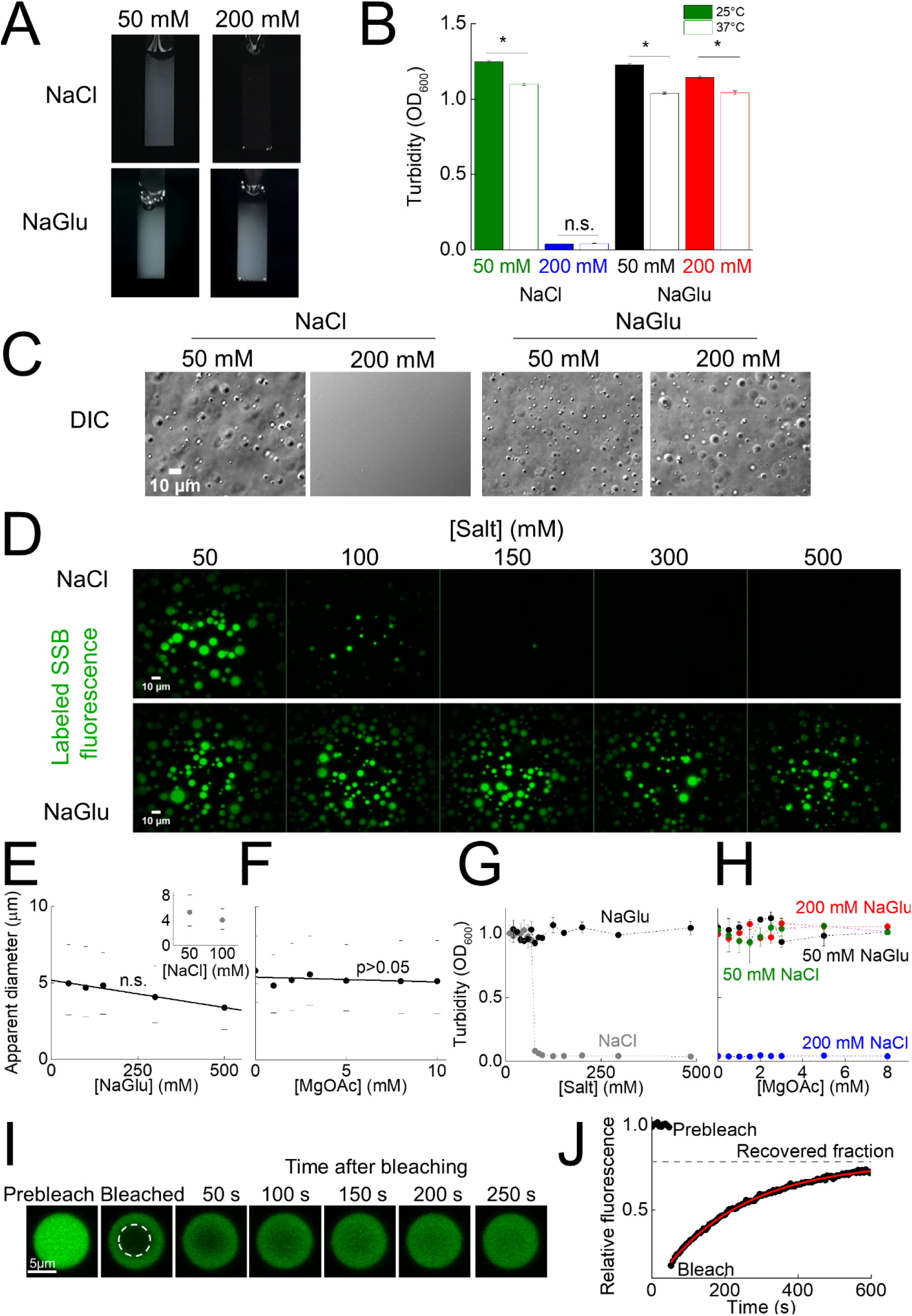
SSB forms phase-separated droplets in physiologically relevant ionic conditions in vitro. **A**, Photographs of cuvettes containing 30 µM SSB (SSB concentrations are expressed as of tetramer molecules throughout the paper) and the indicated salt concentrations (Glu: glutamate). Turbid appearance of samples indicates the presence of light scattering by particles of diameters larger than the wavelength of visible light. (We note that experiments with NaGlu contained a fixed amount of 20 mM NaCl for technical reasons.) **B**, Turbidity (measured *via* optical density at 600 nm (*OD*600)) of the samples at 25°C and 37°C. Turbidity (means ± SEM) decreased with increasing temperature (*: *p* < 0.01; n.s.: *p* = 0.076; n = 3, unpaired T-test). **C**, DIC (differential interference contrast) microscopic images of SSB samples shown in **panel A**. **D**, Fluorescence microscopic images obtained upon mixing 30 µM SSB and 0.3 µM fluorescein-labeled SSB at the indicated salt concentrations. (**Fig. S1A**) shows MgOAc dependence measurements. **E-F**, Salt (**E**, NaGlu, inset: NaCl; **F**, MgOAc) concentration dependence of the apparent droplet diameter (medians and 25/75 percentiles shown as bullets and dashes, respectively), determined from fluorescence microscopic experiments. Lines show linear fits (**E**, NaGlu: *p* < 0.05 indicates that the slope is significantly different from zero; **F**, MgOAc: slope not significantly different from zero (n.s., *p* > 0.05; one-way ANOVA analysis). Distributions are shown in **Fig. S1B-C**. **G-H**, Turbidity of 15 µM SSB samples at the indicated salt concentrations. **I**, Example images obtained during fluorescence recovery after photobleaching (FRAP) experiments in samples containing 30 µM SSB and 0.3 µM fluorescein-labeled SSB (bleached area diameter: 4,9 µm). **J**, After bleaching of fluorophores at the center of droplets (**Fig. S2G-H**), fluorescence intensity recovery (panel **I**) was followed in time. The time course of recovery was fitted by a single exponential decay function (solid line shows best fit). After bleaching, 68 ± 8% of the original fluorescence signal was recovered (recovered fraction) with a half-life (*t*_1/2_) of 175 ± 23 s.

During microscopic visualization, droplets sank toward the bottom of the sample chamber, and frequent fusion events were also observed (**Fig. S2A, Supplementary Video 1**) leading to an increase in apparent droplet diameter in time (**Fig. S2B-C**). This finding supports the liquid nature of SSB droplets. In addition, the turbidity of samples decreased at a rate that showed linear dependence on total SSB concentration, reflecting concentration dependence of droplet fusion frequency (**Fig. S2D-F**). Liquid behavior is also supported by fluorescence recovery after photobleaching (FRAP) experiments (**Fig. 2I**), where fluorescent molecules at the center of a droplet (**Fig. S2G-H**) were photobleached with high-intensity laser light in a confocal microscope and then the fluorescence recovery of the photobleached area was followed in time (**Fig. 2J**). Based on the sizes of the photobleached areas and the measured fluorescence recovery half-life values, the diffusion coefficient of labeled SSB was calculated to be 0.009 ± 0.001 µm^2^/s (**Table S3**). Based on the Einstein-Stokes equation, the theoretical diffusion coefficient of an SSB tetramer (with a Stokes radius of 3.8 nm (Meyer and Laine, 1990)) in water is ∼50 µm^2^/s. This indicates that SSB diffuses ∼5000 times slower in the droplets than it would in water, and thus the apparent dynamic viscosity within the droplets (5000 mPas) is in the range of that of honey (about 3.6 times that of glycerol).

### SSB drives LLPS at cellular concentrations

Next we investigated the effect of protein concentration on droplet formation (**Fig. 3A**) in the presence and absence of 150 mg/ml bovine serum albumin (BSA) used as a molecular crowder to simulate the cellular total protein concentration in *E. coli* (Kuznetsova et al., 2014). In both conditions, droplets were detected even at as low as 0.5 µM total SSB concentration; however, at low SSB concentrations the apparent diameters of SSB droplets were much smaller (**Fig. 3B, S3A-B**) and they rapidly diffused in the solution. The median of the apparent diameter of droplets was dependent on SSB concentration with a steep increase in the 3 to 4 µM concentration range and remained constant above 4 µM independently of the presence of BSA (**Fig. 3B**). SSB concentration-dependent turbidity measurements also confirmed phase separation by SSB in the 0.5 – 30 µM SSB concentration regime in all conditions except for 200 mM NaCl (**Fig. 3C**). There are approximately 2000 SSB tetramers present in *E. coli* cells (Schmidt et al., 2016), which results in about 5 µM intracellular SSB concentration, taking into account the average *E. coli* cell volume (0.65 µm^3^). Thus, our results suggest that SSB droplet formation can occur in cells even without the inducing effect of molecular crowding, with droplet sizes being limited by the amount of SSB within the cell. The findings that droplet frequency and size show a marked increase around the physiological SSB concentration (**Fig. 3A-B**) are suggestive of the possibility that the LLPS propensity of SSB has been evolutionarily shaped by physiological demands.

**Figure 3.**
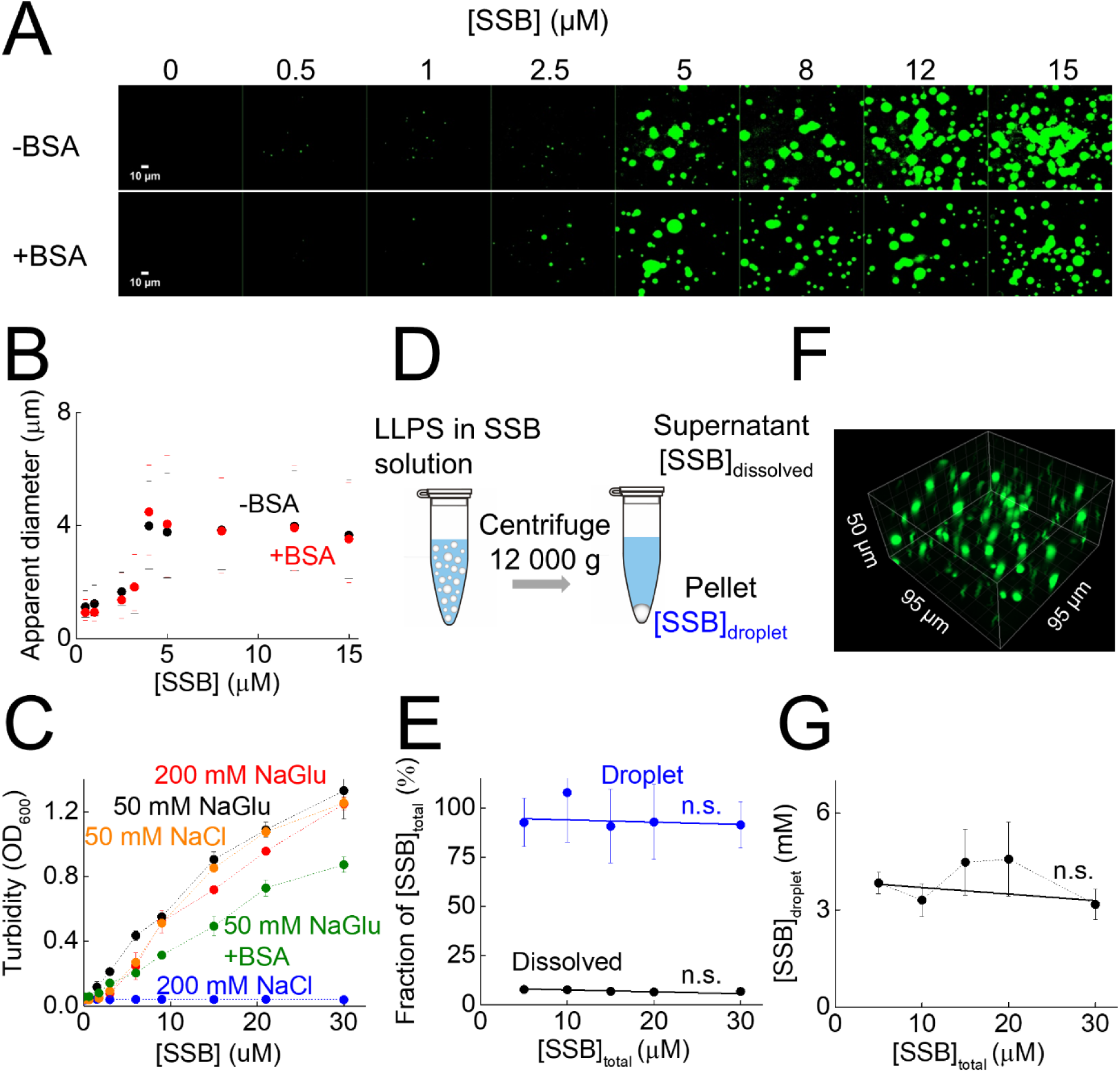
SSB forms highly dense droplets at cellular SSB concentrations in vitro. **A**, Fluorescence images of samples containing SSB at the indicated concentrations (including 0.3 µM fluorescein-labeled SSB) in the absence and presence of bovine serum albumin (BSA, 150 mg/ml). **B**, SSB concentration dependence of the apparent diameter of droplets (medians and 25/75 percentiles marked by bullets and dashes, respectively) in the absence and presence of BSA (150 mg/ml). **Fig. S3A-B** shows diameter distributions. **C**, SSB concentration dependence of turbidity in the presence of the indicated solution components. **D**, Schematics of centrifugation-based concentration determination experiments. **E**, Fraction of SSB in the dissolved (black) and droplet (blue) phases. Solid lines show linear fits. Slopes were not significantly different from zero (n.s., *p* > 0.05; one-way ANOVA analysis). **F**, Representative 3-D fluorescence image obtained in spinning disc microscopic experiments (15 µM SSB, 0.3 µM fluorescein-labeled SSB). **G**, Calculated concentration of SSB in droplets. Solid line shows linear best-fit. The slope was not significantly different from zero (n.s, *p* = 0.804, *n* = 3, one-way ANOVA analysis). **Fig. S4** shows control experiments revealing the effect of molecular crowders.

Measurements using various concentrations of bovine serum albumin (BSA) or polyethylene glycol (PEG_20000_; a 20-000-Da polymer that is routinely used as a synthetic molecular crowding agent) showed that droplet characteristics are influenced by different crowders (**Fig. S4**). Whereas BSA does not influence droplet size distribution, at high polyethylene glycol concentrations the mixing of SSB with labeled SSB is affected and thus non-homogeneous fluorescence distributions and lowered fluorescence intensities were observed in fluorescence microscopic experiments (**Fig. S4A-F**). On the contrary, in DIC experiments, BSA and PEG conditions were indistinguishable (**Fig. S4A-B**). Accordingly, both BSA and PEG altered the turbidity of solutions in a similar fashion (**Fig. S4G-H**) and reduced droplet fusion rates (**Fig. S4I-J**). PEG reduced the diffusion coefficient of labeled SSB in FRAP experiments by 3.9-fold (**Fig. S4K-M**), in line with the altered mixing properties observed in fluorescence experiments (**Fig. S4F**).

To calculate the concentration of SSB in the droplets, we separated the condensed fractions of samples of various SSB concentrations using centrifugation, and then determined the protein concentration from the supernatants and back-diluted pellets (resuspended in 1 M NaCl to starting volume) (**Fig. 3D**). The fraction of SSB molecules localized to the condensed phase remained constant over the investigated SSB concentration regime (5-30 µM) with an average of 95 ± 3 % of SSBs located in the condensed phase (**Fig. 3E**). Next, we estimated the volume fractionation of droplets and surroundings using 3-D images reconstructed from z-stack images prepared using a spinning disc microscope for samples of the same total SSB concentration as those used in the centrifugation experiments (SSB was mixed with fluorescently-labeled SSB in these experiments) (**Fig. 3F**). Based on the total SSB concentration, the fraction of droplet volume within the sample, and the fraction of SSB molecules localized to the droplets, SSB concentration within droplets was calculated to be 3.9 ± 0.3 mM on average, independent of total SSB concentration (**Fig. 3G**). Based on the Stokes radius of SSB, the water content of SSB droplets is calculated to be around 46 ± 4 v/v%, which is markedly lower than that of the cytoplasm (∼70 v/v%).

### Multivalent interactions between SSB IDL regions are required for LLPS, whereas interactions between the CTP and the OB fold enhance LLPS propensity

Previous studies suggested that the IDLs of adjacent SSB molecules can interact with each other to enhance the cooperativity of ssDNA binding, while the IDL was also proposed to interact with the OB fold (Bianco et al., 2017; Kozlov et al., 2015, 2017). Moreover, the central role played by similar low-sequence-complexity, Gln/Gly/Pro-rich IDRs in previously investigated LLPS systems (Bakthavachalu et al., 2018; Mannen et al., 2016; Milovanovic et al., 2018; Sabari et al., 2018) as well as the predicted LLPS propensities suggest a role for the IDL in SSB LLPS (**Fig. 1B-D**). To test how the different structural modules of SSB contribute to LLPS, we purified an SSB variant lacking the C-terminal peptide (SSBdC, comprising amino acids (aa) 1-170) and a variant that also lacks the IDL (SSBdIDL, aa 1-113) (**Fig. 4A, Fig. S5**). Both constructs were previously shown to be able to bind ssDNA (Kozlov et al., 2015). Strikingly, in contrast to SSB, SSBdC only forms droplets in the presence of 150 mg/ml BSA, whereas SSBdIDL forms amorphous non-spherical aggregates even in the presence of BSA, as assessed by DIC microscopy (**Fig. 4B**). In fluorescence experiments, 30 µM SSBdC formed droplets with 0.3 µM fluorescently-labeled SSB both in the presence and absence of BSA (**Fig. 4C**), whereas the formation of SSBdC (30 µM) droplets with 0.3 µM fluorescently-labeled SSBdC was only observed in the presence of BSA (**Fig. 4D**). However, 0.3 µM fluorescently-labeled SSBdC formed droplets with 30 µM SSB even in the absence of BSA (**Fig. 4D**). SSBdIDL was unable to form droplets in any investigated condition, and the aggregated structures had a faint appearance (**Fig. 4B-D**). Interestingly, mixing of 30 µM SSBdC with fluorescently-labeled 79mer homo-deoxythymidine (dT) ssDNA (Cy3-dT_79_) resulted in droplet formation both in the presence and absence of BSA, whereas experiments using SSBdIDL showed amorphous aggregates (**Fig. 4E**). Turbidity measurements also support that, in the absence of ssDNA, SSBdC only forms droplets in the presence of BSA (**Fig. 4F**).

**Figure 4.**
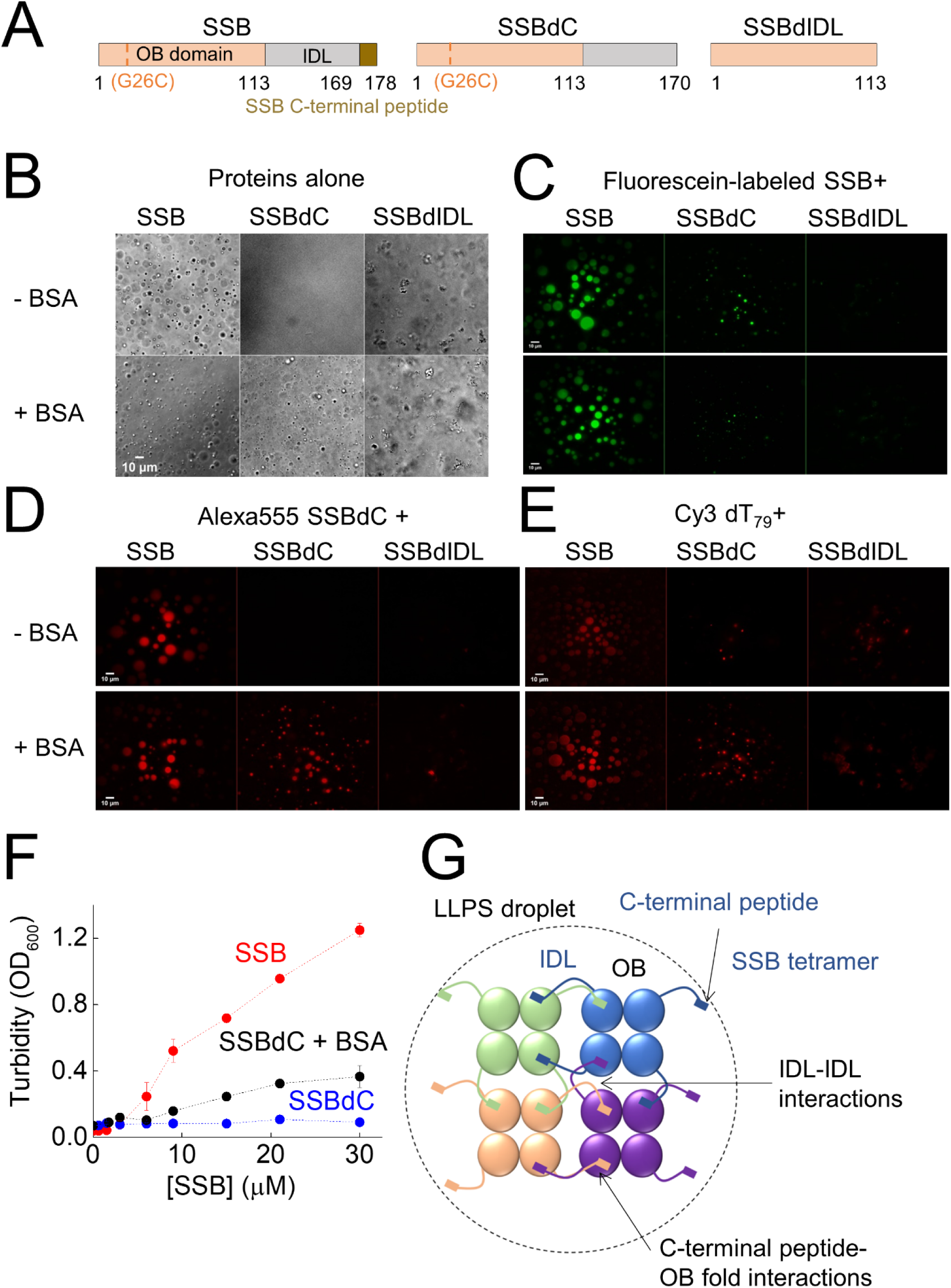
Multifaceted interactions of SSB structural regions are required for efficient LLPS. **A**, Schematic domain structure of SSB constructs (see also **Fig. 1A**). Numbers indicate amino acid positions at boundaries of structural regions. SSBdC lacks the CTP region, whereas SSBdIDL lacks the IDL and CTP regions. For site-specific fluorescent labeling, SSB variants harboring the G26C substitution were used (**Fig. S5**) (Dillingham et al., 2008). In addition to SSB constructs we also generated chimeric proteins of enhanced green fluorescent protein and the SSB C-terminal regions to test the role of the IDL and CTP regions in the absence of the OB fold (**Fig. S6**). **B**, DIC images of 30 µM SSB constructs in the presence and absence of 150 mg/ml BSA. **C-E**, Fluorescence images of 30 µM unlabeled SSB constructs mixed with 0.3 µM (**C**) fluorescein-labeled SSB, (**D**) 0.3 µM Alexa555-labeled SSBdC or (**E**) 0.3 µM cyanine 3 (Cy3) labeled dT_79_ ssDNA (79-nucleotide long homopolymer deoxythymidine) in the presence and absence of 150 mg/ml BSA. **F**, SSBdC concentration dependence of sample turbidity in the presence and absence of 150 mg/ml BSA (in 50 mM NaGlu). SSB data from **Fig. 3C** are shown for comparison. **G**, Model of LLPS-driving inter-tetramer SSB interactions based on the structure of SSB, the SSB-CTP interaction and the revealed roles of structural regions in LLPS.

To test the role of the SSB OB fold in LLPS, we generated two chimeric proteins. We fused the IDL plus CTP regions of SSB C-terminally to monomeric enhanced green fluorescent protein (eGFP) (**Fig. S6A-B**). We also generated an eGFP fusion variant wherein a short flexible (Gly-Gly-Ser)_4_ linker connected the CTP to eGFP instead of the IDL (**Fig. S6A-B**). Neither of these constructs (including eGFP alone) was able to preferentially partition into SSB droplets regardless the presence of BSA (**Fig. S6C-E**). However, when using PEG, the eGFP fusion proteins, but not eGFP alone, were able to enrich in SSB droplets with similar efficiency (**Fig. S6C-E**) and showed similar diffusion characteristics to those of SSB in the absence of PEG (**Fig. S6F-H**). This finding indicates that under PEG-induced crowding conditions the interaction between the eGFP-fused CTP and SSB’s OB fold enables the partitioning of eGFP into phase-separated SSB droplets, with the IDL having no apparent effect. Taken together, the above results highlight that all structural modules of SSB contribute to efficient phase separation. The tetramerization of OB folds brings about multivalency and enhances the local concentration of IDL regions, whereas weak IDL-IDL (and/or IDL-OB fold) interactions drive LLPS and bring about the liquid nature of the condensates (similarly to the IDRs of eukaryotic LLPS driver proteins), while the specific CTP-OB fold interaction enhances this LLPS propensity (**Fig. 4G**).

### ssDNA binding to SSB inhibits phase separation due to competition between ssDNA and CTP binding to the OB fold

A major cellular function of SSB is to cover ssDNA. Accordingly, we observed that fluorescently-labeled ssDNA (Cy3-dT_79_) is enriched in SSB droplets compared to the surroundings (**Fig. 4B, Fig. 5A**). We also found that ssDNA can diffuse within the droplets 2.3 ± 0.5 times faster than SSB, possibly due to its rapid dynamic interaction with adjacent SSB tetramers (**Fig. S7A-C**) (Kozlov and Lohman, 2002). Intriguingly, a labeled ssRNA molecule behaved similarly under these conditions (**Fig. 5A**). SSB was earlier shown to bind ssRNA (Meyer and Laine, 1990; Molineux et al., 1975). Although the interaction of SSB with ssRNA is weaker than that with ssDNA (**Fig. S7D-E**), this interaction is sufficient to drive RNA enrichment in SSB droplets due to the high concentration of SSB in this phase (**Fig. 3G**, **Fig. 5A**), raising the possibility of a yet undiscovered role for SSB in RNA metabolism.

**Figure 5.**
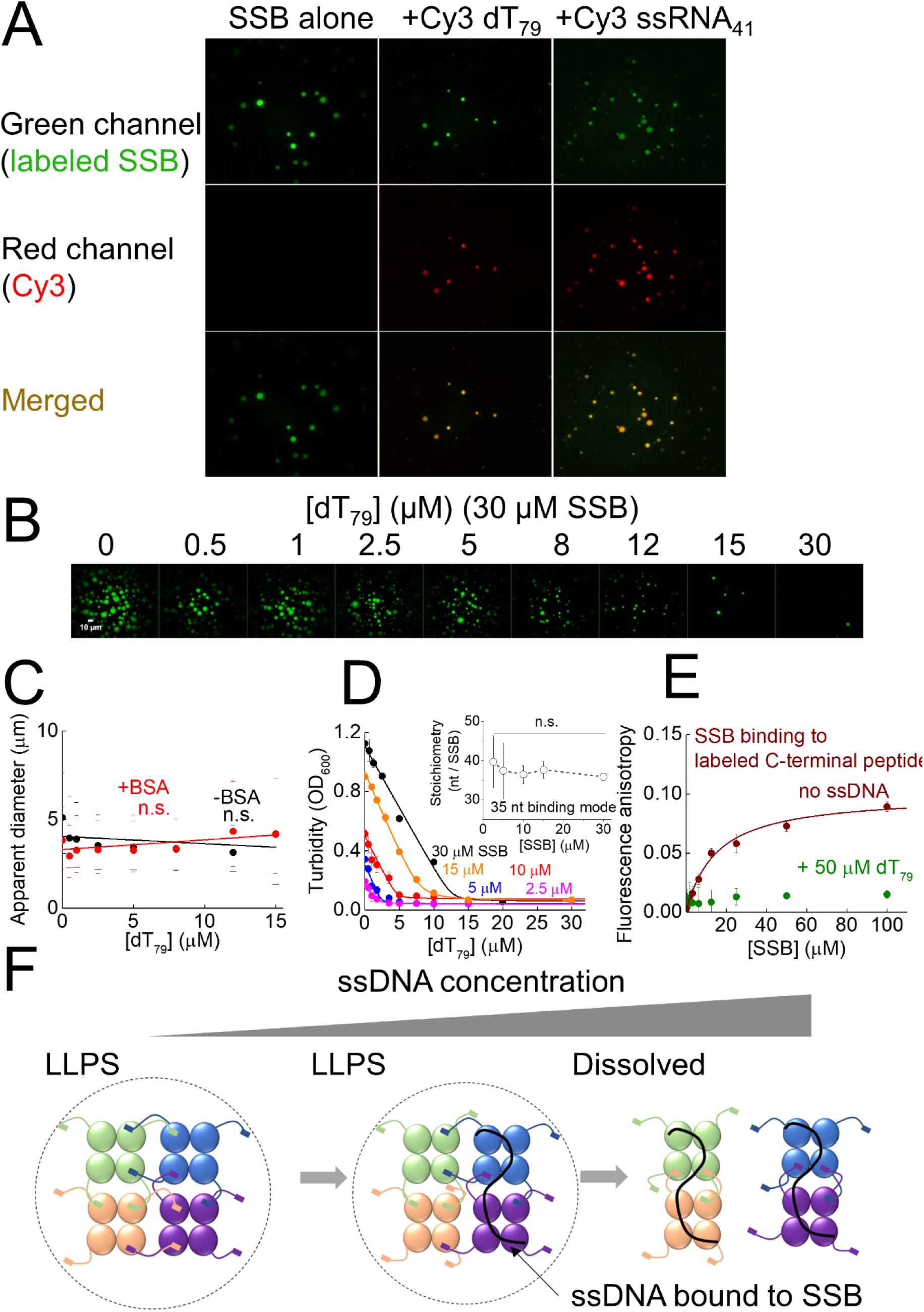
ssDNA regulates SSB phase separation by competing with the SSB C-terminal peptide for OB fold binding. **A**, Fluorescence microscopic images of 30 µM SSB, 0.3 µM fluorescein-labeled SSB (green channel) in the presence and absence of 0.1 µM Cy3-labeled dT_79_ or ssRNA (41 nt, non-homopolymeric) (red channel). **Fig. S7A-E** shows diffusion characteristics of Cy3-dT_79_ in droplets and binding characteristics of SSB to Cy3-dT_79_ and Cy3-ssRNA. **B**, ssDNA dependence of SSB droplet formation, as observed in fluorescence microscopic images of samples containing 30 µM SSB, 0.3 µM fluorescein-labeled SSB and unlabeled dT_79_ at the indicated concentrations. (**Fig. S7F**) shows additional data recorded in the presence of molecular crowding agents (BSA and polyethylene glycol (PEG20000)), using longer ssDNA (poly-dT with average length between 600-1000 nt), and controls for ssDNA dependence measurements. **C**, dT_79_ concentration dependence of apparent droplet diameters (medians and 25/75 percentiles marked by bullets and dashes, respectively) in the presence and absence of 150 mg/ml BSA. See **Fig. S7H-K** for additional experiments and distributions of apparent diameters. Lines show linear best-fits. One-way ANOVA analysis indicates that the slopes are not significantly different from zero (n.s., *p* > 0.05). **D**, ssDNA concentration dependence of turbidity of samples containing SSB at the indicated concentrations. Solid lines show fits using a quadratic binding equation (**Equation 2**, see Methods; best-fit parameters are shown in **Table S4**). Inset: apparent binding site size of SSB at saturating ssDNA concentrations (means ± SEM). Values at different SSB concentrations are not significantly different (n.s., *p* = 0.979, *n* =3, one-way ANOVA). **Fig. S8** shows additional ssDNA concentration-dependent turbidity measurements at higher NaGlu concentration, in the presence of molecular crowders or using poly-dT. **E**, Fluorescence anisotropy-based experiments monitoring SSB binding to 15 nM fluorescein-labeled, isolated SSB C-terminal peptide in the presence and absence of 50 µM dT_79_. Solid line shows best fit based on the Hill equation (**Equation 1**, see Methods). Fit results are shown in **Table S5**. **F**, Schematic model for the LLPS-inhibiting effect of ssDNA (black line). SSB tetramers and structural regions are shown as in **Fig. 4G**.

In the above experiments, substoichiometric amounts of ssDNA (or ssRNA) over SSB DNA binding sites were used. Therefore, we also examined the effect of higher DNA concentrations on SSB-driven LLPS. Strikingly, droplets disappeared with increasing concentrations of unlabeled dT_79_ (**Fig. 5B, Supplementary Video 2-3**) or poly-dT (homo-deoxythymidine ssDNA with average length 600-1000 nt), regardless of the presence of BSA or PEG (**Fig. S7F-G**). The apparent diameter of droplets was not influenced by the presence of ssDNA either in the absence or the presence of BSA (**Fig. 5C**, **Fig. S7H-K**). However, the number of detected droplets decreased with increasing ssDNA concentration in all studied conditions (**Fig. 5B, Fig. S7F-G**) along with the measured turbidity of the samples (**Fig. 5D, Fig. S8**). Strikingly, in the presence of PEG, a shallower ssDNA dependence of turbidity was detected compared to other conditions, indicating a decreased sensitivity of droplet formation to ssDNA (**Fig. S8D**).

SSB can bind ssDNA in two major binding modes that can reversibly interconvert (Antony and Lohman, 2019; Lohman and Ferrari, 1994). Therefore, we tested how biding modes affect droplet formation. At low ionic strengths and high SSB:ssDNA concentration ratios, the binding mode equilibrium is shifted towards the so-called 35-nt mode in which on average two monomers of the SSB tetramer bind ssDNA with an apparent binding site size of 35 nt (Antony and Lohman, 2019; Lohman and Ferrari, 1994). High ionic strengths and low SSB:ssDNA concentration ratios favor the 65-nt binding mode in which ssDNA is wrapped around all four SSB tetramers (Antony and Lohman, 2019; Lohman and Ferrari, 1994). The analysis of ssDNA-dependent turbidity curves revealed that in all conditions, except for PEG, turbidity linearly decreases with ssDNA concentration (as expected for a stoichiometric binding reaction; **Fig. 5D, Fig. S8A-C, Table S4**) until the point at which all SSB molecules become ssDNA-bound and therefore droplet formation is completely inhibited, regardless of the binding mode (**Fig. 5D, Fig. S8**). Previous studies using nuclear magnetic resonance (NMR) spectroscopy showed that the CTP binds to the same region on the OB fold as does ssDNA (Shishmarev et al., 2014) and it was suggested that the IDL, and especially the CTP, influence ssDNA binding (Antony and Lohman, 2019; Bianco, 2017; Kozlov et al., 2010). In line with these observations, we found that when SSB is bound to ssDNA, the binding of the isolated CTP peptide to SSB was significantly weaker than that for DNA-free SSB (**Fig. 5E**). This observation, together with the LLPS-promoting effect of the CTP (**Fig. 4, S6**), suggests that droplet formation is regulated by the competition between the CTP and ssDNA for the OB fold (**Fig. 5F**).

### Even weak interaction partners of SSB enrich in phase-separated SSB condensates

The above results demonstrated that specific interactions with SSB allow enrichment of interactors in SSB droplets. Thus, we tested how the interaction strength and the molecular size of interactors influence their ability to partition into phase-separated SSB condensates (**Fig. 6A-B, Fig. S9A**). *E. coli* RecQ helicase, a 67-kDa protein that was shown to interact with SSB dominantly via the CTP (Mills et al., 2017; Shereda et al., 2007), readily enriches in SSB droplets (**Fig. 6A,C,D**). Interestingly, RecQ point mutants with a markedly weakened affinity to the CTP (Shereda et al., 2009) also enrich in the droplets with similar efficiency (**Fig. 6A,C,D, Table S5**). In contrast, BLM helicase (a human RecQ homolog) and eGFP are both unable to enrich in SSB droplets (**Fig. 6A,C,D, Table S5**), highlighting that specific interactions with SSB are required for proteins to partition into SSB condensates. Probably due to the high SSB concentration within the droplets (**Fig. 3G**), even weak interactions (*K*_d_ in the range of tens of µM) appear to be sufficient for enrichment in the SSB condensates. In line with this proposition, the isolated SSB CTP peptide became enriched in the droplets, but a control peptide that does not interact with SSB was not enriched (**Fig. 6B-C,E, Table S5**). In addition, small molecules, such as deoxycytidine triphosphate (dCTP) or fluorescein, are slightly enriched in droplets, but not as effectively as SSB interactors, indicating that small molecules are able to enter the separated phase and the droplet environment has a slight concentrating effect (**Fig. 6B-C**). As expected for interactors, both RecQ and the isolated CTP peptide can diffuse within the droplets and their rate of diffusion is apparently dictated by their size (**Fig. S9B-D**).

**Figure 6.**
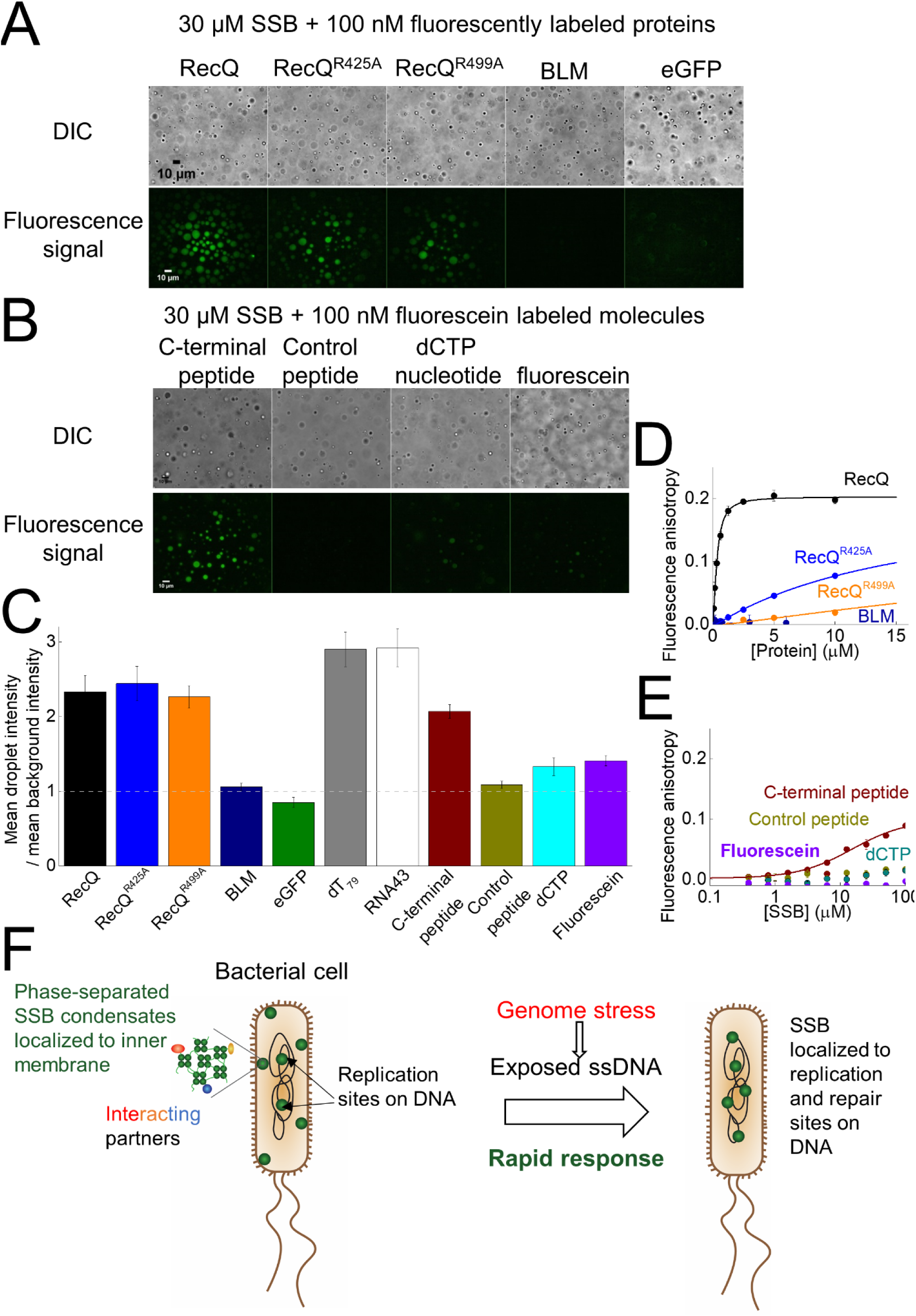
Specific interactions drive the enrichment of SSB-interacting partners in phase-separated SSB droplets. **A**, DIC and fluorescence microscopic images obtained upon mixing 30 µM SSB and 0.1 µM of various fluorescently-labeled proteins (Alexa488-labeled RecQ helicase (wild-type or variants harboring the R425A (RecQ^R425A^) or R499A (RecQ^R499A^) substitutions, fluorescein-labeled human BLM (Bloom’s syndrome) helicase or eGFP. See **Fig. S9** for additional data on protein constructs. **B**, DIC and fluorescence microscopic images obtained upon mixing 30 µM SSB and 0.1 µM fluorescein-labeled isolated SSB C-terminal peptide (CTP), fluorescein-labeled control peptide (12mer), fluorescein-labeled deoxycytidine triphosphate (dCTP) or fluorescein. **C**, Enrichment of various molecular components in SSB droplets, inferred from the ratio of the mean signal intensity within droplets and the mean background intensity, determined from background-uncorrected fluorescence images recorded for the indicated fluorescent molecules. **D-E**, Fluorescence anisotropy titrations of (**D**) 15 nM fluorescein-labeled isolated SSB CTP with unlabeled proteins used in **panel A** (except eGFP) and (**E**) titrations of 15 nM fluorescein-labeled CTP, control peptide, fluorescein labeled deoxycytidine triphosphate (dCTP) or fluorescein with unlabeled SSB. Solid lines show best fits based on the Hill equation (**Equation 1**). Best-fit parameters are shown in **Table S5**. **F**, Proposed model for the *in vivo* role of SSB LLPS, based on data presented here and earlier *in vivo* imaging results of Zhao *et al*. (Zhao et al., 2019).

### Bioinformatics analysis indicates conservation of LLPS propensity in human SSB homologs

Besides Replication Protein A, the long-known main eukaryotic ssDNA-binding protein, recent studies identified human SSB homologs hSSB1/SOSB1 and hSSB2/SOSB2 that are similar in size to bacterial SSBs and similarly possess C-terminal IDL regions (Ashton et al., 2013; Lawson et al., 2019). Using the bioinformatics tools applied in **Fig. 1B-D**, we assessed if the LLPS propensity of bacterial SSB proteins is conserved in these human homologs. The algorithms indicated similar degrees of LLPS propensity for the IDL regions of both human proteins to that for *E. coli* SSB, indicating a potential role for LLPS in eukaryotic DNA metabolic processes (**Fig. S10**).

## DISCUSSION

Taken together, here we show that *E. coli* SSB undergoes LLPS and forms viscous, liquid-state protein droplets under physiologically relevant ionic conditions and protein concentrations, both in the presence and absence of molecular crowders. Efficient phase separation requires all structural modules of SSB and is regulated by the specific interaction between the CTP and the OB fold as well as the stoichiometry of available SSB and ssDNA. Saturation of SSB binding sites by ssDNA, independent of SSB’s DNA-binding mode, prevents LLPS because ssDNA and the CTP compete for the same binding site on the OB folds. We also observed an LLPS inhibitory effect specifically for Cl^−^ ions, which were shown to be able to interact with the OB fold, likely through lysine side chains that were previously shown to interact with phosphate groups of ssDNA (Kozlov and Lohman, 2006). As the CTP interaction site of the OB fold is located at its ssDNA binding site, it is possible that Cl^−^ ions compete with the CTP for binding to the OB, as does ssDNA. In line with proposition, in NMR studies increasing NaCl concentrations inhibited the OB-fold – CTP interaction, whereas even 300 mM NaGlu had no effect on the interaction (Su et al., 2014). While other effects of Cl^−^ cannot be excluded, these results highlight the importance of the OB-fold – CTP interaction in LLPS.

Given that *E. coli* cells contain around 2,000 SSB tetramers and that we found the concentration of SSB to be steadily around 3 mM within the droplets, the theoretical maximum for intracellular SSB droplet size is ∼117 nm in diameter that can be achieved if the entire pool of cellular SSB molecules form a single droplet. Although particles of this size range can be visualized by superresolution microscopy, the compelling *in vivo* demonstration of LLPS by SSB is highly challenging. Strikingly, however, a recent study showed that in *E. coli* cells SSB localizes not only to DNA replication forks, but also forms multiple DNA-free protein foci near the inner cell membrane (Zhao et al., 2019). Membrane localization is explained by specific binding of SSB to lipid membrane components. Upon DNA damage, membrane-localized SSB foci were shown to disperse and, concomitantly, SSB re-localized to sites on the bacterial chromosome, likely to those containing exposed ssDNA (Zhao et al., 2019). Importantly, the ability of SSB to undergo ssDNA concentration-dependent LLPS, discovered in our current study, can explain the observed localization patterns. We propose that, when only a small amount of ssDNA is exposed, the majority of cellular SSB molecules (together with interacting partner proteins) will localize to membrane-proximal condensates; while an increase in exposed ssDNA sites (e.g. upon DNA damage) can trigger very rapid dispersion of condensates and relocalization of SSB to sites along the genome (**Fig. 6F**). Based on the diffusion coefficient of phase-separated SSB determined in our study (**Table S3**), the contents of an intracellular SSB droplet with a 100-nm diameter can be released with a half-time of 70 milliseconds even in the absence of additional signaling components.

Our current finding that even weak interaction partners of SSB are specifically enriched in the condensates is in line with previous findings showing that SSB interaction partners can form complexes with SSB even in the absence of ssDNA *in vivo* (Yu et al., 2016). Importantly, our findings highlight that the view that weak protein-protein interactions are functionally irrelevant needs to be revisited in the light of recently discovered properties of phase-separated protein condensates.

Based on the above findings, we propose that bacterial cells constitutively store a pool of SSB and SSB-interacting proteins in a condensed form at the inner membrane (**Fig. 6F**). This pool can be rapidly mobilized for rapid response to genomic stress and initiation of DNA repair. Interestingly, it was previously shown that bacterial cells expressing an IDL-deleted, but CTP-containing, SSB variant in place of wild-type SSB were viable in stress-free laboratory environment (Curth et al., 1996), likely due to the ability of SSB to efficiently bind to replication sites independent of its LLPS capability. However, another study showed that, when the IDL of *E. coli* SSB is removed or its composition is altered, the sensitivity of cells to UV damage is increased (Kozlov et al., 2015). The finding that the LLPS propensity of the IDL is highly conserved among bacteria (**Fig. 1E**) suggests that, while the ability of SSB to drive LLPS is not necessarily essential for cell viability under stress-free laboratory conditions, it confers important adaptive advantages in free-living bacteria, such as the adaptability to environmental stress and related DNA damage.

All in all, along with the few recently discovered cases of bacterial LLPS (Al-Husini et al., 2018; Heinkel et al., 2019; Monterroso et al., 2019), our findings highlight that LLPS is a fundamental, universally exploited mechanism that is governed by similar molecular mechanisms in bacteria and eukaryotes. The LLPS propensity predicted for human SSB homologs hSSB1/SOSB1 and hSSB2/SOSB2 suggests broad evolutionary conservation of this feature (**Fig. S10**), raising the possibility that SSB phase separation also plays a role in eukaryotic DNA metabolic processes.

## MATERIALS AND METHODS

### Reagents

All reagents and oligonucleotides were from Sigma-Aldrich unless otherwise stated. Fluorescent reagents were from Thermo Fisher Scientific. For concentration determination, *ε*_260_ values of 8400 M^−1^ cm^−1^nt^−1^ and 10300 M^−1^cm^−1^nt^−1^ were used for dT homopolymers and for non-homopolymeric oligonucleotides, respectively. DNA concentrations are expressed as those of oligo- or polynucleotide molecules (as opposed to those of constituent nt units) unless otherwise stated. SSB concentrations are expressed as those of tetramer molecules.

### General reaction conditions

Measurements were performed at 25°C unless otherwise indicated. Most measurements were carried out in LLPS buffer containing 20 mM Tris-acetate (Tris-OAc) pH 7.5, 5 mM MgOAc, 50 mM NaGlu, 20 mM NaCl unless otherwise indicated. Due to the NaCl content of SSB stock solutions, all NaGlu experiments contained a constant concentration of 20 mM NaCl. Experiments monitoring the effect of NaCl were performed in the absence of NaGlu.

### Bioinformatics analysis: datasets

Representative sets of bacterial reference proteomes at 15 % proteome similarity threshold were obtained from the PIR database (Protein Information Resource; release 2019 06; (Wu and Nebert, 2004)) for all 15 large bacterial phylogenetic groups defined in PIR (a total of 647 species). All proteins with a protein name exactly matching or including the phrase “Single-stranded DNA-binding protein” were selected (1004 sequences) and sequences labeled as “(Fragment)” were removed subsequently. While preserving at least one SSB protein for each species, for species with multiple SSB-like proteins, orthologs with missing/truncated structural modules (identified through visual screening of class-specific sequence alignments) were removed manually (275 excluded orthologs, 717 SSBs retained for analysis).

### Bioinformatics analysis: LLPS propensity prediction

We used the web servers of three dedicated prediction methods to obtain the LLPS propensity profile of *E. coli* and human SSBs. The PLAAC (Lancaster et al., 2014) method predicts prion-like domains based on sequence composition properties. It was used with parameters Core Length = 30 and α = 100. The PScore method (Vernon et al., 2018); default parameters used) estimates the ability of proteins to undergo LLPS solely based on their propensity for long-range planar pi-pi contacts. The CatGranule method (Bolognesi et al., 2016); default parameters used) is a machine learning approach that was trained on a set of granule-forming proteins and predicts LLPS propensity based on RNA-binding, structural disorder and amino acid composition features. Although these methods are preliminary and only take into account a restricted set of features indicative of LLPS, they — especially in combination — are highly useful for exploratory assessment of LLPS propensity (Vernon and Forman-Kay, 2019).

To predict the LLPS propensity of the clean set of 717 SSBs, the PLAAC and CatGranule web servers were used. Results were summarized for 15 major bacterial phylogenetic groups (**Table S1**) and for individual proteins (**Table S2**). Through visual screening of certain CatGranule propensity profiles that scored close to the original (basic) threshold value (0.0) we noticed that proteins with an overall CatGranule score between 0.0 and 0.5 often do not have the characteristic profile observed for *E. coli* SSB. The propensities were lower and sometimes dispersed throughout the sequence, and they did not necessarily have the characteristic maximum at the IDL. Therefore we decided to apply a more stringent threshold value of 0.5 to identify sequences with true LLPS propensity, resulting in 502/717 (70 %) positively assigned SSBs. PLAAC identified 28.5 % of SSBs with prion-like domains. 72.1 % of sequences scored positive with at least one of the methods. The fact that this proportion is only slightly higher than that of SSBs identified by CatGranule alone indicates that CatGranule could identify the subset of SSBs with prion-like features but it also identified LLPS candidates based on other features. Therefore we selected the CatGranule results for visualization (**Fig. 1D-E**).

### Cloning, protein expression and purification

Cloning and expression of SSB, SSB^G26C^ and SSBdC (comprising aa 1-170) is described in ref. (Mills et al., 2017). SSB^G26C^-dC was generated from SSBdC using the QuikChange (Agilent) mutagenesis kit. Mutagenesis was verified by DNA sequencing. SSB^G26C^-dC was purified similar to other SSB constructs with modifications as follows. In the case of SSB^G26C^-dC, only a HiTrap Heparin column (GE Healthcare) purification was applied. SSB^G26C^-dC did not bind to heparin column but appeared in the flow-through at high purity. SSBdIDL expression vector (encoding SSB aa 1-113) was generated from the vector encoding SSB using QuikChange mutagenesis. Purification of SSBdIDL was carried out similar to that for SSB, with changes as follows. During polyethyleneimine (PolyminP) fractionation, SSBdIDL remained in the supernatant instead of precipitating along with polyethyleneimine (as does SSB). The supernatant resulting from polyethyleneimine fractionation was purified applying the steps described for SSB.

The expression vector encoding full-length human BLM (aa 1-1417) was a generous gift from Dr. Lumir Krejci (Masaryk University, Brno, Czech Republic). BLM was expressed and purified based on ref. (Gyimesi et al., 2013). Cloning of RecQ is described in ref. (Sarlós et al., 2012). RecQ^R425A^ and RecQ^R499A^ expression plasmids were generated from that for wild-type RecQ using QuikChange mutagenesis. RecQ proteins were purified as in ref. (Harami et al., 2017).

EGFP-pBAD, allowing expression of N-terminally histidine-tagged eGFP, was a gift from Michael Davidson (Addgene plasmid # 54762). The coding sequence of eGFP was subcloned into a pTXB3 vector using the NheI and BsrGI restriction sites for IPTG-inducible protein expression. eGFP-IDL-CTP was cloned as follows. The coding sequence for histidine-tagged eGFP was amplified by PCR and cloned into the pTXB3 vector using the BsrGI and EcoRI restriction sites without a stop codon. The IDL-CTP encoding sequence of wild-type SSB (aa 118-178) was amplified by PCR using the wild-type SSB pET vector (Mills et al., 2017). The amplified region was inserted in-frame downstream of the histidine-tagged eGFP-coding region using the BsrGI and EcoRI restriction sites. In case of eGFP-(GGS)_4_-CTP (encoding four repeats of the GGS tripeptide and SSB’s C-terminal peptide motif (SSB aa 169-178)), both strands of the GGS_4_-CTP coding region were ordered as oligonucleotides that were designed to harbor the BsrGI and EcoRI restrictions sites. Oligonucleotides were annealed and ligated downstream of the histidine-tag-eGFP encoding sequence of the digested pTXB3 histidine-tag eGFP vector. Resulting constructs were verified by DNA sequencing.

eGFP constructs were transformed into *E. coli* KRX strain. Cells were grown in 1 L 2YT medium until reaching OD_600_ = 0.3. Protein expression was induced by addition of IPTG (1 mM) and rhamnose (0.2 w/v%). After overnight incubation at 18°C, cells were harvested and lysed in Lysis buffer (50 mM Tris-HCl pH 7.5, 0.5 M NaCl, 10 % v/v glycerol, 0.1 % v/v Igepal, 2 mM β-mercaptoethanol, 40 mM Imidazole, 1 mM PMSF (Phenylmethanesulfonyl fluoride) by sonication. After centrifugation of the lysate (JA-20 rotor, Beckman Coulter, 25000 g, 60 min, 4°C) the supernatant was loaded onto a Ni-NTA Agarose (Qiagen) column pre-equilibrated with Lysis buffer. After washing with 3 column volumes of Lysis buffer, the protein was eluted by the same buffer supplemented with imidazole (500 mM). Pooled protein fractions were dialyzed against Storage buffer (20 mM Tris-HCl pH 7.5, 200 mM NaGlu, 1 mM DTT, 10 % v/v glycerol) and concentrated using an Amicon Ultra 15-mL 10k (Merck Millipore) spin column. Protein concentrations were measured based on eGFP absorbance (*ε*_488_ = 56000 M^−1^ cm^−1^) and using *ε*_280_ eGFP-IDL-CTP = 29005 M^−1^ cm^−1^ and *ε*_280_ eGFP-GGS_4_-CTP = 23505 M^−1^ cm^−1^. Methods using different wavelengths yielded similar values that were averaged.

Purity was checked by SDS-PAGE for all proteins. Concentrations of purified proteins were measured using the Bradford method except for eGFP constructs (see above). Purified proteins were flash-frozen and stored in liquid N_2_ in 20-µL droplets.

### Fluorescent labeling of proteins

SSB variants harboring the G26C substitution were labeled as in ref. (Gyimesi et al., 2012) with the following exceptions. SSB^G26C^ was labeled with FITC (Fluorescein 5-isothiocyanate) instead of MDCC (*N*-[2-(1-maleimidyl) ethyl]-7-diethylaminocoumarin-3-carboxamide). SSB^G26C^-dC was labeled with AF555 (Alexa Fluor 555 C2 Maleimide). FITC was applied in 5-fold, AF555 in 2-fold molar excess over SSB monomers. Labeling reactions were performed for 5 h for FITC and 2 h for AF555 under argon gas. FITC labeling was stopped by the addition of Tris-HCl (to 50 mM final concentration). Maleimide reactions were stopped by the addition of 5 mM DTT. The labeled proteins were purified using a PD10 (GE Healthcare) gel filtration column, pre-equilibrated with SSB Storage buffer (50 mM Tris-HCl pH 7.5, 200 mM NaCl, 3 mM DTT and 20 % glycerol). Proteins were dialyzed against Storage buffer and repetitively filtrated using an Amicon Ultra 30K spin column to remove residual free dye.

All RecQ variants and BLM were labeled with AF488 (Alexa Fluor 488 carboxylic acid, succinimidyl ester). Proteins were dialyzed against labeling buffer containing 50 mM MES pH 6.5, 200 mM NaCl and 5 mM DTT. AF488 was applied in 3-fold molar excess over proteins. Labeling reactions were carried out for 8 h in the case of RecQ, and for 2 h in the case of BLM. Reactions were stopped by the addition of Tris-HCl (to 50 mM final concentration). Labeling buffers were exchanged by dialysis against Storage buffers (RecQ variants: 50 mM Tris-HCl pH 7.5, 200 mM NaCl, 1 mM DTT and 10 v/v% glycerol; BLM: 100 mM Tris-HCl pH 7.5, 500 mM NaCl, 1 mM DTT, 10 v/v% glycerol). Proteins were filtrated using an Amicon Ultra 50K spin column (Sigma-Aldrich) to remove residual free dye.

Protein concentrations were determined using the Bradford method. Dye concentrations were determined also by visible light spectrometry using *ε*_492_ = 78000 M^−1^ cm^−1^ for NHS-fluorescein and FITC, *ε*_555_ = 158000 M^−1^ cm^−1^ for AF555 and ε_488_ = 73000 M^−1^ cm^−1^ for AF488. The degrees of labeling were 25 ± 2.5 % for SSB^G26C^-FITC, 82 % for SSB^G26C^dC-AF555; 13 % for RecQ-AF488, 10 % for RecQ^R425A^ – AF488, 13 % for RecQ^R499A^ and 22 % for BLM-AF488.

Proteins were checked for purity and integrity using SDS-PAGE, frozen in liquid N_2_ in small aliquots and stored at −80 °C (SSB variants) or in liquid N_2_ (RecQ variants and BLM).

### Peptides

N-terminally fluorescein-labeled SSB C-terminal peptide (Flu-MDFDDDIPF), unlabeled C-terminal peptide (WMDFDDDIPF) and unlabeled control peptide (WDFMMDDPFID) were dissolved in 20 mM Tris-HCl pH 7.5, 10 mM ammonium bicarbonate, 500 mM NaCl. The N-terminally fluorescein-labeled control peptide (Flu-TRTKIDWNKILS) was dissolved in water.

### Fluorescence anisotropy titrations

15 nM of fluorescein-labeled molecules were titrated with increasing concentrations of SSB in LLPS buffer (50 mM Tris-OAc pH 7.5, 20 mM NaCl, 50 mM NaGlu and 5 mM MgOAc). Fluorescence polarization of 20-µl samples was measured in 384-well low-volume nontransparent microplates (Greiner Bio-one, PN:784900) using a Synergy *H4* Hybrid Multi-Mode Microplate Reader (BioTek) at 25°C and converted to anisotropy values.

To obtain dissociation constants from fluorescence anisotropy experiments, fits using the Hill equation were used. Here, the observed signal in binding experiments will be determined by

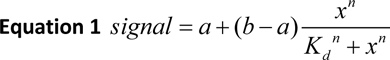

where *a* is the signal from the reporter in the absence of binding partner, *b* is the signal at saturation, *x* is the concentration of the binding partner, *K*_d_ is the dissociation constant of the interaction, and *n* is the Hill coefficient. *n* > 1 or *n* < 1 indicates positive or negative cooperativity, respectively.

To measure the effect of PEG_20000_ on ssDNA binding, we used a 3’-fluorescein-labeled oligonucleotide (TCCTTTTGATAAGAGGTCATTTTTGCGGATGGCTTAGAGCTTAATTGCGCAACG-Fluorescein).

### Turbidity measurements

Turbidity of 40-µl samples (light depletion at 600 nm wavelength) was measured in 384-well transparent microplates (Nunc Thermo Fisher PN:242757) in a Tecan Infinite Nano^+^ plate reader instrument using a 600 ± 20 nm band pass filter at 25°C or 37°C. The optical path length for the samples was 0.476 cm in all experiments. SSB or other buffer components have no light absorption at 600 nm based on absorption spectra. For phase-separated samples, light scattering by phase-separated droplets leads to apparent light absorption. Between reads, samples were mixed by gentle shaking using the orbital shaker of the plate reader instrument.

ssDNA concentration-dependent turbidity measurements were fitted using a quadratic binding equation derived from the law of mass action:

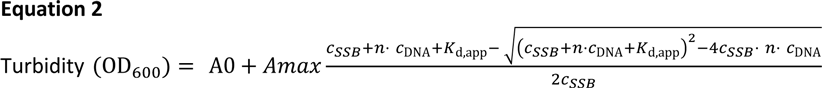

where *A0* is the turbidity of the sample at saturating ssDNA concentration, *Amax* is the turbidity in the absence of ssDNA, *c_SSB_* is enzyme concentration, *n* is the stoichiometry (mol SSB/mol ssDNA), *c*_DNA_ is the concentration of ssDNA, and *K*_d,app_ is the apparent dissociation constant of the SSB-DNA interaction. Binding site sizes were calculated by dividing the length (nt) of the ssDNA molecule with the determined stoichiometry value (mol protein/mol DNA molecule).

### Centrifugation-based concentration determination

500 µl SSB in various concentrations was incubated in LLPS buffer (20 mM Tris-OAc pH 7.5, 20 mM NaCl, 50 mM NaGlu and 5 mM MgOAc) at 25°C for 5 minutes. Samples were centrifuged using a tabletop Eppendorf centrifuge (13400 rpm (12000 g), 30 min). After centrifugation, a gel-like pellet was observable at the bottom of Eppendorf tubes. The supernatant was carefully decanted, and the pellet was resuspended in 500 µl buffer supplemented with 1 M NaCl as high NaCl concentration disperses the phase-separated SSB condensates. The absorbance of the supernatant and the resuspended pellet at 280 nm was measured in a quartz cuvette (1-cm pathlength) in a Beckman Coulter DU 7400 spectrophotometer. SSB concentration was determined spectrophotometrically using *ε*_280_ = 113000 M^−1^ cm^−1^ for the SSB tetramer.

### DIC microscopy

DIC images were captured using a Zeiss AxioImager M2 upright microscope (with objective Plan-Apochromat 20x 0.8 M27 and camera Axiocam 503 mono) equipped with an DIC filter and prism 0.55x. The Zen 2.3 (Zeiss) software was used to examine the obtained data. Raw, unprocessed DIC images are shown in the paper.

### Electrophoretic mobility shift assay (EMSA)

200 nM 5’-Cy3-labeled, 41-nt ssRNA (5’-Cy3-UAAGAGCAAGAUGUUCUAUAAAAGAUGUCCUAGCAAGGCAC-3’) or 41-nt ssDNA (5’ Cy3 – dT_41_) was titrated with increasing concentrations of SSB (0 – 500 nM) in LLPS buffer (20 mM Tris-OAc pH 7.5, 50 mM NaGlu, 5 mM MgOAc, 20 mM NaCl). Reactions were run on a 1 % w/v agarose gel at 120 V for 20 minutes. Fluorescent molecular species were detected upon excitation with a 532-nm laser in a Typhoon Trio+ Variable Mode Imager (Amersham Biosciences). Fits to the data were performed using the Hill and quadratic binding equations (**Equations 1-2**).

### Epifluorescence microscopy

A Nikon Eclipse Ti-E TIRF microscope with an apo TIRF 100x oil immersion objective (numerical aperture (NA) = 1.49) was used in epifluorescent mode. Samples were excited using a Cyan 488-nm laser (Coherent) or a 543-nm laser (25-LGP-193–230, Melles Griot). Fluorescence was guided through a dichroic mirror (ZT405/488/561/640rpc BS (dichroic cube)). Light was positioned into a Zyla sCMOS (ANDOR) camera, and images were captured using the NIS-Elements AR 4.50.00 imaging software. Measurements of 20-µl samples were carried out in µ-Slide Angiogenesis (Ibidi) microscope slides at 25°C. Before the measurements, chambers were blocked with 1 mg/ml Roche Blocking Reagent (Roche) in a buffer containing 20 mM Tris-OAc, pH 7.5 for 20 min. The blocking reagent was then removed and the wells were washed with the appropriate reaction buffer two times. Uniform experimental sets were obtained using the same optical setup (2×2 binning, 200-ms laser exposure). Sample components were mixed freshly before imaging and images were captured after 1 min following mixing.

### Image processing and particle analysis for epifluorescence experiments

Raw, unprocessed images from epifluorescence experiments were analyzed using the Fiji (ImageJ) software. For illustration purposes, images were background corrected as follows. Images from a given experiment were treated together as an image stack and the same processing was performed in parallel on all images in the stack. Brightness and contrast values were linearly adjusted using the automatic detection algorithm of Fiji. Images were background corrected using the built-in rolling ball background correction algorithm of Fiji (a rolling ball radius of 80 pixels was used).

For determination of the apparent diameter of fluorescent droplets, images were converted to 8-bit form and thresholded using the in-built image thresholder of Fiji (Huang method). The Fiji particle analyzer algorithm (smallest detected size was set to 0.2 µm^2^, circularity 0.2 – 1) was used to detect circular objects and calculate their areas (in turn, volumes). Detected areas were converted to apparent diameter values supposing that all fluorescent entities result from droplets. It must be noted that this method yields an apparent diameter for observable fluorescent circles, which may differ from the real diameter of the droplets due to the nature of epifluorescence imaging. Nevertheless, this method is suitable for analyzing changes in droplet size distribution under different conditions.

Raw images were used to determine the partitioning of the fluorescence signal between the droplets and their surroundings. As the fields were illuminated unevenly, only the droplets/circles present in the central area of acquired images (in a 40-µm circle around the image center) were analyzed. The mean intensity of a given circle was divided by the mean intensity of background adjacent to the measured object.

### Confocal fluorescence microscopy and Fluorescence Recovery After Photobleaching (FRAP) analysis

FRAP experiments were performed in Cellview cell culture slides with glass bottom (Greiner bio-one) containing 20 µl SSB solution in different conditions as indicated in the text. Images were acquired at 23°C with an inverted LSM800 confocal microscope (Zeiss) with a PlanApoChromat 63× oil immersion objective (NA = 1.4, Zeiss) using immersion oil (*n* = 1.518 at 23°C, Zeiss). Imaging parameters were 1.2-1.5× zoom, 1 Airy unit confocal pinhole and 500×500 pixels scanning resolution. For excitation of eGFP, Alexa 488 and fluorescein, a 488-nm diode laser was used. For excitation of Cy3 and A555, a 561-nm diode laser was used. Prebleach and postbleach imaging was carried out at 0.2-0.3 % laser power. Emission was detected at 400-650 nm (eGFP, Cy3, fluorescein) or at 450-700 nm (A555). Bleaching was performed in circular ROIs (*r* = ¼ *d* of the drops) in the middle of the drops using maximum laser power with 30 iterations to reach 80-90 % fluorescence reduction. Time-lapse images were collected in 5-s intervals. Pre- and postbleach periods were 10 and 90/120 frames, respectively. Before measurements, chambers were blocked with 1 mg/ml Roche Blocking Reagent (Roche) in a buffer containing 20 mM Tris-OAc, pH 7.5 for 20 min. The blocking reagent was then removed and wells were washed with the appropriate reaction buffer two times. Samples were allowed to settle to the bottom of the microscope slides during a 2-h incubation to prevent diffusion of droplets that could interfere with photobleaching. Confocal microscopic images were not background corrected.

Analysis of fluorescence signal recovery was performed using easyFRAP software (Rapsomaniki et al., 2012) with double normalization and single-exponential model (*I* = *I*_0_−*A* exp(−*t*/*τ*)) fitting using the raw images. Mobile fraction (mf = *A*/1-(*I*_0_-*A*)) and half-time of recovery (*t*_1/2_ = ln2/*τ*) values were calculated by the software. Diffusion coefficient (*D*) was calculated from *t*_1/2_ data using the equation *D* = 0.25 × *r*^2^/*t*_1/2_ (Kang et al., 2012), where *r* is the radius of the used bleaching ROI.

To calculate the diffusion coefficient of SSB and the dynamic viscosity of SSB droplets, the Einstein-Stokes equation was used: *D* = *kT*/(6π*ηr*) where *k* is the Boltzmann constant (1.38064852 × 10^−23^ m^2^ kg s^−2^ K^−1^), *T* is the absolute temperature, *η* is the dynamic viscosity (for water the latter is considered to be 0.001 kg/(ms) = 1 mPa·s) and *r* is the radius of the diffusing spherical object. For *r* in case of SSB, a Stokes radius of 3.8 x10^−9^ m was used (Meyer and Laine, 1990).

### Spinning disk microscopy and data analysis

For evaluation of droplet volumes, microscopic images were taken on a Zeiss Spinning Disk system equipped with Yogogawa SD and a 25x LD LCI Plan Apochromat 0.8 NA objective at 24°C. The imaged area was 285×285 µm in x-y axis and 50 µm in the z-axis direction. Z-stack images were acquired at 0.5-µm intervals using a 488-nm laser line and 100 ms exposure time. Cellview cell culture slides with glass bottom (Greiner bio-one) containing 150 µl SSB solution in different conditions (as indicated in the text) were used. Images from spinning disk experiments were not background corrected.

During image acquisition, droplets constantly sank toward the bottom of the imaging chamber. Thus, during capturing a z-stack, different cross sections of the droplets were visualized. To correct our measurements for the sinking effect, first we manually determined a fluorescence intensity threshold for detected objects and designated all voxels of an image above this threshold to be part of Surface objects by using Imaris software (Oxford instruments). Next, for each separate object we examined all images in the Z-stack (i.e. all recorded horizontal planes) and identified circles in the images by using a custom-written script for Imaris, utilizing the MatLab function imfindcircles. The largest circle for each object was selected, assuming that it represents the diameter of the sinking sphere/droplet. The volume of each droplet was thus estimated from the radius of the selected circles.

The concentration of SSB within droplets was calculated by multiplying the concentration of phase-separated SSB within the total sample volume (determined from centrifugation experiments) with the ratio of the total imaged volume and the cumulative volume of fluorescent spheres.

The water content of droplets was calculated by subtracting the product of the number of SSB particles in the droplet (derived from the intra-droplet SSB concentration, see above) and their Stokes volume from the total volume of the droplet.

### General data analysis

Mean ± SEM (standard error of mean) values are reported in the paper unless otherwise specified. For apparent droplet size distributions, mean ± SD (standard deviation) values and medians are shown. Sample sizes (*n*) are given for the number of observed particles in fluorescence microscopic experiments or the number of ensemble *in vitro* measurements performed (*n* = 3 unless otherwise specified). Data analysis was performed using OriginLab 8.0 (Microcal corp.). Pixel densitometry was performed using GelQuant Pro software (DNR Bio Imaging Ltd.). T-test and one-way ANOVA statistical analyses, together with the Tukey post-hoc test adjusted for multiple comparisons, were performed in OriginLab 8.0. Significance levels are indicated in the figure legends.

## ACKNOWLEDGMENTS

This work was supported by the Human Frontier Science Program (RGY0072/2010 to M.K.), the “Momentum” Program of the Hungarian Academy of Sciences (LP2011-006/2011 to M.K.), ELTE KMOP-4.2.1/B-10-2011-0002, NKFIH K-116072, NKFIH K-123989 and NKFIH ERC_HU 117680 grants to M.K. and NKFIH FK-128133 to R.P. G.M.H. and R.P. are supported by the Premium Postdoctoral Program of the Hungarian Academy of Sciences (PREMIUM-2017-17 to G.M.H. and PREMIUM-2017-48 to R.P.). The project was supported by the NRDIO (VEKOP-2.3.3-15-2016-00007 to ELTE) grant. We are grateful to Dr. Tibor Kovács for the help with DIC measurements and Gábor Szegvári for the custom-written script for volume estimation of droplets observed in spinning disc fluorescence microscopic experiments.

## AUTHOR CONTRIBUTIONS

GMH, RP, AMC and MK conceptualized the project. GMH designed the experiments. GMH, ZJK, JP, VB, RP and KT performed research. GMH, KT and RP analyzed data. GMH, KT, RP, AMC and MK wrote the paper.

## SUPPLEMENTARY INFORMATION

### SUPPLEMENTARY FIGURES

**Supplementary Figure 1.**
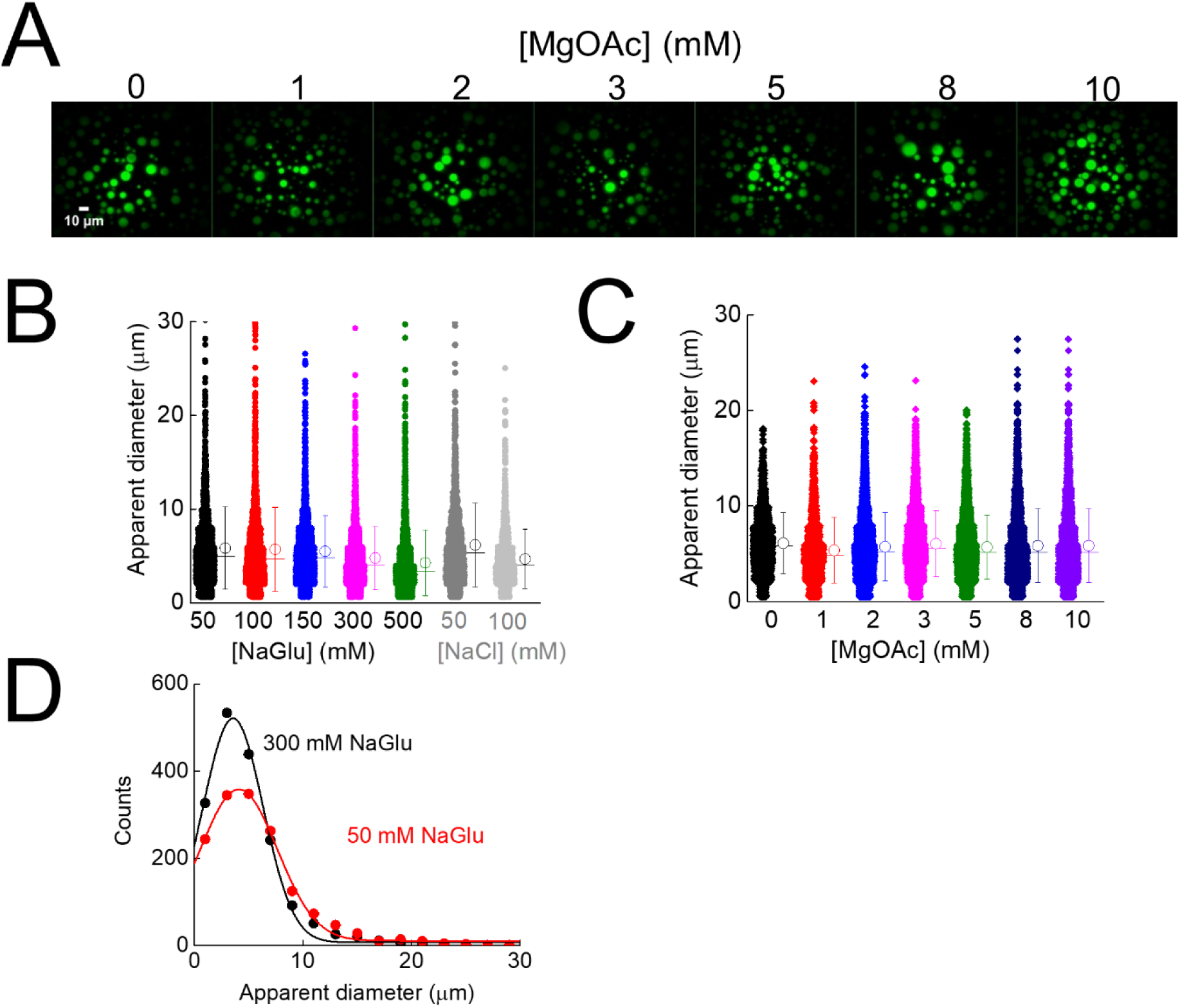
MgOAc dependence of droplet formation and distribution of apparent diameters of droplets. **A**, Fluorescence microscopic images recorded upon mixing 30 µM SSB and 0.3 µM fluorescein-labeled SSB in a buffer containing 20 mM Tris-OAc pH 7.5, 50 mM NaGlu, 20 mM NaCl and MgOAc at the indicated concentrations. **B-C**, Determined apparent diameter of droplets from NaGlu, NaCl (**Fig. 2D**) and MgOAc (panel **A**) concentration-dependent fluorescence microscopic experiments. Data were aggregated from multiple observation fields. Means (circles) ± SD and the median (horizontal line) are shown on the right side of raw data. **D**, Binned data can be fitted with a Gaussian function (solid lines); however, based on the Shapiro-Wilks normality test, both untransformed and log-transformed data significantly differ (*p* > 0.05) from a normal or log-normal distribution, due to the presence of a significant fraction of fluorescent droplets with large diameters (> 10 µm). For this reason, median values of apparent diameter distributions were used to detect concentration-dependent trends (**Fig. 2E-F;** in most cases, means were almost identical to the medians).

**Supplementary Figure 2.**
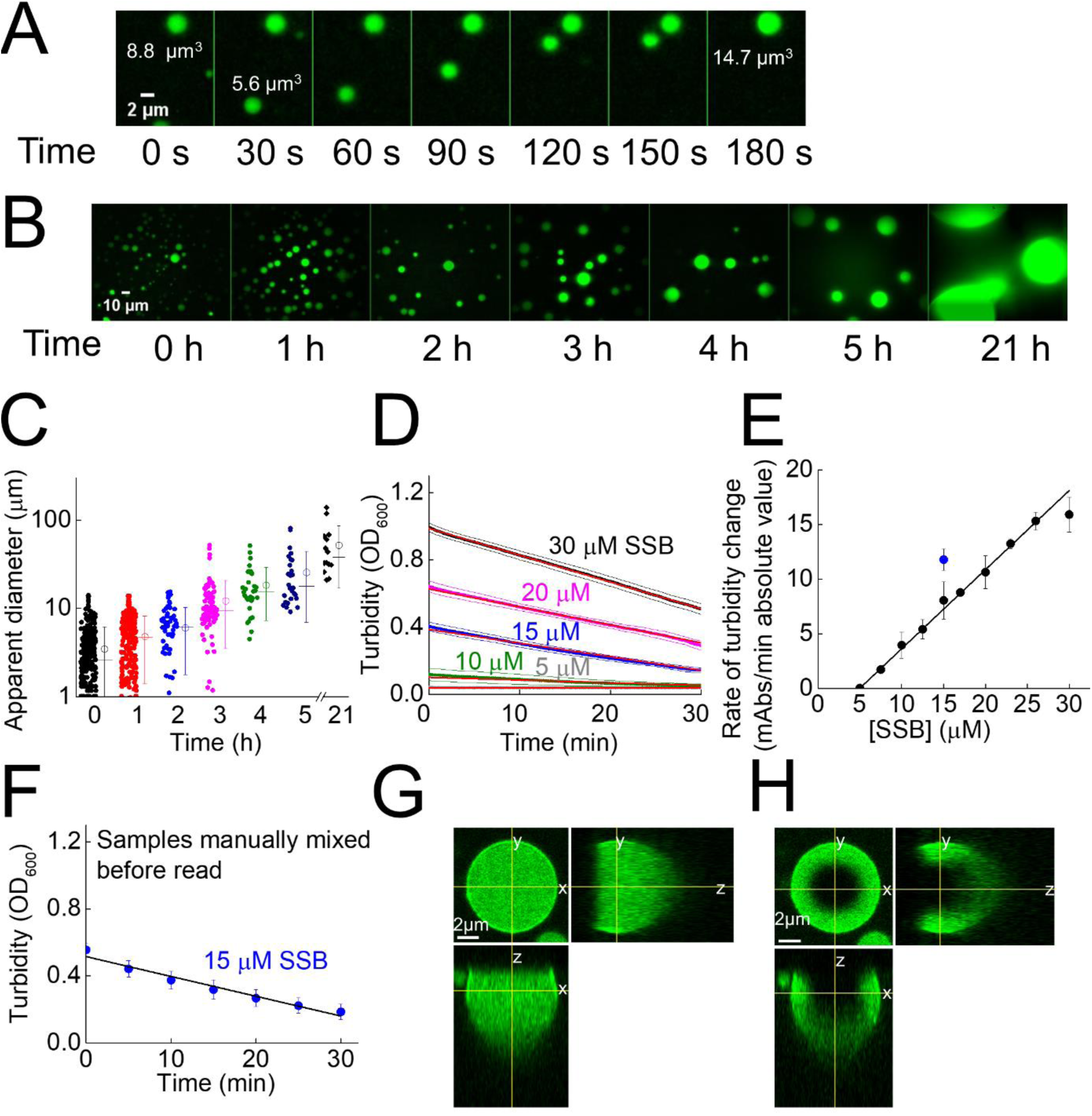
Liquid nature of SSB droplets is supported by observation of droplet fusion events and diffusion of SSB in droplets. **A**, Representative time series of fluorescence microscopic images obtained upon mixing 30 µM SSB and 0.3 µM fluorescein-labeled SSB in LLPS buffer (20 mM Tris-OAc pH 7.5, 50 mM NaGlu, 5 mM MgOAc, 20 mM NaCl) showing fusion of droplets. Fluorescence of samples was monitored in 5-s intervals. Droplet volumes were calculated based on apparent diameters. The volume of the fused droplet matched the sum of the fusing droplets within 2 % error. See **Supplementary Video 1** for a video showing droplet fusion. **B-C**, Representative time-lapse fluorescence images obtained upon mixing 30 µM SSB and 0.3 µM fluorescein-labeled SSB in LLPS buffer, showing progressive droplet fusion. Due to fusion, the number of observed droplets decreased whereas the apparent droplet diameter increased in time. **C**, Determined apparent diameter of droplets. Means (circles) ± SD and the median (horizontal line) are shown on the right side of raw data. **D**, Turbidity of unlabeled SSB samples (protein concentrations indicated) followed in time in a plate reader instrument in LLPS buffer (see Methods). Averaged traces with SD are shown (*n* = 3 independent experiments). Solid lines represent linear fits. **E**, The absolute value of the rate of turbidity change (panel **D**) increased linearly with increasing SSB concentration in the investigated SSB concentration regime (slope 0.72 ± 0.01 (mAbs min^−1^ µM^−1^), best-fit parameter and fitting error). The x-axis intercept of 4.9 ± 0.1 µM indicates that droplet formation and fusion become infrequent below an apparent critical SSB concentration of about 5 µM, in line with the abrupt change observed in droplet frequency and size in the same concentration regime in fluorescence imaging experiments (**Fig. 3A-B**). The time-dependent decrease in turbidity is likely to be caused by the decreasing number and increasing diameter of droplets due to fusion, in line with the results of fluorescence microscopic experiments (panel **A-B**). As the diameter of droplets is significantly larger than the wavelength of light used (600 nm), based on Fraunhofer diffraction theory, turbidity will decrease with increasing diameter (Raut and Kalonia, 2016) in addition to the effect resulting from the decreasing number of light scattering particles. The blue dot represents the result from experiments shown in panel **F**. **F**, To rule out that the observed turbidity decrease results from sinking of droplets toward the bottom of the microplate cell, we have performed similar time-dependent experiments in which the turbidity of 15 µM SSB was monitored in 5-min intervals with manual remixing of samples before each read. These measurements showed identical turbidity decrease kinetics (cf. panels **D-E**), confirming that turbidity changes result from droplet fusion. **G-H**, Example confocal microscopic images of SSB droplets settled to the bottom of the sample chamber in different cross-sections indicated by axis designations (30 µM SSB and 0.3 µM fluorescein-labeled SSB in LLPS buffer) (**G**) before and (**H**) after photobleaching. Droplets appear slightly oval due to moderate imaging distortion during Z-stack image capturing in the confocal microscope.

**Supplementary Figure 3.**
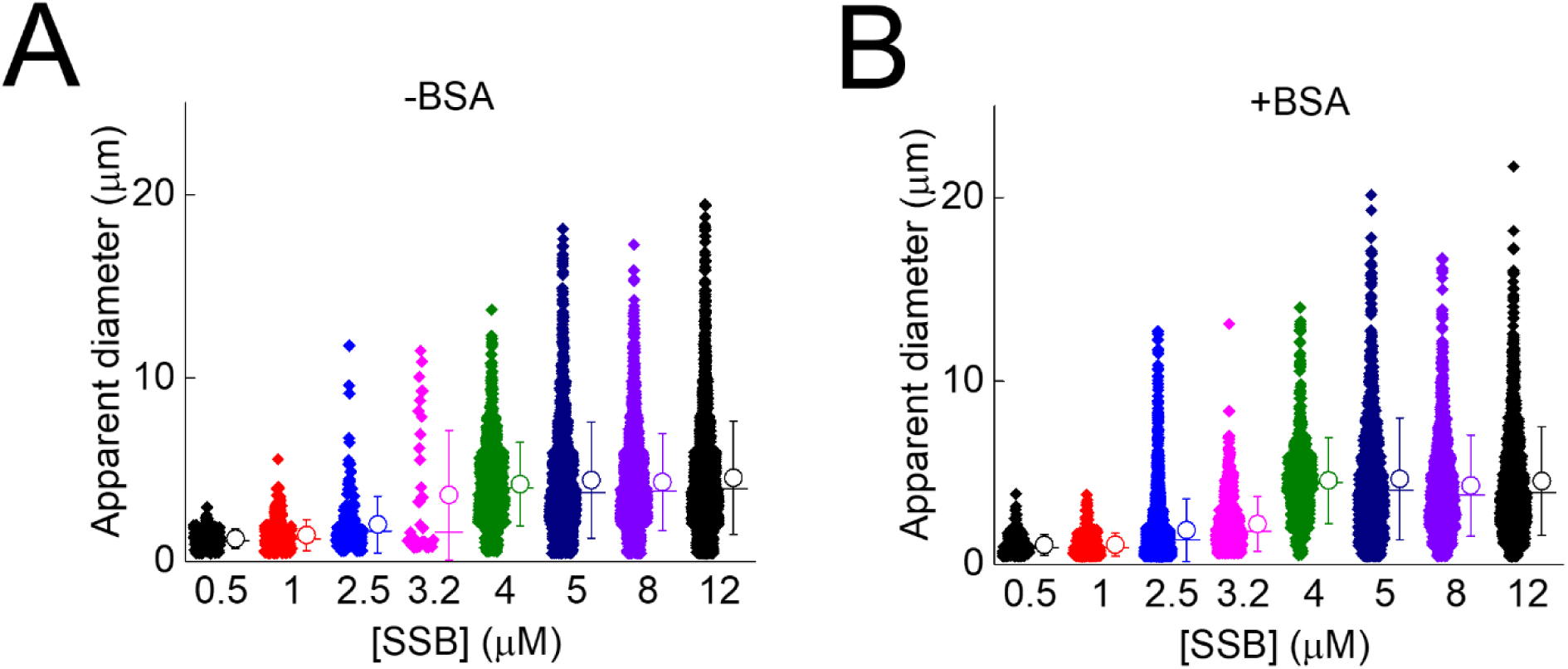
Apparent droplet diameters of samples with varying total SSB concentration. **A-B**, Determined apparent diameter of droplets from SSB concentration-dependent fluorescence microscopic experiments performed in the (**A**) absence or (**B**) presence of 150 mg/ml BSA in LLPS buffer (cf. **Fig. 3A**). Total SSB concentrations are shown. All samples contained 0.3 µM fluorescein-labeled SSB besides unlabeled SSB. Data were aggregated from two experiments, mean ± SD (circles and error bars) and median (horizontal lines) values are shown on the right side of raw data.

**Supplementary Figure 4.**
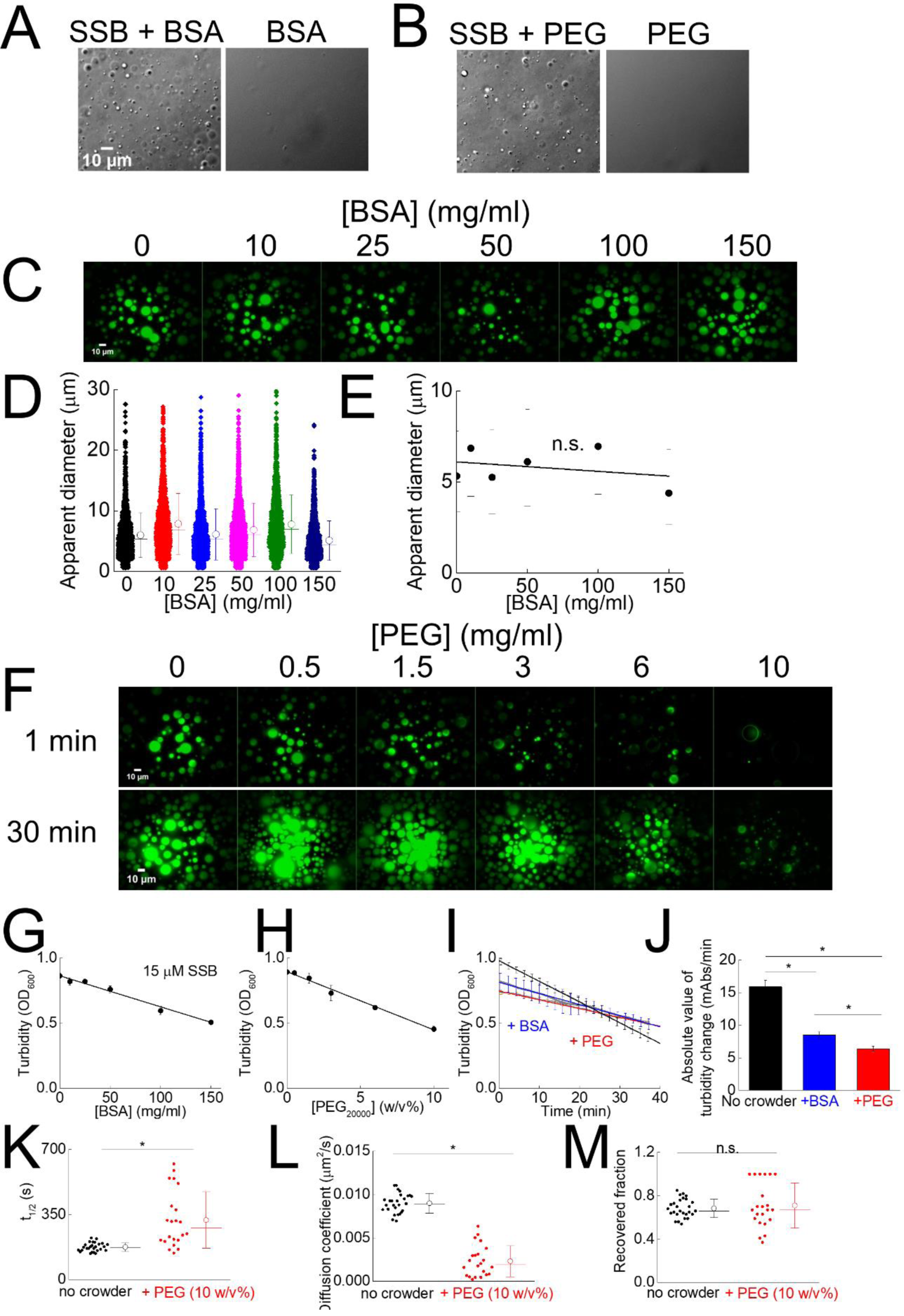
Effect of molecular crowders on LLPS by SSB. **A-B**, Example DIC images of 30 µM SSB in the presence of (**A**) 150 mg/ml BSA or (**B**) 10 w/v% PEG20000 along with the controls containing only BSA or PEG in LLPS buffer (20 mM Tris-OAc pH = 7.5, 5 mM MgOAc, 50 mM NaGlu, 20 mM NaCl). No objects were detected in the BSA and PEG controls. **C-E**, Effect of BSA concentration on LLPS by SSB. (**C**) Fluorescence images recorded for samples containing 30 µM SSB and 0.3 µM fluorescein-labeled SSB in LLPS buffer. (**D**) For diameter distributions, mean ± SD (circles) and median (horizontal lines) values are shown on the right side of raw data. (**E**) The median (bullets) of the apparent droplet diameter was independent of BSA concentration (slope of linear fit was not significantly different from zero (n.s., *p* > 0.05; one-way ANOVA analysis; 25/75 percentiles shown as dashes). **F**, Effect of PEG20000 concentration on SSB LLPS. Fluorescence images were obtained for samples containing 30 µM SSB plus 0.3 µM fluorescein-labeled SSB 1 min or 30 min (as indicated) after diluting SSB into the LLPS buffer. Interestingly, PEG slowed down the mixing of SSB with labeled SSB, and non-homogeneous fluorescence distributions in droplets were observed. The frequent appearance of a florescent halo around weakly fluorescent droplets is indicative of slow mixing. Moreover, after 30 min, more droplets with homogenous fluorescence distribution were observed. In contrast to fluorescence experiments, DIC images in the presence of 150 mg/ml BSA or 10 w/v% PEG were indistinguishable (panel **A-B**). Due to the above observations and the fact that after 30 min large droplets clustered together even in 0.5 w/v% PEG, we were unable to reliably determine the apparent diameter distribution of droplets. **G-H**, Both (**G**) BSA and (**H**) PEG20000 slightly decreased the turbidity of 15 µM SSB in LLPS buffer in a linearly concentration-dependent manner (mean ± SEM values shown, *n* = 3) the slopes (BSA: −0.0023 ± 0.0002 (mg/ml)^−1^; PEG: −0.044 ± 0.001 (mg/ml)^−1^) were significantly different from 0 (*p* < 0.05, one-way ANOVA). This effect of crowders on turbidity is striking and unexpected as the size distribution of droplets was similar in the presence and absence of BSA and the frequency of droplets also appeared similar (**Fig. 3A-B**, panel **C**). **I-J**, Strikingly, both 150 mg/ml BSA and 10 w/v% PEG20000 decreased the rate of turbidity change of 15 µM SSB in LLPS buffer. Solid lines show linear fits to time-dependent measurements. Averages of kinetic traces ± SD (for every 20^th^ datapoint) are shown. (**J**) BSA and PEG decreased the mean rate of turbidity change (mean ± SEM values shown, *n* = 3) by 1.9 ± 0.2 and 2.5 ± 0.2 fold, respectively. The effect of crowders on droplet fusion rate is possibly brought about by the increased dynamic viscosity of BSA and PEG solutions (150 mg/ml BSA ∼2.5 mPas, 10 w/v% PEG20000 ∼100 mPas). Nevertheless, the results indicate that molecular crowding and/or solution viscosity can influence droplet fusion dynamics. **K-M**, Effect of PEG20000 on SSB diffusion in droplets. 30 µM SSB and 0.3 µM fluorescein SSB was mixed in LLPS buffer containing 10 w/v% PEG. Droplets were allowed to settle to the bottom of microscopy chamber during a 2-h incubation. Droplets with uniform fluorescence distribution were photobleached and the half-life of fluorescence recovery (*t*_1/2_) and recovered fractions were determined as in **Fig. 2I**. (**K**) Diffusion coefficients calculated from the photobleached area and *t*_1/2_ (**K**). The presence of PEG20000 significantly decreased the diffusion rate of SSB by 4.5 ± 2.3 fold (*p* < 0.05, unpaired T-test). PEG-dependent changes in the diffusion rate may indicate that in the presence of PEG the dynamic viscosity of droplets increases or the increased viscosity of the droplet environment slows down SSB diffusion into the droplets. (**M**) The recovered fractions did not statistically differ for the two conditions (*p* > 0.05, unpaired T-test). The ∼70 % recovered fraction likely indicates that a portion of photobleached SSB remained in the droplet. Mean ± SD values for *t*_1/2_, diffusion coefficients and recovered fractions are shown in **Table S3**.

**Supplementary Figure 5.**
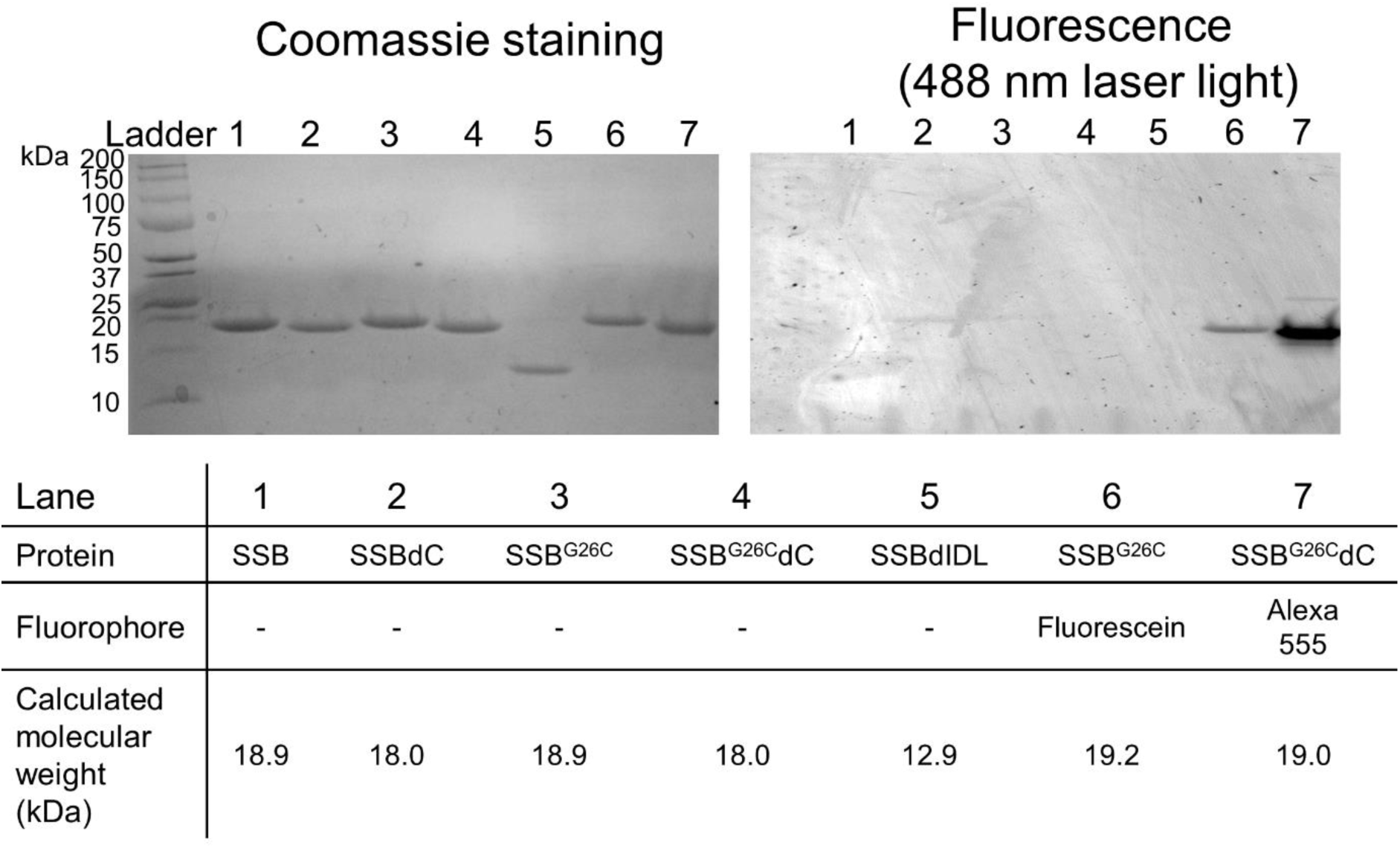
SDS-polyacrylamide gel electrophoretic analysis of unlabeled and labeled SSB constructs. Unlabeled SSB constructs (wild-type SSB, SSBdC lacking the last 8 amino acids of SSB, SSB^G26C^, SSB^G26^dC, and SSBdIDL lacking amino aa 113 onwards, fluorescein-labeled SSB^G26C^ and Alexa Fluor 555 (Alexa555) labeled SSB^G26C^dC (**Fig. 4A**; 5 µg each except for SSBdIDL, 3.5 µg) were separated on a custom-cast SDS-containing Tris-tricin polyacrylamide gel comprised of a stacking layer (4 w/v% acrylamide-bisacrylamide) and two separation layers (10 w/v% and 16 w/v% acrylamide-bisacrylamide). 15.5:1 and 32:1 acrylamide-bisacrylamide w/w ratios were used for the stacking and separation layers, respectively. Coomassie stained image is shown on the left. The same gel was scanned using a gel scanner instrument with a 546 nm laser light which excited both fluorescein and Alexa555 fluorophores covalently attached to SSBdC^G26C^ and SSBdC^G26C^dC, respectively. Precious Plus Protein Standard (Bio Rad) was used as molecular weight marker (Ladder). Calculated molecular weights for unlabeled and labeled proteins are shown in the table.

**Supplementary Figure 6.**
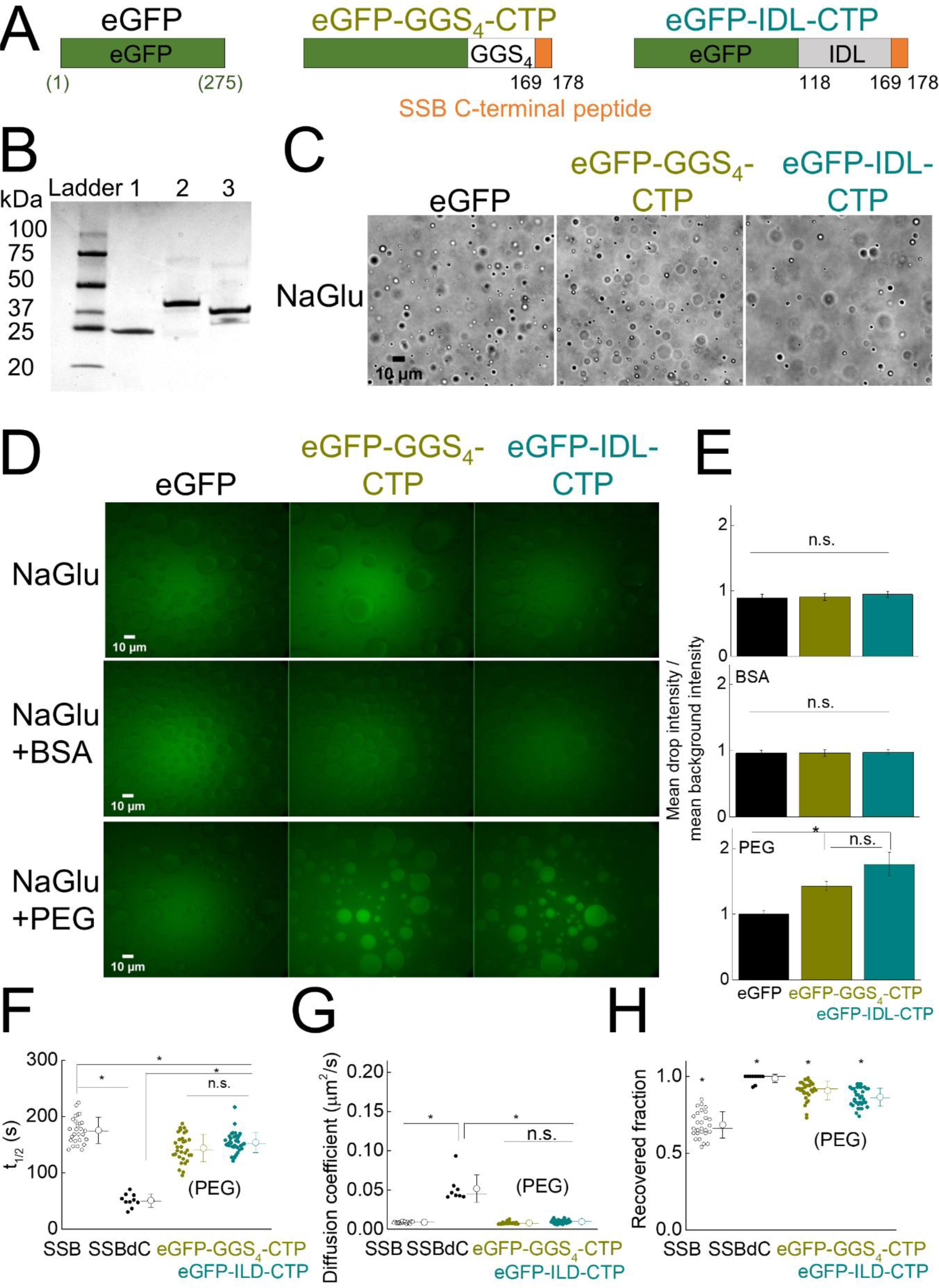
Chimeric eGFP-SSB constructs highlight the role of the OB fold in LLPS. **A**, Schematic domain structure of N-terminally His-tagged eGFP (His tag not shown) and chimeric eGFP proteins (aa residue numbers for eGFP (green) and those for the wild-type SSB sequence (black) are shown). In case of eGFP-GGS4-CTP, the last nine amino acids of SSB (MDFDDDIPF) are fused to the C-terminal of eGFP, following a fourfold repeated glycine-glycine-serine tripeptide sequence. In case of eGFP-IDL-CTP, the wild-type SSB sequence from aa position 118 onwards was fused to the C-terminal of eGFP. **B**, eGFP and eGFP chimeric constructs (3 µg each; cf. panel **A**) were separated using SDS-PAGE in a Mini-PROTEAN TGX 4-20% precast gel (Bio-Rad) and visualized by Coomassie staining. As a molecular weight marker, the Precious Plus Protein Standard (Bio Rad) was used. Calculated molecular weights are 27 kDa, 36.8 kDa and 32.7 kDa for eGFP (lane 1), eGFP-IDL-CTP (lane 2) and eGFP-GGS4-CTP (lane 3), respectively. **C**, Representative DIC images of 30 µM SSB in LLPS buffer in the presence of 0.1 µM eGFP constructs as indicated. In control experiments no objects were detected without SSB. **D**, Fluorescence images of 30 µM SSB in LLPS buffer in the presence of 0.1 µM eGFP constructs and in the presence or absence of 150 mg/ml BSA or 10 w/v% PEG20000. Images were not background corrected. **E**, Ratios of mean droplet intensity and the mean background intensity (mean ± SEM values are shown, *n* = 10) determined from fluorescent experiments (panel **E**). In the presence of PEG20000, both eGFP-GGS4-CT and eGFP-IDL-CTP were enriched in the droplets compared to eGFP (*p* < 0.05 for both constructs, one-way ANOVA). The enrichment values did not significantly differ between CTP-containing constructs, indicating that the CTP region alone (even in the absence of the IDL on eGFP) enables enrichment in SSB droplets. **F-H**, Diffusion of SSB and SSBdC (in the absence of PEG) compared to that of eGFP-IDL-CTP and eGFP-GGS4-CTP (in 10 w/v% PEG20000) in SSB droplets. 30 µM SSB and 0.1 µM fluorescein-labeled SSB or fluorescein-labeled SSBdC in the absence of PEG, or 30 µM SSB and 0.1 µM eGFP-IDL-CTP and eGFP-GGS4-CTP in the presence 10 w/v% PEG in LLPS buffer was mixed. Droplets were allowed to settle to the bottom of the microscopy chamber during a 2-h incubation. Droplets with uniform fluorescence distribution were photobleached and the half-life of fluorescence recovery (*t*_1/2_) and recovered fraction values were determined as in **Fig. 2J**. (**G**) Diffusion coefficients were calculated from the photobleached area and (**F**) *t*_1/2_. The diffusion coefficient of SSBdC was significantly higher compared to that of SSB and chimeric eGFP constructs, whereas the diffusion coefficients for the latter three constructs did not significantly differ (using a *p* threshold of 0.05, one-way ANOVA). (**H**) Intriguingly, the recovered fractions were higher for all constructs compared to those for SSB, and all recovered fractions were significantly different from each-other (*p* < 0.05, one-way ANOVA). Mean ± SD values for t_1/2_ values, diffusion coefficients and recovered fractions are shown in **Table S3**.

**Supplementary Figure 7.**
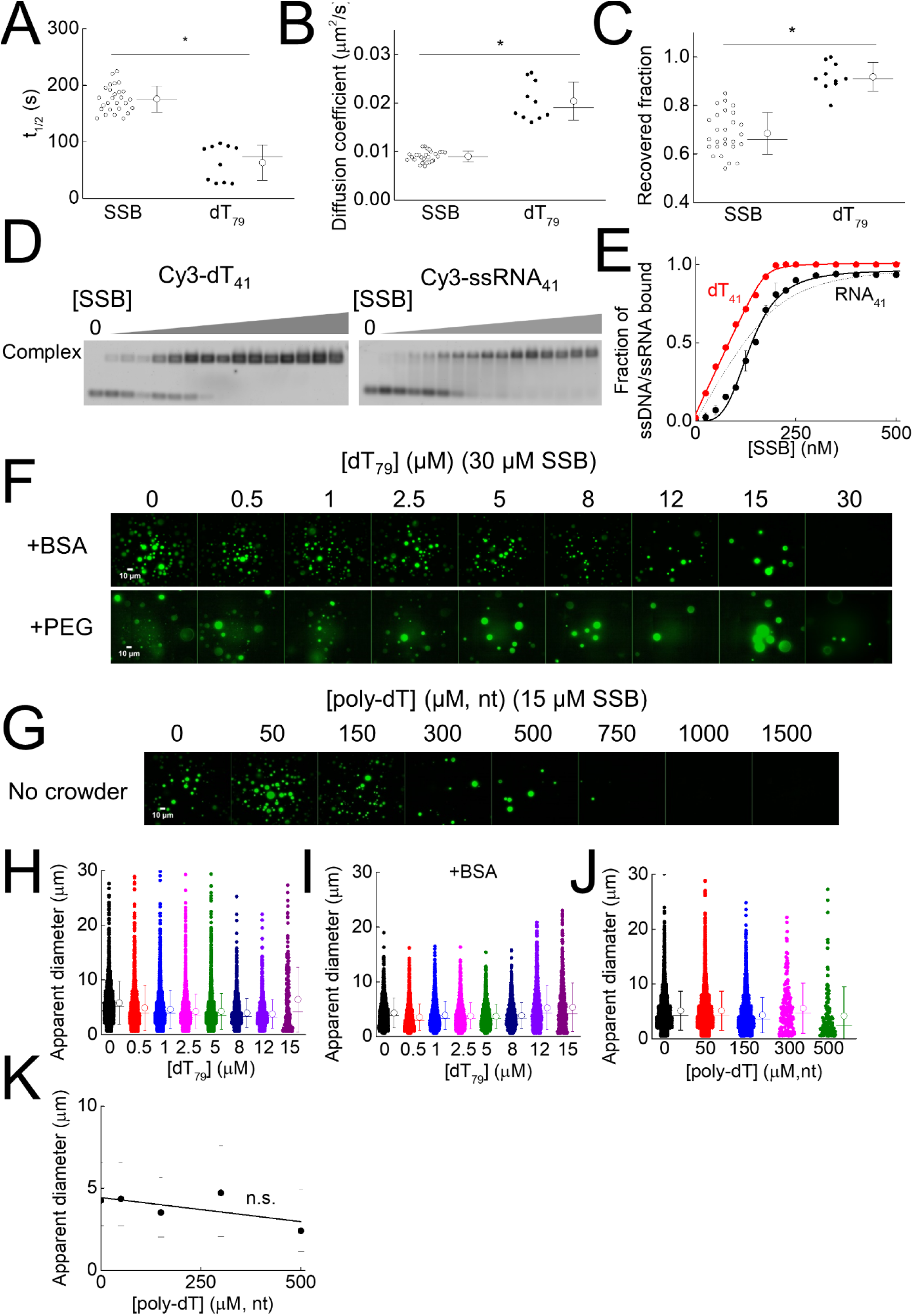
Additional data on the effect of oligonucleotides on LLPS by SSB. **A-C**, Diffusion of Cy3-labeled dT_79_ in SSB droplets. 30 µM SSB and 0.1 µM Cy3-labeled dT_79_ were mixed together in LLPS buffer. Droplets were allowed to settle to the bottom of the microscopy chamber during a 2-h incubation. Half-life values of fluorescence recovery (*t*_1/2_) and recovered fractions were determined as in **Fig. 2J**. (**B**) Diffusion coefficients were calculated from the photobleached area and (**A**) *t*_1/2_. The diffusion coefficient of dT_79_ was significantly higher compared to that of SSB (* *p* < 0.05, one-way ANOVA). (**C**) The recovered fraction for dT_79_ was also significantly higher compared to that of SSB (* *p* < 0.05, one-way ANOVA). Mean ± SD values for *t*_1/2_ values, diffusion coefficients and recovered fractions are shown in **Table S3**. **D**, Example electrophoretograms of electrophoretic mobility shift assays (EMSA) where a constant concentration of 0.2 µM Cy3-labeled dT_41_ or *ssRNA*_41_ was titrated with SSB (0 – 500 nM) in LLPS buffer. Molecular species were separated on a 1 w/v% agarose gel and visualized using a fluorescent gel scanner instrument. Upon SSB binding, the electrophoretic mobility of dT_41_ and *ssRNA*_41_ decreased. The single low-mobility band representing the complex indicates a 1:1 stoichiometry of both nucleic acid species with SSB. **E**, Fractions of labeled dT_41_ (red) or *ssRNA*_41_ (black) bound by SSB. The red solid and black segmented lines show fits using the quadratic binding (**Equation 2**) equation. The black solid line shows best fit of the Hill (**Equation 1**, see Methods) equation. SSB binds dT_41_ with high affinity (*K*_d_ = 2.3 ± 1.9 nM) and 1:1 stoichiometry. However, a sigmoidal binding isotherm was observed for ssRNA, reminiscent of a cooperative behavior. Fitting of the Hill equation revealed a dissociation constant of 161.7 ± 4.3 nM, and a Hill coefficient 4.7 ± 0.5. **F**, Representative images of fluorescence experiments with samples containing 30 µM SSB, 0.3 µM fluorescein-labeled SSB and the indicated concentrations of unlabeled dT_79_ in the presence of molecular crowding agents 150 mg/ml BSA and 10 w/v% PEG_20000_ in LLPS buffer. BSA had no apparent effect on droplet behavior. However, in the presence of PEG, droplets with a fluorescent halo and uneven fluorescence distribution were dominant (cf. **Fig. S4F**). The number of objects observed per field decreased with increasing ssDNA concentration. However, in contrast to BSA-containing or crowder-free conditions, numerous droplets were observed even for 30 µM dT_79_, suggesting that PEG renders the SSB droplets more resistant to the presence of ssDNA. **G**, Fluorescence images obtained upon mixing 15 µM SSB, 0.3 µM fluorescein-labeled SSB and the indicated concentrations of poly-dT (homo-deoxythymidine ssDNA with average length 600-100 nt) in the absence of crowding agents in LLPS buffer. **H-J**, Apparent droplet diameters determined from fluorescence images (**Fig. 5B**, panel **F**) obtained with dT_79_ in the absence (panel **H**, cf. **Fig. 5B**) and presence (panel **I**, cf. panel **F**) of 150 mg/ml BSA and with poly-dT in the absence of molecular crowders (panel **J**, cf. panel **F**). Mean ± SD (circles and error bars) and medians (horizontal lines) values are shown next to data points. Due to the non-ideal optical behavior observed in PEG20000 (cf. **Fig. S4F**) we were unable to reliably determine droplet diameters. However, we note that in the presence of PEG, we usually observed a higher fraction of larger droplets with increasing ssDNA concentration. **K**, Dependence of droplet diameter from poly-dT concentration (medians and 25/75 percentiles shown as bullets and dashes, respectively). Based on linear regression, poly-dT had no statistically significant effect on droplet diameter (the slope was not significantly different from 0, *p* > 0.05, one-way ANOVA). See also **Fig. 5C** for results obtained with dT_79_ in the presence and absence of BSA.

**Supplementary Figure 8.**
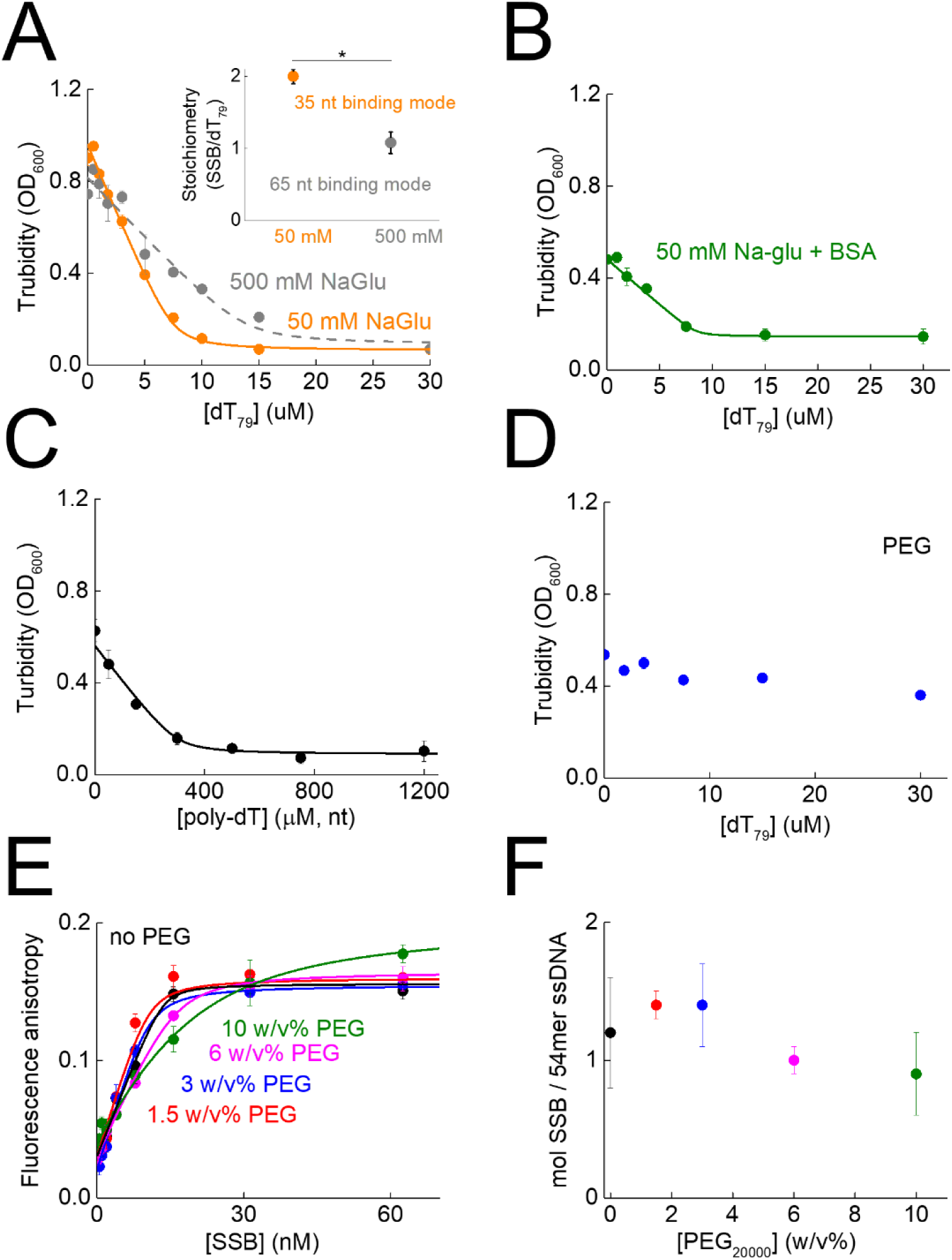
Turbidity measurements in the absence of ssDNA confirm the inhibitory effect of ssDNA binding to SSB on droplet formation. **A**, Turbidity of 15 µM SSB in LLPS buffer containing 50 or 500 mM NaGlu and increasing concentrations of dT_79_. The turbidity decreased quasi-linearly with increasing dT_79_ concentration and reached saturation. Based on fluorescence microscopic results, turbidity decrease indicates a decrease in the number of droplets in the solution. Solid lines represent fits using a quadratic binding equation (**Equation 2**, see Methods) used to determine the SSB:dT_79_ stoichiometry where saturation occurred. In the quadratic binding model, we assume that, due to the high concentration of SSB and its high affinity to ssDNA, binding of DNA molecules is stoichiometric at the applied concentrations. Determined stoichiometries (inset) at the two salt concentrations were significantly different (*p* < 0.05, *n* = 3, unpaired T-test). In 50 mM NaGlu, on average two SSB molecules bind to one dT_79_ molecule, whereas in 500 mM NaGlu, one SSB is bound per dT_79_, in line with the previously described preference for the 35-nt and 65-nt ssDNA binding modes of SSB at low and high salt concentrations, respectively (Lohman and Ferrari, 1994). **B**, Turbidity of 15 µM SSB in LLPS buffer at increasing concentrations of poly-dT. The solid line represents best-fit using a quadratic binding equation (**Equation 2**). The determined apparent SSB binding site size on ssDNA (mole nt /mol SSB where saturation occurs) calculated from the stoichiometry values is 2.2 ± 0.4 fold smaller for poly-dT than for dT_79_. These results may be indicative of an additional effect of ssDNA length besides binding to SSB, as the determined apparent binding site size (17 ± 3 nt/SSB) is smaller than that for the 35-nt binding mode of SSB. **C**, Turbidity of 15 µM SSB in LLPS buffer containing 150 mg/ml BSA and increasing concentrations of dT_79_. The solid line represents best-fit using a quadratic binding equation (**Equation 2**). The presence of BSA did not significantly affect the stoichiometry value compared to SSB alone (no BSA: 2.09 ± 0.04 SSB per dT_79_; BSA: 1.78 ± 0.18 SSB per dT_79_; *p* > 0.05, *n* = 3, unpaired T-test). This finding indicates the BSA as a molecular crowder does not influence the inhibitory effect of ssDNA binding to SSB on droplet formation. **D**, Turbidity of 15 µM SSB in LLPS buffer containing 10 w/v% PEG20000 and increasing concentrations of dT_79_. In line with fluorescence microscopic results (**Fig. S7F**), in PEG the droplets were more resistant to the presence of ssDNA. These results show that the physicochemical properties of different molecular crowders can influence droplet behavior markedly differently as this effect is only observed for PEG and not for BSA. **E-F**, To test whether PEG20000 confers stability of droplets by decreasing the ssDNA binding affinity of SSB, we performed fluorescence anisotropy-based binding experiments in which 15 nM fluorescein-labeled 54mer oligonucleotide was titrated with increasing SSB concentrations in LLPS buffer at various PEG20000 concentrations. Solid lines show fits using the quadratic binding equation (**Equation 2**). Due to the strong affinity of SSB to ssDNA, we were unable to determine the precise *K*_d_ up to the 6 w/v% PEG concentration (*K*_d_ < 1 nM). However, in the case of 10 w/v% PEG we observed reduced but still relatively strong binding (*K*_d_ = 6.8 ± 0.75 nM). The stoichiometry of binding (panel **F**) was not influenced by PEG (*p* = 0.613, *n* = 2, one-way ANOVA). As the binding to ssDNA is still strong in the presence of PEG, these results indicate that the specific physicochemical properties and molecular crowding effect of PEG20000, which is different from that of BSA, stabilizes droplets in the presence of ssDNA. Parameters determined from turbidity measures are shown in **Table S4**.

**Supplementary Figure 9.**
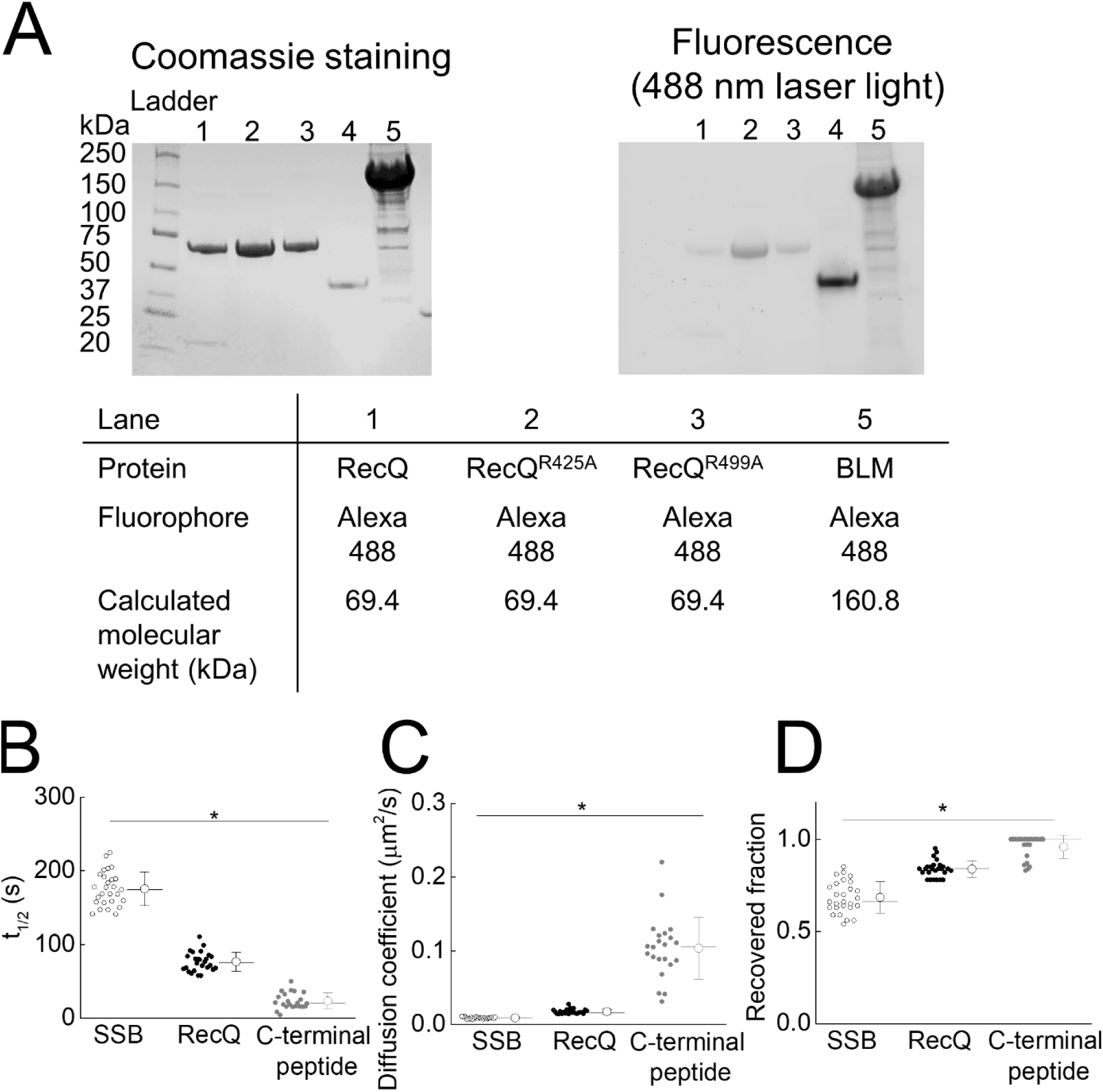
Proteins used for testing the enrichment of SSB interaction partners in SSB droplets, and diffusion properties of RecQ helicase and the isolated SSB C-terminal peptide within SSB droplets. **A**, Alexa Fluor 488 (Alexa488)-labeled *E. coli* RecQ helicase, RecQ^R499A^ and RecQ^R425A^ and human BLM helicase (8 µg each except for BLM, 32 µg) were separated using SDS-PAGE with a Mini-PROTEAN TGX 4-20 % precast gel (Bio-Rad) and visualized by Coomassie staining (left) and fluorescence detection (right) via a gel scanner instrument (488-nm excitation). Precious Plus Protein Standard (Bio Rad) was used as molecular weight marker. Calculated molecular weights are indicated in the table. Sample in lane 4 is not relevant to this study. **B-D**, Diffusion of Alexa488-labeled RecQ and fluorescein-labeled isolated SSB CTP peptide in SSB droplets. 30 µM SSB and 0.3 µM fluorescent RecQ and 0.1 µM Cy3-labeled dT_79_ were mixed in LLPS buffer. Droplets were allowed to settle to the bottom of microscopy chamber during a 2-h incubation. Half-lives of fluorescence recovery (*t*_1/2_) and recovered fractions were determined as in **Fig. 2J**. Diffusion coefficients (panel **C**) were calculated from the photobleached area and *t*_1/2_ (panel **A**). The diffusion coefficient of dT_79_ was significantly higher compared to that of SSB (* *p* < 0.05, one-way ANOVA). The recovered fraction (panel **D**) for dT_79_ was also significantly higher compared to that of SSB (* *p* < 0.05, one-way ANOVA). Mean ± SD values for t_1/2_ values, diffusion coefficients and recovered fractions are shown in **Table S3**. The diffusion coefficient for RecQ was significantly higher compared to that for SSB (effect size 2.0 ± 0.4-fold), whereas SSB CTP diffused more rapidly than SSB (11.4 ± 1.3-fold) and RecQ (5.7 ± 1.0-fold), in line with its smaller size and weaker interaction with SSB (**Fig. 6D-E**). A similar trend was observed for recovered fractions (panel **D**).

**Supplementary Figure 10.**
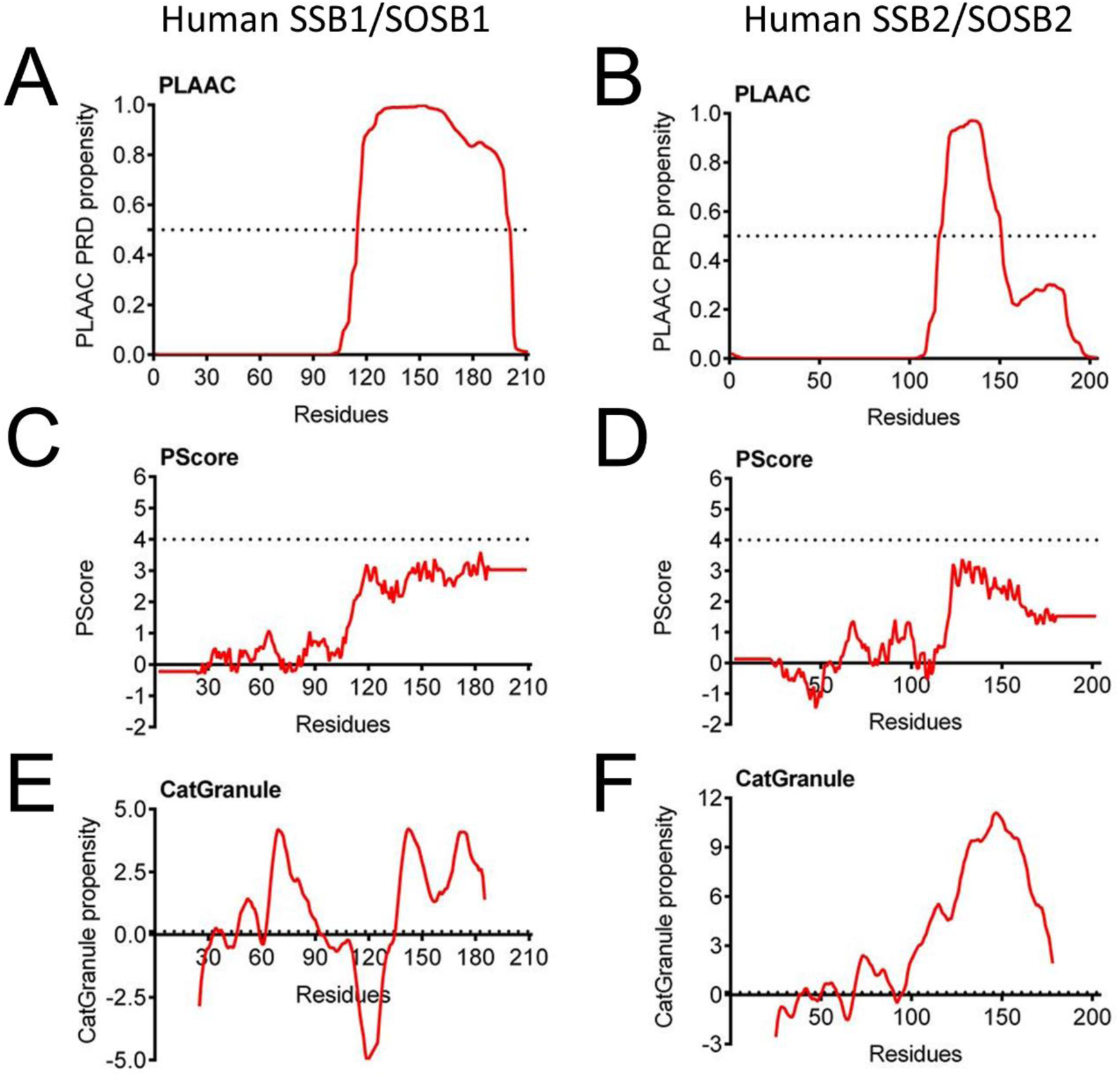
Human SSB1 and SSB2 show positive predicted LLPS propensities in their IDLs. **A-B**, The PLAAC (prion-like amino acid composition), **C-D**, PScore and **E-F**, CatGranule algorithms indicate prion-like features (PLAAC) and a propensity for LLPS (PScore, CatGranule) for the C-terminal region of the two human SSB homologs, hSSB1/SOSB1 and hSSB2/SOSB2. Predicted propensities are shown as red lines, while the minimum threshold values of the methods are indicated as black dashed lines on each graph. For the CatGranule method the threshold value is at *y* = 0; the dashed line is slightly shifted to enhance its visibility. Note that CatGranule does not provide predicted values for protein termini, resulting in its prediction curve being shorter than the SSB sequence.

### SUPPLEMENTARY VIDEO LEGENDS

**Supplementary Video 1. Fusion of phase-separated SSB droplets**

The fluorescence signal of droplets formed by 30 µM SSB and 0.3 µM fluorescein-labeled SSB in LLPS buffer (20 mM Tris-OAc pH 7.5, 50 mM NaGlu, 5 mM MgOAc, 20 mM NaCl) was followed in time using epifluorescence microscopy. The video was background corrected.

**Supplementary Video 2. ssDNA disperses phase-separated SSB droplets**

The fluorescence signal of droplets formed by 30 µM SSB and 0.3 µM fluorescein-labeled SSB in LLPS buffer (20 mM Tris-OAc pH 7.5, 50 mM NaGlu, 5 mM MgOAc, 20 mM NaCl) was followed in time after dropwise addition of a solution of dT_79_ (2 µl, dissolved in water) into the microscopy chamber to reach a final concentration of 30 µM dT_79_. As dT_79_ mixes with the phase-separated SSB solution, SSB droplets disperse. The video was background corrected.

**Supplementary Video 3. Control measurement for dT_79_ addition experiment**

The experiment was performed similarly to that in **Supplementary Video 2** but, instead of dT_79_ solution, water was dropped into the phase-separated SSB solution. SSB droplets remained intact, supporting that dT_79_ caused dispersion.

### SUPPLEMENTARY TABLES

***Supplementary Table 1. LLPS propensity of representative bacterial SSB proteins, determined by the PLAAC and CatGranule methods, summarized for 15 major phylogenetic groups of Bacteria***

***Supplementary Table 2. LLPS propensities of representative bacterial SSB proteins by the PLAAC and CatGranule methods***

**Supplementary Table 3.** Mean ± SD values of t_1/2_, diffusion coefficients and recovered fractions determined from FRAP experiments.

| Fluorescent molecule examined with SSB droplet | $t_{1/2}$ (s) | Diffusion coefficient ($\mu\text{m}^2/\text{s}$) | Recovered fraction | $n$ (droplets photobleached) |
| --- | --- | --- | --- | --- |
| Fluorescein – SSB | 175.5 ± 23.1 | 0.009 ± 0.001 | 0.68 ± 0.08 | 27 |
| Fluorescein - SSB (10 w/v% PEG<sub>20000</sub>) | 321.4 ± 150.8 | 0.002 ± 0.002 | 0.71 ± 0.20 | 22 |
| Alexa 555 – SSBdC | 51.1 ± 11.8 | 0.051 ± 0.017 | 0.99 ± 0.03 | 8 |
| Cy3 - dT<sub>79</sub> | 63.1 ± 31.6 | 0.020 ± 0.004 | 0.92 ± 0.06 | 10 |
| eGFP-GGS<sub>4</sub>-CTP (10 w/v% PEG<sub>20000</sub>) | 144.2 ± 24.6 | 0.008 ± 0.002 | 0.91 ± 0.06 | 31 |
| eGFP-IDL-CTP (10 w/v% PEG<sub>20000</sub>) | 154.1 ± 18. | 0.009 ± 0.002 | 0.86 ± 0.06 | 35 |
| Alexa488 – RecQ | 76.6 ± 13.0 | 0.018 ± 0.003 | 0.84 ± 0.05 | 26 |
| Fluorescein – CTP | 23.4 ± 10.9 | 0.103 ± 0.042 | 0.96 ± 0.06 | 22 |

**Supplementary Table 4.** Results of fits using the quadratic binding equation to ssDNA-dependent turbidity measurements.

| [SSB] ($\mu\text{M}$) titrated with dT<sub>79</sub> in 50 mM NaGlu | $K_d$ ($\mu\text{M}$)<sup>a</sup> | stoichiometry (mol SSB/mol DNA molecule) (best-fit parameter and fitting error) | Apparent binding site size (nt/SSB) |
| --- | --- | --- | --- |
| 2.5 | <0.3 | 1.99 ± 0.33 | 39.7 ± 6.6 |
| 5 | <0.3 | 2.11 ± 0.40 | 37.5 ± 7.1 |
| 10 | <0.3 | 2.16 ± 0.12 | 36.5 ± 2.2 |
| 15 | <0.3 | 2.09 ± 0.04 | 37.7 ± 2.1 |
| 30 | <0.3 | 2.38 ± 0.13 | 35.9 ± 0.9 |
| [SSB] ($\mu\text{M}$) titrated with dT<sub>79</sub> in 500 mM NaGlu | | | |
| 15 | <0.3 | 1.08 ± 0.15 | 73.1 ± 10.2 |
| [SSB] ($\mu\text{M}$) titrated with dT<sub>79</sub> in BSA and 50 mM NaGlu | | | |
| 15 | <0.3 | 1.78 ± 0.18 | 44.4 ± 4.5 |
| [SSB] ($\mu\text{M}$) titrated with dT<sub>79</sub> in PEG<sub>20000</sub> and 50 mM NaGlu | | | |
| 15 | n.d. | n.d. | n.d. |
| [SSB] ($\mu\text{M}$) titrated with poly-dT in 50 mM NaGlu | | | |
| 15 | 0.9 ± 0.5 ($\mu\text{M}$, nt) | 0.06 ± 0.01 (mol SSB / mol nt) | 17 ± 3 |

<sup>a</sup> Due to strong binding of SSB to ssDNA, dissociation constants could not be determined precisely and an upper limit is indicated.

**Supplementary Table 5.** Dissociation constants for binding experiments obtained upon fitting the Hill binding equation (Equation 1)

| Binding reaction | $K_d$ ($\mu\text{M}$)<br>(best-fit<br>parameter and<br>fitting error) | $n$ (Hill<br>coefficient)<br>(best-fit<br>parameter and<br>fitting error) |
| --- | --- | --- |
| SSB + Flu-CTP | $15.15 \pm 3.74$ | $1.04 \pm 0.20$ |
| SSB + dT<sub>79</sub> + Flu-CTP | n.d. | n.d. |
| RecQ + Flu-CTP | $0.34 \pm 0.03$ | $1.40 \pm 0.19$ |
| RecQ<sup>R425A</sup> + Flu-CTP | $17.20 \pm 0.99$ | $0.94 \pm 0.04$ |
| RecQ<sup>R499A</sup> + Flu-CTP | $59.22 \pm 7.74$ | $1.11 \pm 0.24$ |
| BLM + Flu-CTP | n.d. | n.d. |
| SSB + Flu-Control peptide | n.d. | n.d. |
| SSB + Flu-dCTP | n.d. | n.d. |
| SSB + Fluorescein | n.d. | n.d. |

n.d.: Binding was not observed.

## REFERENCES

Alberti, S., and Dormann, D. (2019). Liquid-Liquid Phase Separation in Disease. Annu. Rev. Genet.

Alberti, S., Gladfelter, A., and Mittag, T. (2019). Considerations and Challenges in Studying Liquid-Liquid Phase Separation and Biomolecular Condensates. Cell 176, 419–434.

Al-Husini, N., Tomares, D.T., Bitar, O., Childers, W.S., and Schrader, J.M. (2018). α-Proteobacterial RNA Degradosomes Assemble Liquid-Liquid Phase-Separated RNP Bodies. Mol. Cell 71, 1027–1039.e14.

Antony, E., and Lohman, T.M. (2019). Dynamics of E. coli single stranded DNA binding (SSB) protein-DNA complexes. Semin. Cell Dev. Biol. 86, 102–111.

Antony, E., Weiland, E., Yuan, Q., Manhart, C.M., Nguyen, B., Kozlov, A.G., McHenry, C.S., and Lohman, T.M. (2013). Multiple C-terminal tails within a single E. coli SSB homotetramer coordinate DNA replication and repair. J. Mol. Biol. 425, 4802–4819.

Ashton, N.W., Bolderson, E., Cubeddu, L., O’Byrne, K.J., and Richard, D.J. (2013). Human single-stranded DNA binding proteins are essential for maintaining genomic stability. BMC Mol. Biol. 14, 9.

Bakthavachalu, B., Huelsmeier, J., Sudhakaran, I.P., Hillebrand, J., Singh, A., Petrauskas, A., Thiagarajan, D., Sankaranarayanan, M., Mizoue, L., Anderson, E.N., et al. (2018). RNP-Granule Assembly via Ataxin-2 Disordered Domains Is Required for Long-Term Memory and Neurodegeneration. Neuron 98, 754–766.e4.

Banani, S.F., Rice, A.M., Peeples, W.B., Lin, Y., Jain, S., Parker, R., and Rosen, M.K. (2016). Compositional Control of Phase-Separated Cellular Bodies. Cell 166, 651–663.

Banani, S.F., Lee, H.O., Hyman, A.A., and Rosen, M.K. (2017). Biomolecular condensates: organizers of cellular biochemistry. Nat. Rev. Mol. Cell Biol. 18, 285–298.

Bennett, B.D., Kimball, E.H., Gao, M., Osterhout, R., Van Dien, S.J., and Rabinowitz, J.D. (2009). Absolute metabolite concentrations and implied enzyme active site occupancy in Escherichia coli. Nat. Chem. Biol. 5, 593–599.

Bianco, P.R. (2017). The tale of SSB. Prog. Biophys. Mol. Biol. 127, 111–118.

Bianco, P.R., Pottinger, S., Tan, H.Y., Nguyenduc, T., Rex, K., and Varshney, U. (2017). The IDL of *E. coli* SSB links ssDNA and protein binding by mediating protein-protein interactions: SSB IDL Links ssDNA and Protein Binding. Protein Sci. 26, 227–241.

Bolognesi, B., Lorenzo Gotor, N., Dhar, R., Cirillo, D., Baldrighi, M., Tartaglia, G.G., and Lehner, B. (2016). A Concentration-Dependent Liquid Phase Separation Can Cause Toxicity upon Increased Protein Expression. Cell Rep. 16, 222–231.

Curth, U., Genschel, J., Urbanke, C., and Greipel, J. (1996). In vitro and in vivo function of the C-terminus of Escherichia coli single-stranded DNA binding protein. Nucleic Acids Res. 24, 2706–2711.

Dillingham, M.S., Tibbles, K.L., Hunter, J.L., Bell, J.C., Kowalczykowski, S.C., and Webb, M.R. (2008). Fluorescent single-stranded DNA binding protein as a probe for sensitive, real-time assays of helicase activity. Biophys. J. 95, 3330–3339.

Gyimesi, M., Harami, G.M., Sarlós, K., Hazai, E., Bikádi, Z., and Kovács, M. (2012). Complex activities of the human Bloom’s syndrome helicase are encoded in a core region comprising the RecA and Zn-binding domains. Nucleic Acids Res. 40, 3952–3963.

Gyimesi, M., Pires, R.H., Billington, N., Sarlós, K., Kocsis, Z.S., Módos, K., Kellermayer, M.S.Z., and Kovács, M. (2013). Visualization of human Bloom’s syndrome helicase molecules bound to homologous recombination intermediates. FASEB J. 27, 4954–4964.

Harami, G.M., Seol, Y., In, J., Ferencziová, V., Martina, M., Gyimesi, M., Sarlós, K., Kovács, Z.J., Nagy, N.T., Sun, Y., et al. (2017). Shuttling along DNA and directed processing of D-loops by RecQ helicase support quality control of homologous recombination. Proc. Natl. Acad. Sci. 114, E466–E475.

Heinkel, F., Abraham, L., Ko, M., Chao, J., Bach, H., Hui, L.T., Li, H., Zhu, M., Ling, Y.M., Rogalski, J.C., et al. (2019). Phase separation and clustering of an ABC transporter in Mycobacterium tuberculosis. Proc. Natl. Acad. Sci. U. S. A. 116, 16326–16331.

Heinrich, B.S., Maliga, Z., Stein, D.A., Hyman, A.A., and Whelan, S.P.J. (2018). Phase Transitions Drive the Formation of Vesicular Stomatitis Virus Replication Compartments. MBio 9.

Kang, M., Day, C.A., Kenworthy, A.K., and DiBenedetto, E. (2012). Simplified equation to extract diffusion coefficients from confocal FRAP data. Traffic Cph. Den. 13, 1589–1600.

Kozlov, A.G., and Lohman, T.M. (2002). Kinetic mechanism of direct transfer of Escherichia coli SSB tetramers between single-stranded DNA molecules. Biochemistry 41, 11611–11627.

Kozlov, A.G., and Lohman, T.M. (2006). Effects of monovalent anions on a temperature-dependent heat capacity change for Escherichia coli SSB tetramer binding to single-stranded DNA. Biochemistry 45, 5190–5205.

Kozlov, A.G., Cox, M.M., and Lohman, T.M. (2010). Regulation of single-stranded DNA binding by the C termini of Escherichia coli single-stranded DNA-binding (SSB) protein. J. Biol. Chem. 285, 17246–17252.

Kozlov, A.G., Weiland, E., Mittal, A., Waldman, V., Antony, E., Fazio, N., Pappu, R.V., and Lohman, T.M. (2015). Intrinsically disordered C-terminal tails of E. coli single-stranded DNA binding protein regulate cooperative binding to single-stranded DNA. J. Mol. Biol. 427, 763–774.

Kozlov, A.G., Shinn, M.K., Weiland, E.A., and Lohman, T.M. (2017). Glutamate promotes SSB protein-protein Interactions via intrinsically disordered regions. J. Mol. Biol. 429, 2790–2801.

Kuznetsova, I.M., Turoverov, K.K., and Uversky, V.N. (2014). What macromolecular crowding can do to a protein. Int. J. Mol. Sci. 15, 23090–23140.

Lancaster, A.K., Nutter-Upham, A., Lindquist, S., and King, O.D. (2014). PLAAC: a web and command-line application to identify proteins with prion-like amino acid composition. Bioinforma. Oxf. Engl. 30, 2501–2502.

Lawson, T., El-Kamand, S., Kariawasam, R., Richard, D.J., Cubeddu, L., and Gamsjaeger, R. (2019). A Structural Perspective on the Regulation of Human Single-Stranded DNA Binding Protein 1 (hSSB1, OBFC2B) Function in DNA Repair. Comput. Struct. Biotechnol. J. 17, 441–446.

Lin, Y.-H., Forman-Kay, J.D., and Chan, H.S. (2018). Theories for Sequence-Dependent Phase Behaviors of Biomolecular Condensates. Biochemistry 57, 2499–2508.

Lohman, T.M., and Ferrari, M.E. (1994). Escherichia coli single-stranded DNA-binding protein: multiple DNA-binding modes and cooperativities. Annu. Rev. Biochem. 63, 527–570.

Mannen, T., Yamashita, S., Tomita, K., Goshima, N., and Hirose, T. (2016). The Sam68 nuclear body is composed of two RNase-sensitive substructures joined by the adaptor HNRNPL. J. Cell Biol. 214, 45–59.

Meyer, R.R., and Laine, P.S. (1990). The single-stranded DNA-binding protein of Escherichia coli. Microbiol. Rev. 54, 342–380.

Mills, M., Harami, G.M., Seol, Y., Gyimesi, M., Martina, M., Kovács, Z.J., Kovács, M., and Neuman, K.C. (2017). RecQ helicase triggers a binding mode change in the SSB–DNA complex to efficiently initiate DNA unwinding. Nucleic Acids Res. 45, 11878–11890.

Milovanovic, D., Wu, Y., Bian, X., and De Camilli, P. (2018). A liquid phase of synapsin and lipid vesicles. Science 361, 604–607.

Mittag, T., and Parker, R. (2018). Multiple Modes of Protein-Protein Interactions Promote RNP Granule Assembly. J. Mol. Biol. 430, 4636–4649.

Molineux, I.J., Pauli, A., and Gefter, M.L. (1975). Physical studies of the interaction between the Escherichia coli DNA binding protein and nucleic acids. Nucleic Acids Res. 2, 1821–1837.

Monterroso, B., Zorrilla, S., Sobrinos-Sanguino, M., Robles-Ramos, M.A., López-Álvarez, M., Margolin, W., Keating, C.D., and Rivas, G. (2019). Bacterial FtsZ protein forms phase-separated condensates with its nucleoid-associated inhibitor SlmA. EMBO Rep. 20.

Mukherjee, S., Stamatis, D., Bertsch, J., Ovchinnikova, G., Verezemska, O., Isbandi, M., Thomas, A.D., Ali, R., Sharma, K., Kyrpides, N.C., et al. (2017). Genomes OnLine Database (GOLD) v.6: data updates and feature enhancements. Nucleic Acids Res. 45, D446–D456.

Nikolic, J., Le Bars, R., Lama, Z., Scrima, N., Lagaudrière-Gesbert, C., Gaudin, Y., and Blondel, D. (2017). Negri bodies are viral factories with properties of liquid organelles. Nat. Commun. 8, 58.

Nikolic, J., Lagaudrière-Gesbert, C., Scrima, N., Blondel, D., and Gaudin, Y. (2018). [Rabies virus factories are formed by liquid-liquid phase separation]. Med. Sci. MS 34, 203–205.

Pancsa, R., and Tompa, P. (2016). Essential functions linked with structural disorder in organisms of minimal genome. Biol. Direct 11, 45.

Pancsa, R., Schad, E., Tantos, A., and Tompa, P. (2019a). Emergent functions of proteins in non-stoichiometric supramolecular assemblies. Biochim. Biophys. Acta Proteins Proteomics 1867, 970–979.

Pancsa, R., Kovacs, D., and Tompa, P. (2019b). Misprediction of Structural Disorder in Halophiles. Mol. Basel Switz. 24.

Paradžik, T., Filić, Ž., and Vujaklija, D. (2017). Variations in amino acid composition in bacterial single stranded DNA–binding proteins correlate with GC content. Period. Biol. 118.

Raghunathan, S., Ricard, C.S., Lohman, T.M., and Waksman, G. (1997). Crystal structure of the homo-tetrameric DNA binding domain of Escherichia coli single-stranded DNA-binding protein determined by multiwavelength x-ray diffraction on the selenomethionyl protein at 2.9-A resolution. Proc. Natl. Acad. Sci. U. S. A. 94, 6652–6657.

Rapsomaniki, M.A., Kotsantis, P., Symeonidou, I.-E., Giakoumakis, N.-N., Taraviras, S., and Lygerou, Z. (2012). easyFRAP: an interactive, easy-to-use tool for qualitative and quantitative analysis of FRAP data. Bioinforma. Oxf. Engl. 28, 1800–1801.

Reimer, L.C., Vetcininova, A., Carbasse, J.S., Söhngen, C., Gleim, D., Ebeling, C., and Overmann, J. (2019). BacDive in 2019: bacterial phenotypic data for High-throughput biodiversity analysis. Nucleic Acids Res. 47, D631–D636.

Sabari, B.R., Dall’Agnese, A., Boija, A., Klein, I.A., Coffey, E.L., Shrinivas, K., Abraham, B.J., Hannett, N.M., Zamudio, A.V., Manteiga, J.C., et al. (2018). Coactivator condensation at super-enhancers links phase separation and gene control. Science 361.

Sarlós, K., Gyimesi, M., and Kovács, M. (2012). RecQ helicase translocates along single-stranded DNA with a moderate processivity and tight mechanochemical coupling. Proc. Natl. Acad. Sci. U. S. A. 109, 9804–9809.

Savvides, S.N., Raghunathan, S., Fütterer, K., Kozlov, A.G., Lohman, T.M., and Waksman, G. (2004). The C-terminal domain of full-length E. coli SSB is disordered even when bound to DNA. Protein Sci. Publ. Protein Soc. 13, 1942–1947.

Schmidt, A., Kochanowski, K., Vedelaar, S., Ahrné, E., Volkmer, B., Callipo, L., Knoops, K., Bauer, M., Aebersold, R., and Heinemann, M. (2016). The quantitative and condition-dependent Escherichia coli proteome. Nat. Biotechnol. 34, 104–110.

Schultz, S.G., Wilson, N.L., and Epstein, W. (1962). Cation transport in Escherichia coli. II. Intracellular chloride concentration. J. Gen. Physiol. 46, 159–166.

Shereda, R.D., Bernstein, D.A., and Keck, J.L. (2007). A central role for SSB in Escherichia coli RecQ DNA helicase function. J. Biol. Chem. 282, 19247–19258.

Shereda, R.D., Kozlov, A.G., Lohman, T.M., Cox, M.M., and Keck, J.L. (2008). SSB as an organizer/mobilizer of genome maintenance complexes. Crit. Rev. Biochem. Mol. Biol. 43, 289–318.

Shereda, R.D., Reiter, N.J., Butcher, S.E., and Keck, J.L. (2009). Identification of the SSB Binding Site on E. coli RecQ Reveals a Conserved Surface for Binding SSB’s C Terminus. J. Mol. Biol. 386, 612–625.

Shin, Y., and Brangwynne, C.P. (2017). Liquid phase condensation in cell physiology and disease. Science 357.

Shishmarev, D., Wang, Y., Mason, C.E., Su, X.-C., Oakley, A.J., Graham, B., Huber, T., Dixon, N.E., and Otting, G. (2014). Intramolecular binding mode of the C-terminus of Escherichia coli single-stranded DNA binding protein determined by nuclear magnetic resonance spectroscopy. Nucleic Acids Res. 42, 2750–2757.

Su, X.-C., Wang, Y., Yagi, H., Shishmarev, D., Mason, C.E., Smith, P.J., Vandevenne, M., Dixon, N.E., and Otting, G. (2014). Bound or free: interaction of the C-terminal domain of Escherichia coli single-stranded DNA-binding protein (SSB) with the tetrameric core of SSB. Biochemistry 53, 1925–1934.

Vernon, R.M., and Forman-Kay, J.D. (2019). First-generation predictors of biological protein phase separation. Curr. Opin. Struct. Biol. 58, 88–96.

Vernon, R.M., Chong, P.A., Tsang, B., Kim, T.H., Bah, A., Farber, P., Lin, H., and Forman-Kay, J.D. (2018). Pi-Pi contacts are an overlooked protein feature relevant to phase separation. ELife 7.

Wu, C., and Nebert, D.W. (2004). Update on genome completion and annotations: Protein Information Resource. Hum. Genomics 1, 229–233.

Yu, C., Tan, H.Y., Choi, M., Stanenas, A.J., Byrd, A.K., D Raney, K., Cohan, C.S., and Bianco, P.R. (2016). SSB binds to the RecG and PriA helicases in vivo in the absence of DNA. Genes Cells Devoted Mol. Cell. Mech. 21, 163–184.

Zhao, T., Liu, Y., Wang, Z., He, R., Zhang, J.X., Xu, F., Lei, M., Deci, M.B., Nguyen, J., and Bianco, P.R. (2019). Super-resolution imaging reveals changes in SSB localization in response to DNA damage. BioRxiv 664532. doi: https://doi.org/10.1101/664532

Zhou, Y., Su, J.M., Samuel, C.E., and Ma, D. (2019). Measles Virus Forms Inclusion Bodies with Properties of Liquid Organelles. J. Virol. doi: https://doi.org/10.1128/JVI.00948-19

## SUPPLEMENTARY REFERENCES

Raut, A.S., and Kalonia, D.S. (2016). Pharmaceutical Perspective on Opalescence and Liquid-Liquid Phase Separation in Protein Solutions. Mol. Pharm. 13, 1431–1444.

