## Supplementary Table 1 for "Phase separation by ssDNA binding protein controlled *via* protein-protein and protein-DNA interactions"

**Supplementary Table 1. LLPS propensities of representative bacterial SSB proteins by the PLAAC and catGranule methods summarized for 15 bacterial classes**

| **Bacterial Clad** | **Taxon ID** | **Species in PIR RPs 15%** | **SSB-named proteins** | **"(Fragment)"s deleted** | **SSB-like orthologs removed** | **SSBs analyzed** | **SSBs with PRD by PLAA** | **SSBs with >0.5 by CatGranule** |
|---|---|---|---|---|---|---|---|---|
| Actinobacteria | 201174 | **80** | 185 | 0 | 96 | **89** | **46** | **78** |
| Alpha-proteobacteria | 28211 | **67** | 83 | 1 | 12 | **70** | **13** | **62** |
| Aquificae | 200783 | **4** | 4 | 0 | 0 | **4** | **0** | **1** |
| Beta-proteobacteria | 28216 | **30** | 44 | 1 | 14 | **29** | **0** | **23** |
| Bacteroidetes | 976 | **56** | 83 | 0 | 24 | **59** | **5** | **32** |
| Chlamydiae | 204428 | **3** | 4 | 0 | 1 | **3** | **1** | **3** |
| Cyanobacteria | 1117 | **5** | 7 | 0 | 1 | **6** | **0** | **1** |
| Deinococcus-Thermus | 1297 | **3** | 5 | 0 | 2 | **3** | **0** | **3** |
| Delta/Epsilon-proteobacteria | 68525 | **48** | 64 | 2 | 11 | **51** | **21** | **36** |
| Firmicutes | 1239 | **222** | 361 | 3 | 92 | **266** | **52** | **152** |
| Fusobacteria | 32066 | **6** | 9 | 0 | 1 | **8** | **0** | **4** |
| Gamma-proteobacteria | 1236 | **101** | 129 | 5 | 17 | **107** | **61** | **88** |
| Chlorobi | 1090 | **1** | 2 | 0 | 1 | **1** | **1** | **1** |
| Planctomycetes | 203682 | **18** | 20 | 0 | 2 | **18** | **4** | **18** |
| Thermotogae | 200918 | **3** | 4 | 0 | 1 | **3** | **0** | **0** |
| **Sum** | **-** | **647** | **1004** | **12** | **275** | **717** | **204** | **502** |

*PIR release 2019 06*
