## Supplementary Table 2 for "Phase separation by ssDNA binding protein controlled *via* protein-protein and protein-DNA interactions"

**Supplementary Table S2: LLPS propensities of representative bacterial SSB proteins by the PLAAC and CatGranule methods**

| UniProt AC | Taxon ID | Class taxon I | CatGranule value | CatGranule Positiv | PLAAC score | PLAAC PRD length | Has PLAAC PRD? Y | Positive with 0/1/2 methods? | IDL Length | Remarks |
|---|---|---|---|---|---|---|---|---|---|---|
| A0A0C3RCU9 | 1547597 | 976 | 0.23118 | 0 | NaN | 0 | 0 | 0 | 35 | |
| A0A1T5ASN4 | 889453 | 976 | 1.02838 | 1 | 13.932 | 38 | 1 | 2 | 57 | |
| Q11NU5 | 269798 | 976 | 1.23245 | 1 | NaN | 0 | 0 | 1 | 47 | |
| R7F3A0 | 1262927 | 976 | -0.07431 | 0 | NaN | 0 | 0 | 0 | 6 | MOTIF MISSING, FRAGMENT? |
| H6L379 | 984262 | 976 | 0.81001 | 1 | NaN | 0 | 0 | 1 | 60 | |
| M7N0N8 | 1279009 | 976 | 1.67033 | 1 | 10.376 | 52 | 1 | 2 | 53 | |
| A0A085L0J3 | 1453500 | 976 | 0.551779 | 1 | NaN | 0 | 0 | 1 | 45 | |
| A0A1F1DYT8 | 1739394 | 976 | -0.340872 | 0 | NaN | 0 | 0 | 0 | 39 | |
| A0A0M3CBN8 | 1338009 | 976 | 0.277148 | 0 | NaN | 0 | 0 | 0 | 44 | |
| A0A0M3CEV4 | 1338009 | 976 | 0.220055 | 0 | NaN | 0 | 0 | 0 | 26 | HAS ORTHOLOG WITH LARGER LLPS PROPENSITY |
| A0A0S2I3P2 | 1307839 | 976 | 0.62769 | 1 | NaN | 0 | 0 | 1 | 41 | |
| A0A1Q3GZI5 | 1905359 | 976 | 1.10242 | 1 | NaN | 0 | 0 | 1 | 44 | |
| A0A2A3UG51 | 2021370 | 976 | 0.705365 | 1 | NaN | 0 | 0 | 1 | 31 | |
| A0A060R893 | 1433126 | 976 | -0.100349 | 0 | NaN | 0 | 0 | 0 | 33 | |
| A0A0C1I8N4 | 1463156 | 976 | 1.02369 | 1 | 14.103 | 37 | 1 | 2 | 52 | |
| A0A0C1I6M2 | 1463156 | 976 | 0.995647 | 1 | NaN | 0 | 0 | 1 | 40 | HAS ORTHOLOG WITH LARGER LLPS PROPENSITY |
| A0A379MNE0 | 28139 | 976 | 1.02548 | 1 | NaN | 0 | 0 | 1 | 48 | |
| A0A1T5KGE0 | 1945890 | 976 | 1.20288 | 1 | 19.48 | 36 | 1 | 2 | 68 | |
| I4A2G1 | 867902 | 976 | 0.489574 | 0 | NaN | 0 | 0 | 0 | 31 | |
| A0A2N3IJ03 | 2016530 | 976 | 1.10745 | 1 | NaN | 0 | 0 | 1 | 43 | |
| U5Q9T7 | 1400053 | 976 | -0.123741 | 0 | NaN | 0 | 0 | 0 | 35 | |
| A0A0S7BUF8 | 1678841 | 976 | 0.626155 | 1 | NaN | 0 | 0 | 1 | 36 | |
| F4L5F7 | 760192 | 976 | 0.387673 | 0 | NaN | 0 | 0 | 0 | 51 | |
| B6YQX2 | 511995 | 976 | 1.01812 | 1 | NaN | 0 | 0 | 1 | 45 | |
| L0JB55 | 908937 | 976 | 0.171375 | 0 | NaN | 0 | 0 | 0 | 51 | |
| A0A2G5L616 | 1889772 | 976 | 0.479136 | 0 | NaN | 0 | 0 | 0 | 43 | |
| A0A0K8QUK3 | 1688776 | 976 | 0.343273 | 0 | NaN | 0 | 0 | 0 | 43 | |
| I4AKI7 | 880071 | 976 | 0.711818 | 1 | NaN | 0 | 0 | 1 | 61 | |
| A0A1I2AI26 | 1003 | 976 | 0.448441 | 0 | NaN | 0 | 0 | 0 | 49 | |
| A0A1I1YW39 | 385682 | 976 | 0.477045 | 0 | NaN | 0 | 0 | 0 | 64 | |
| A0A2P7TPN9 | 2116516 | 976 | 1.81371 | 1 | NaN | 0 | 0 | 1 | 45 | |
| A0A127B0V1 | 1379909 | 976 | 1.27915 | 1 | NaN | 0 | 0 | 1 | 53 | |
| A0A2P7TP60 | 2116516 | 976 | 1.2642 | 1 | NaN | 0 | 0 | 1 | 42 | HAS ORTHOLOG WITH LARGER LLPS PROPENSITY |
| A0A1G6HF62 | 1640674 | 976 | 0.386317 | 0 | NaN | 0 | 0 | 0 | 37 | |
| A6H058 | 402612 | 976 | 0.176472 | 0 | NaN | 0 | 0 | 0 | 33 | |
| A0A142EJ30 | 1727163 | 976 | 0.877901 | 1 | NaN | 0 | 0 | 1 | 51 | |
| A0A142L277 | 1690483 | 976 | 1.22478 | 1 | NaN | 0 | 0 | 1 | 7 | MOTIF MISSING, FRAGMENT? |
| A0A0B8YCZ1 | 1122941 | 976 | 0.697459 | 1 | NaN | 0 | 0 | 1 | 42 | |
| A0A401U862 | 2482724 | 976 | 0.956147 | 1 | NaN | 0 | 0 | 1 | 38 | |
| A0A2S2DRX4 | 2183547 | 976 | 0.828911 | 1 | NaN | 0 | 0 | 1 | 49 | |
| D7JFM1 | 575590 | 976 | 0.074212 | 0 | NaN | 0 | 0 | 0 | 41 | |
| A0A1I7NB04 | 1393122 | 976 | 0.718287 | 1 | NaN | 0 | 0 | 1 | 39 | MOTIF MISSING, FRAGMENT? |
| E4TPK4 | 643867 | 976 | 1.16271 | 1 | NaN | 0 | 0 | 1 | 45 | |
| A0A3S9P7G8 | 2494373 | 976 | 0.420361 | 0 | NaN | 0 | 0 | 0 | 56 | |
| G8R207 | 926562 | 976 | 1.32002 | 1 | NaN | 0 | 0 | 1 | 39 | |
| I2EUM7 | 929562 | 976 | 1.24091 | 1 | NaN | 0 | 0 | 1 | 52 | HAS ORTHOLOG WITH LARGER LLPS PROPENSITY |
| A0A180F7M9 | 1633202 | 976 | 0.585071 | 1 | NaN | 0 | 0 | 1 | 50 | |
| B3EUI3 | 452471 | 976 | 0.471736 | 0 | NaN | 0 | 0 | 0 | 31 | |
| A0A098BVY9 | 1562970 | 976 | 0.728986 | 1 | NaN | 0 | 0 | 1 | 41 | |
| A0A1H4BKP8 | 908615 | 976 | 0.48023 | 0 | NaN | 0 | 0 | 0 | 42 | |
| A0A0C1ENE9 | 510955 | 976 | 0.479119 | 0 | NaN | 0 | 0 | 0 | 5 | MOTIF MISSING, FRAGMENT? |
| A0A3N7BE77 | 2153357 | 976 | 0.384619 | 0 | NaN | 0 | 0 | 0 | 39 | |
| A0A3N7A821 | 2153357 | 976 | -0.134833 | 0 | NaN | 0 | 0 | 0 | 26 | HAS ORTHOLOG WITH LARGER LLPS PROPENSITY |
| A0A2G0CE81 | 438751 | 976 | -0.0274599 | 0 | NaN | 0 | 0 | 0 | 54 | HAS ORTHOLOG WITH LARGER LLPS PROPENSITY |
| A0A2G0CBS2 | 438751 | 976 | 0.546487 | 1 | NaN | 0 | 0 | 1 | 33 | |
| E4T8K4 | 694427 | 976 | -0.292911 | 0 | NaN | 0 | 0 | 0 | 35 | |
| R7HTQ0 | 1262924 | 976 | -0.167259 | 0 | NaN | 0 | 0 | 0 | 8 | MOTIF MISSING, FRAGMENT? |
| A0A383U2G1 | 2497989 | 976 | 0.793446 | 1 | NaN | 0 | 0 | 1 | 47 | |
| Q8A7M7 | 226186 | 976 | 0.000669754 | 0 | 22.285 | 44 | 1 | 1 | 55 | |
| A4SCY9 | 290318 | 1090 | 1.17743 | 1 | 14.511 | 52 | 1 | 2 | 61 | |
| D3ENS7 | 1453429 | 1117 | 0.321188 | 0 | NaN | 0 | 0 | 0 | 49 | |
| Q7VD81 | 167539 | 1117 | -0.398079 | 0 | NaN | 0 | 0 | 0 | 55 | |
| Q2JQ03 | 321332 | 1117 | 0.345315 | 0 | NaN | 0 | 0 | 0 | 18 | MOTIF MISSING, FRAGMENT? |
| Q7NHP8 | 251221 | 1117 | -0.0923117 | 0 | NaN | 0 | 0 | 0 | 14 | MOTIF MISSING, FRAGMENT?, HAS ORTHOLOG WITH LARGER LLPS |
| Q7NCN6 | 251221 | 1117 | 0.206431 | 0 | NaN | 0 | 0 | 0 | 19 | MOTIF MISSING, FRAGMENT? |
| Q55499 | 1111708 | 1117 | 0.816104 | 1 | NaN | 0 | 0 | 1 | 20 | MOTIF MISSING, FRAGMENT? |
| V5V910 | 470 | 1236 | 2.76703 | 1 | 23.76 | 79 | 1 | 2 | 87 | |
| C4K426 | 572265 | 1236 | 0.0909232 | 0 | NaN | 0 | 0 | 0 | 33 | |
| D4G909 | 515618 | 1236 | 0.949123 | 1 | NaN | 0 | 0 | 1 | 50 | |
| A0A143HJG4 | 252514 | 1236 | 1.86014 | 1 | 16.164 | 54 | 1 | 2 | 72 | |
| A0A317MRC3 | 886464 | 1236 | 0.588583 | 1 | NaN | 0 | 0 | 1 | 53 | |
| A0A1I3XDM9 | 930118 | 1236 | 1.79437 | 1 | 27.023 | 61 | 1 | 2 | 68 | |
| A0A3E0H756 | 748120 | 1236 | 0.526851 | 1 | NaN | 0 | 0 | 1 | 60 | |
| A0A370DJ98 | 2200906 | 1236 | 1.17594 | 1 | NaN | 0 | 0 | 1 | 43 | |
| A0A0K2DX89 | 1238 | 1236 | 1.26752 | 1 | 21.304 | 115 | 1 | 2 | 135 | |
| A0A370GYY6 | 254246 | 1236 | 0.300785 | 0 | NaN | 0 | 0 | 0 | 40 | |
| A0A0P9GLL2 | 381306 | 1236 | 2.6118 | 1 | NaN | 0 | 0 | 1 | 53 | |
| A0A099LB14 | 1535422 | 1236 | 1.83586 | 1 | 25.928 | 89 | 1 | 2 | 106 | |
| A0A220VC41 | 1755811 | 1236 | 1.20739 | 1 | 18.454 | 39 | 1 | 2 | 59 | |
| A0A0A6S9K9 | 1003181 | 1236 | 0.453995 | 0 | 16.329 | 54 | 1 | 1 | 68 | |
| I9DRF5 | 1195246 | 1236 | 1.50051 | 1 | 18.753 | 68 | 1 | 2 | 87 | |
| S3DL69 | 1236703 | 1236 | 0.82187 | 1 | NaN | 0 | 0 | 1 | 41 | |
| A0A0W1RSL9 | 1766620 | 1236 | 3.07321 | 1 | 12.228 | 58 | 1 | 2 | 73 | |
| A0A0K1XGY7 | 1697053 | 1236 | 0.492171 | 0 | 15.306 | 52 | 1 | 1 | 54 | |
| A1WZ36 | 349124 | 1236 | 1.96321 | 1 | NaN | 0 | 0 | 1 | 52 | |
| A0A0K1XDY7 | 1697053 | 1236 | 1.51435 | 1 | 26.476 | 83 | 1 | 2 | 91 | |
| A0A250KZW4 | 1432792 | 1236 | 1.01696 | 1 | NaN | 0 | 0 | 1 | 50 | |
| A0A1I2JWY5 | 1076937 | 1236 | 0.421716 | 0 | NaN | 0 | 0 | 0 | 47 | |
| A8PMM3 | 59196 | 1236 | -0.125113 | 0 | NaN | 0 | 0 | 0 | 44 | |
| A0A372BWT1 | 2301224 | 1236 | 1.44711 | 1 | NaN | 0 | 0 | 1 | 38 | |
| A0A348HFN5 | 33074 | 1236 | 1.05107 | 1 | 26.302 | 59 | 1 | 2 | 74 | HAS ORTHOLOG WITH LARGER LLPS PROPENSITY |
| A0A348HBX9 | 33074 | 1236 | 1.86994 | 1 | 26.105 | 87 | 1 | 2 | 102 | |
| A0A3M0A0X9 | 569599 | 1236 | 0.220594 | 0 | 20.541 | 94 | 1 | 1 | 109 | |
| A0A2Z6E6Y8 | 2010829 | 1236 | 1.85432 | 1 | NaN | 0 | 0 | 1 | 58 | |
| A0A1B1YQ43 | 1810504 | 1236 | 2.69682 | 1 | NaN | 0 | 0 | 1 | 60 | |
| A0A2Z2NLT5 | 1192854 | 1236 | 1.36254 | 1 | 12.837 | 38 | 1 | 2 | 54 | |
| D3RQK6 | 572477 | 1236 | 1.54182 | 1 | NaN | 0 | 0 | 1 | 60 | |
| D3RW92 | 572477 | 1236 | 0.219679 | 0 | NaN | 0 | 0 | 0 | 44 | HAS ORTHOLOG WITH LARGER LLPS PROPENSITY |
| A0A1Q2SPF8 | 1630141 | 1236 | 0.7571 | 1 | NaN | 0 | 0 | 1 | 53 | |
| Q5NE96 | 177416 | 1236 | 1.38559 | 1 | 24.569 | 56 | 1 | 2 | 56 | |
| A0A1H8USY7 | 406100 | 1236 | 2.36159 | 1 | 12.354 | 30 | 1 | 2 | 47 | |
| S6BGJ6 | 1248727 | 1236 | 0.730653 | 1 | NaN | 0 | 0 | 1 | 51 | |
| A0A1Y1R861 | 1940822 | 1236 | 1.31115 | 1 | 20.998 | 65 | 1 | 2 | 70 | |
| A0A2X0V3K1 | 179995 | 1236 | 1.625 | 1 | 20.993 | 98 | 1 | 2 | 149 | |
| B3PK66 | 498211 | 1236 | 2.05072 | 1 | 21.448 | 72 | 1 | 2 | 89 | |
| Q603V6 | 243233 | 1236 | 2.54703 | 1 | NaN | 0 | 0 | 1 | 63 | |
| A0A1R1LLS8 | 1897630 | 1236 | 0.824745 | 1 | 20.368 | 79 | 1 | 2 | 83 | |
| A0A0J8GT23 | 1513271 | 1236 | 1.31198 | 1 | 21.885 | 62 | 1 | 2 | 86 | |
| A0A4P9VSW3 | 202772 | 1236 | 1.0409 | 1 | 22.906 | 71 | 1 | 2 | 83 | |
| A0A0F7M398 | 1620392 | 1236 | 1.50299 | 1 | 11.656 | 50 | 1 | 2 | 65 | |
| A0A127FAB8 | 465721 | 1236 | 1.3697 | 1 | NaN | 0 | 0 | 1 | 44 | |
| A0A328TQ04 | 252393 | 1236 | 1.52771 | 1 | 18.269 | 65 | 1 | 2 | 70 | |
| A0A1E8CFT8 | 1524254 | 1236 | 1.49862 | 1 | 14.477 | 47 | 1 | 2 | 66 | |
| Q1N2F3 | 207949 | 1236 | 0.871644 | 1 | 22.549 | 85 | 1 | 2 | 101 | |
| A0A4Q5VMQ6 | 1889775 | 1236 | 1.69364 | 1 | 16.726 | 56 | 1 | 2 | 71 | |
| A0A3N1Y7R2 | 1750597 | 1236 | 0.941413 | 1 | NaN | 0 | 0 | 1 | 38 | |
| A0A2K8KS57 | 1336806 | 1236 | 2.13317 | 1 | 22.425 | 93 | 1 | 2 | 99 | |
| H3NWG8 | 745014 | 1236 | 0.352144 | 0 | NaN | 0 | 0 | 0 | 49 | |
| C4K6G2 | 572265 | 1236 | 0.276968 | 0 | NaN | 0 | 0 | 0 | 70 | |
| Q2S941 | 349521 | 1236 | 1.60326 | 1 | 18.034 | 89 | 1 | 2 | 100 | |
| A0A1R3VPD0 | 233100 | 1236 | 1.41475 | 1 | NaN | 0 | 0 | 1 | 56 | |
| A0A0Q9YEP6 | 437022 | 1236 | 0.244754 | 0 | NaN | 0 | 0 | 0 | 46 | |
| A0A2U2AFL4 | 472582 | 1236 | 1.2964 | 1 | 18.76 | 72 | 1 | 2 | 83 | |
| A0A1H6FGV4 | 1899563 | 1236 | -0.249328 | 0 | NaN | 0 | 0 | 0 | 53 | |
| A0A162KGN1 | 1822241 | 1236 | 0.880629 | 1 | 20.974 | 64 | 1 | 2 | 82 | |
| Q1YRM7 | 314287 | 1236 | 2.05484 | 1 | 32.181 | 46 | 1 | 2 | 64 | |
| A0A1A8TAC6 | 295068 | 1236 | 2.05786 | 1 | 17.885 | 87 | 1 | 2 | 118 | |

| | | | | | | | | | | |
|---|---|---|---|---|---|---|---|---|---|---|
| A0A193LHZ3 | 1548547 | 1236 | 1.27278 | 1 | NaN | 0 | 0 | 1 | 37 | |
| A0A1X1QJG2 | 1840472 | 1236 | 0.780051 | 1 | NaN | 0 | 0 | 1 | 44 | |
| A1AWF7 | 413404 | 1236 | 0.488577 | 0 | NaN | 0 | 0 | 0 | 54 | |
| Q5ZYL6 | 272624 | 1236 | 0.258786 | 0 | NaN | 0 | 0 | 0 | 52 | |
| C4L8J0 | 595494 | 1236 | 0.69295 | 1 | 12.424 | 67 | 1 | 2 | 84 | |
| W6M5K6 | 1400863 | 1236 | 0.787369 | 1 | NaN | 0 | 0 | 1 | 60 | |
| R4VPZ6 | 1260251 | 1236 | 0.84487 | 1 | NaN | 0 | 0 | 1 | 46 | |
| A0A2S2DZE5 | 2183582 | 1236 | 2.41832 | 1 | 23.12 | 61 | 1 | 2 | 68 | |
| E1V5N6 | 768066 | 1236 | 2.55224 | 1 | 23.223 | 74 | 1 | 2 | 98 | |
| A0A1B4XID1 | 1620215 | 1236 | 1.23419 | 1 | NaN | 0 | 0 | 1 | 49 | |
| A0A1H6UQ56 | 64971 | 1236 | 0.494736 | 0 | 18.958 | 102 | 1 | 1 | 120 | |
| A0A1H9Z104 | 1123402 | 1236 | 0.976795 | 1 | NaN | 0 | 0 | 1 | 76 | |
| A0A395JGV0 | 644221 | 1236 | 1.8768 | 1 | 17.528 | 45 | 1 | 2 | 62 | |
| A0A3N2D590 | 933926 | 1236 | 0.400546 | 0 | 22.309 | 86 | 1 | 1 | 103 | |
| A0A1Y1SE80 | 1317117 | 1236 | 3.25311 | 1 | 14.843 | 51 | 1 | 2 | 61 | |
| Q0VSH0 | 393595 | 1236 | 2.63483 | 1 | 19.9 | 48 | 1 | 2 | 65 | |
| K2IWX5 | 745411 | 1236 | 4.12593 | 1 | 23.397 | 100 | 1 | 2 | 119 | |
| A5EWP4 | 246195 | 1236 | 0.320943 | 0 | NaN | 0 | 0 | 0 | 63 | |
| A0A1P8UFP8 | 1765967 | 1236 | 0.608888 | 1 | 12.407 | 30 | 1 | 2 | 46 | HAS ORTHOLOG WITH LARGER LLPS PROPENSITY |
| A0A1P8UJP1 | 1765967 | 1236 | 0.868867 | 1 | NaN | 0 | 0 | 1 | 50 | |
| A0A1N6GQR7 | 364032 | 1236 | 1.34245 | 1 | 13.692 | 43 | 1 | 2 | 85 | |
| A0A0F6TPU7 | 914150 | 1236 | 1.40871 | 1 | 13.104 | 38 | 1 | 2 | 43 | |
| D5V9T9 | 1236608 | 1236 | 1.40923 | 1 | 24.842 | 118 | 1 | 2 | 130 | |
| U2FYN2 | 1033802 | 1236 | 2.87398 | 1 | 17.819 | 49 | 1 | 2 | 74 | |
| A0A251XB99 | 1570016 | 1236 | 0.401477 | 0 | 13.183 | 42 | 1 | 1 | 74 | |
| A0A495DDK7 | 1304900 | 1236 | 0.928263 | 1 | 11.552 | 34 | 1 | 2 | 56 | |
| I3BZ36 | 870187 | 1236 | 2.04827 | 1 | NaN | 0 | 0 | 1 | 49 | |
| A0A432ZKZ7 | 337250 | 1236 | 1.51863 | 1 | 24.212 | 50 | 1 | 2 | 70 | |
| A0A451CMM4 | 2126341 | 1236 | -0.132187 | 0 | NaN | 0 | 0 | 0 | 95 | |
| E1VGN3 | 83406 | 1236 | 1.41743 | 1 | NaN | 0 | 0 | 1 | 70 | |
| A0A2P1PZT9 | 2021234 | 1236 | 3.3858 | 1 | NaN | 0 | 0 | 1 | 84 | HAS ORTHOLOG WITH LARGER LLPS PROPENSITY |
| A0A2P1PPH4 | 2021234 | 1236 | 3.61064 | 1 | NaN | 0 | 0 | 1 | 84 | |
| A0A063Y1S7 | 267850 | 1236 | 1.02539 | 1 | 17.161 | 50 | 1 | 2 | 68 | |
| A0A0C5W171 | 1445510 | 1236 | 1.20735 | 1 | 19.166 | 74 | 1 | 2 | 95 | |
| S6H895 | 1330036 | 1236 | 1.52704 | 1 | 27.477 | 82 | 1 | 2 | 107 | |
| A0A1E3GTP9 | 291169 | 1236 | 1.47223 | 1 | 12.267 | 45 | 1 | 2 | 61 | |
| F3LC44 | 937772 | 1236 | 1.78547 | 1 | 25.922 | 74 | 1 | 2 | 81 | |
| P57610 | 107806 | 1236 | 0.919736 | 1 | NaN | 0 | 0 | 1 | 60 | |
| Q83EP4 | 227377 | 1236 | 1.23055 | 1 | NaN | 0 | 0 | 1 | 51 | |
| P0AGE0 | 83333 | 1236 | 1.88477 | 1 | 16.574 | 62 | 1 | 2 | 67 | |
| P44409 | 71421 | 1236 | 1.26931 | 1 | 16.548 | 50 | 1 | 2 | 62 | |
| P40947 | 208964 | 1236 | 0.771867 | 1 | NaN | 0 | 0 | 1 | 55 | |
| Q8EA81 | 211586 | 1236 | 0.827277 | 1 | 18.855 | 119 | 1 | 2 | 128 | |
| Q9KUW2 | 243277 | 1236 | 0.992151 | 1 | 16.763 | 61 | 1 | 2 | 65 | |
| Q8D254 | 36870 | 1236 | 1.27235 | 1 | NaN | 0 | 0 | 1 | 52 | |
| Q8P778 | 190485 | 1236 | 2.46685 | 1 | 14.643 | 49 | 1 | 2 | 66 | |
| H1BI13 | 457402 | 1239 | 0.265108 | 0 | NaN | 0 | 0 | 0 | 42 | |
| A0A162TG23 | 1765683 | 1239 | 0.0317071 | 0 | NaN | 0 | 0 | 0 | 51 | |
| A0A3E2STE1 | 1849041 | 1239 | 0.69509 | 1 | NaN | 0 | 0 | 1 | 51 | |
| A0A173Y6I4 | 853 | 1239 | 0.274942 | 0 | NaN | 0 | 0 | 0 | 58 | |
| A0A0B4S0X3 | 33033 | 1239 | 0.649106 | 1 | NaN | 0 | 0 | 1 | 42 | |
| A0A0R2HIL1 | 1410657 | 1239 | 0.758129 | 1 | 18.468 | 31 | 1 | 2 | 71 | |
| F9VMA2 | 1029718 | 1239 | 0.45344 | 0 | NaN | 0 | 0 | 0 | 43 | |
| R6BKT7 | 1262778 | 1239 | 0.333603 | 0 | NaN | 0 | 0 | 0 | 68 | |
| G2KTA2 | 714313 | 1239 | 1.7123 | 1 | 19.45 | 39 | 1 | 2 | 69 | |
| R6DCZ8 | 1263007 | 1239 | 1.63147 | 1 | 12.736 | 34 | 1 | 2 | 56 | |
| A0A1E5G2Y0 | 766136 | 1239 | 0.916779 | 1 | NaN | 0 | 0 | 1 | 51 | |
| W8TJI2 | 1286171 | 1239 | 0.912809 | 1 | NaN | 0 | 0 | 1 | 35 | |
| W8U9E7 | 1286171 | 1239 | 0.646721 | 1 | NaN | 0 | 0 | 1 | 40 | HAS ORTHOLOG WITH LARGER LLPS PROPENSITY |
| A0A176U568 | 1715004 | 1239 | 0.542011 | 1 | NaN | 0 | 0 | 1 | 50 | |
| R6Q6K2 | 1262815 | 1239 | 0.360687 | 0 | NaN | 0 | 0 | 0 | 41 | |
| A0A1E9AGZ0 | 1739304 | 1239 | 0.173875 | 0 | NaN | 0 | 0 | 0 | 43 | |
| R5EAX7 | 1263000 | 1239 | 1.2682 | 1 | 12.761 | 31 | 1 | 2 | 62 | |
| R6HDB5 | 1262819 | 1239 | 0.118252 | 0 | NaN | 0 | 0 | 0 | 52 | |
| H6LCF9 | 931626 | 1239 | 0.63344 | 1 | NaN | 0 | 0 | 1 | 45 | |
| Q2G112 | 93061 | 1239 | 1.39696 | 1 | 19.098 | 51 | 1 | 2 | 66 | |
| A0A151YWL9 | 1811386 | 1239 | 0.96028 | 1 | NaN | 0 | 0 | 1 | 53 | |
| H6LDD9 | 931626 | 1239 | 0.569471 | 1 | NaN | 0 | 0 | 1 | 38 | HAS ORTHOLOG WITH LARGER LLPS PROPENSITY |
| R7EVC4 | 1262703 | 1239 | -0.123643 | 0 | NaN | 0 | 0 | 0 | 47 | |
| G8U005 | 679936 | 1239 | 0.159238 | 0 | NaN | 0 | 0 | 0 | 28 | |
| G8TUY0 | 679936 | 1239 | -0.0376206 | 0 | NaN | 0 | 0 | 0 | 29 | HAS ORTHOLOG WITH LARGER LLPS PROPENSITY |
| A0A4P6ZJ31 | 1720083 | 1239 | 2.57978 | 1 | 22.383 | 90 | 1 | 2 | 107 | |
| A0A0K2SQQ5 | 1555112 | 1239 | -0.26519 | 0 | NaN | 0 | 0 | 0 | 32 | |
| H1HWU8 | 796942 | 1239 | 1.23506 | 1 | NaN | 0 | 0 | 1 | 54 | |
| A0A396QCQ7 | 2292273 | 1239 | -0.16128 | 0 | NaN | 0 | 0 | 0 | 37 | |
| R7BIL2 | 1262991 | 1239 | -0.0392051 | 0 | NaN | 0 | 0 | 0 | 44 | |
| A0A143WYD9 | 1780379 | 1239 | -0.12838 | 0 | NaN | 0 | 0 | 0 | 58 | |
| R7BHP7 | 1262991 | 1239 | 0.446481 | 0 | NaN | 0 | 0 | 0 | 46 | |
| H3NL41 | 883114 | 1239 | 0.769351 | 1 | NaN | 0 | 0 | 1 | 52 | |
| A0A1V2YLN2 | 1884656 | 1239 | 0.398422 | 0 | NaN | 0 | 0 | 0 | 59 | |
| R6AYP4 | 1262775 | 1239 | 1.12787 | 1 | 19.763 | 42 | 1 | 2 | 92 | |
| Q8DNH4 | 171101 | 1239 | -0.0965099 | 0 | NaN | 0 | 0 | 0 | 31 | |
| A0A161QCQ8 | 520767 | 1239 | 0.201057 | 0 | NaN | 0 | 0 | 0 | 49 | |
| Q3AG26 | 246194 | 1239 | 0.0157311 | 0 | NaN | 0 | 0 | 0 | 42 | HYPERTH |
| A0A1C0BWY8 | 1768196 | 1239 | 0.283747 | 0 | 13.678 | 32 | 1 | 1 | 55 | |
| R5CH32 | 1262993 | 1239 | 0.913005 | 1 | NaN | 0 | 0 | 1 | 66 | |
| Q2RM71 | 264732 | 1239 | 1.07873 | 1 | NaN | 0 | 0 | 1 | 53 | |
| R5C772 | 1262993 | 1239 | 0.9708 | 1 | NaN | 0 | 0 | 1 | 58 | |
| A0A1I5EJ74 | 398199 | 1239 | 0.937754 | 1 | NaN | 0 | 0 | 1 | 40 | |
| A0A1I4ZKA1 | 398199 | 1239 | 0.523225 | 1 | NaN | 0 | 0 | 1 | 36 | HAS ORTHOLOG WITH LARGER LLPS PROPENSITY |
| A0A419SGI5 | 66863 | 1239 | 0.997607 | 1 | NaN | 0 | 0 | 1 | 46 | |
| A0A1S1V4U8 | 39480 | 1239 | 1.13303 | 1 | NaN | 0 | 0 | 1 | 46 | |
| A0A1S1V927 | 39480 | 1239 | 1.25563 | 1 | NaN | 0 | 0 | 1 | 36 | |
| A0A1S1V4S9 | 39480 | 1239 | 0.724217 | 1 | NaN | 0 | 0 | 1 | 32 | HAS ORTHOLOG WITH LARGER LLPS PROPENSITY |
| A0A1U7M817 | 1123403 | 1239 | 0.712831 | 1 | NaN | 0 | 0 | 1 | 36 | |
| A0A136WI26 | 36847 | 1239 | 0.445468 | 0 | NaN | 0 | 0 | 0 | 42 | |
| A0A1W1ZCB4 | 371602 | 1239 | 1.64255 | 1 | 14.542 | 62 | 1 | 2 | 85 | |
| A0A087N3J7 | 1473546 | 1239 | 1.01737 | 1 | 24.853 | 52 | 1 | 2 | 85 | |
| A0A3S9T1H9 | 1323375 | 1239 | 0.0125456 | 0 | NaN | 0 | 0 | 0 | 30 | HAS ORTHOLOG WITH LARGER LLPS PROPENSITY |
| A0A090JPR1 | 1912856 | 1239 | 0.571643 | 1 | 15.075 | 39 | 1 | 2 | 56 | |
| A0A3Q9HPG4 | 1323375 | 1239 | 0.392345 | 0 | NaN | 0 | 0 | 0 | 31 | |
| K9ES07 | 883081 | 1239 | 1.18679 | 1 | 25.637 | 49 | 1 | 2 | 82 | |
| A0A0R3JXP1 | 908809 | 1239 | 0.0563574 | 0 | NaN | 0 | 0 | 0 | 36 | |
| R6XRZ0 | 1262797 | 1239 | 0.104465 | 0 | NaN | 0 | 0 | 0 | 29 | |
| D9QUL1 | 574087 | 1239 | 0.309606 | 0 | NaN | 0 | 0 | 0 | 27 | |
| V1CFG6 | 1321814 | 1239 | 1.0382 | 1 | NaN | 0 | 0 | 1 | 35 | |
| A0A1V4IFH4 | 1450648 | 1239 | 1.22519 | 1 | NaN | 0 | 0 | 1 | 49 | |
| A0A421BEL4 | 2315861 | 1239 | 1.09453 | 1 | NaN | 0 | 0 | 1 | 74 | |
| E8LE33 | 626939 | 1239 | 0.453325 | 0 | NaN | 0 | 0 | 0 | 42 | |
| D7CKA0 | 643648 | 1239 | 0.266431 | 0 | NaN | 0 | 0 | 0 | 48 | |
| Q5WAH4 | 66692 | 1239 | 1.66043 | 1 | 18.019 | 37 | 1 | 2 | 60 | |
| Q5WE28 | 66692 | 1239 | 1.46046 | 1 | NaN | 0 | 0 | 1 | 39 | HAS ORTHOLOG WITH LARGER LLPS PROPENSITY |
| A0A0X8D601 | 1450761 | 1239 | 0.684178 | 1 | NaN | 0 | 0 | 1 | 54 | |
| K8ZAX7 | 1234409 | 1239 | 0.322772 | 0 | 10.82 | 57 | 1 | 1 | 74 | |
| R6H9V2 | 1263004 | 1239 | 1.01948 | 1 | 13.559 | 36 | 1 | 2 | 68 | |
| E6SLI9 | 644966 | 1239 | 0.562414 | 1 | NaN | 0 | 0 | 1 | 38 | |
| C7HV65 | 655811 | 1239 | 1.2742 | 1 | 22.905 | 42 | 1 | 2 | 68 | |
| C0E890 | 537013 | 1239 | 0.627233 | 1 | NaN | 0 | 0 | 1 | 56 | |
| D2RN22 | 591001 | 1239 | 1.06127 | 1 | 15.492 | 32 | 1 | 2 | 47 | |
| F2JQT2 | 642492 | 1239 | 0.681205 | 1 | NaN | 0 | 0 | 1 | 32 | |
| F2JRY8 | 642492 | 1239 | 0.132448 | 0 | NaN | 0 | 0 | 0 | 48 | HAS ORTHOLOG WITH LARGER LLPS PROPENSITY |
| R5L041 | 1262878 | 1239 | 0.124738 | 0 | NaN | 0 | 0 | 0 | 53 | |
| A0A410Q030 | 2507162 | 1239 | 0.508311 | 1 | NaN | 0 | 0 | 1 | 47 | HAS ORTHOLOG WITH LARGER LLPS PROPENSITY |
| A0A410Q504 | 2507162 | 1239 | 0.637557 | 1 | NaN | 0 | 0 | 1 | 52 | |
| H3NIJ2 | 883113 | 1239 | 1.22941 | 1 | 20.905 | 44 | 1 | 2 | 74 | |
| A0A1V4I639 | 29349 | 1239 | 0.827097 | 1 | NaN | 0 | 0 | 1 | 41 | |
| R6DH82 | 1262782 | 1239 | 0.790212 | 1 | NaN | 0 | 0 | 1 | 63 | |
| A0A1H9NC94 | 137733 | 1239 | 0.987316 | 1 | 24.055 | 44 | 1 | 2 | 79 | |
| R6DN65 | 1262782 | 1239 | 0.552351 | 1 | NaN | 0 | 0 | 1 | 48 | HAS ORTHOLOG WITH LARGER LLPS PROPENSITY |

| | | | | | | | | | | |
|---|---|---|---|---|---|---|---|---|---|---|
| C0GI94 | 555088 | 1239 | 0.0885872 | 0 | NaN | 0 | 0 | 0 | 28 | |
| Q67J49 | 292459 | 1239 | 0.851584 | 1 | NaN | 0 | 0 | 1 | 38 | |
| A0A1S6IRQ2 | 708126 | 1239 | 0.853871 | 1 | 23.777 | 40 | 1 | 2 | 75 | |
| A0A1G9WGR7 | 258515 | 1239 | -0.175998 | 0 | NaN | 0 | 0 | 0 | 47 | |
| D3QYV9 | 699246 | 1239 | 0.121471 | 0 | NaN | 0 | 0 | 0 | 72 | |
| A0A1M5XGN9 | 1121316 | 1239 | 0.967876 | 1 | 13.49 | 38 | 1 | 2 | 56 | |
| A0A292YIZ2 | 1348429 | 1239 | 2.15176 | 1 | NaN | 0 | 0 | 1 | 62 | |
| A0A095XHP9 | 1230734 | 1239 | 1.42785 | 1 | 10.655 | 35 | 1 | 2 | 71 | |
| A0A095XHU2 | 1230734 | 1239 | 0.492988 | 0 | NaN | 0 | 0 | 0 | 37 | HAS ORTHOLOG WITH LARGER LLPS PROPENSITY |
| A0A2N6S3P5 | 2069309 | 1239 | 1.00997 | 1 | 26.756 | 49 | 1 | 2 | 89 | HAS ORTHOLOG WITH LARGER LLPS PROPENSITY |
| A0A2N6S6D3 | 2069309 | 1239 | 1.626 | 1 | 26.199 | 83 | 1 | 2 | 105 | |
| C8WV95 | 521098 | 1239 | 0.0756005 | 0 | NaN | 0 | 0 | 0 | 55 | HAS ORTHOLOG WITH LARGER LLPS PROPENSITY |
| A0A0B4RFI6 | 1526927 | 1239 | 0.613348 | 1 | NaN | 0 | 0 | 1 | 58 | |
| R5QDS7 | 1263108 | 1239 | 0.261269 | 0 | NaN | 0 | 0 | 0 | 46 | |
| C8WYC0 | 521098 | 1239 | 0.341999 | 0 | NaN | 0 | 0 | 0 | 55 | |
| A0A1M6I0R6 | 1121476 | 1239 | 0.0104874 | 0 | NaN | 0 | 0 | 0 | 43 | |
| R5D149 | 1263030 | 1239 | 1.00922 | 1 | NaN | 0 | 0 | 1 | 48 | |
| A0A3A9HTQ7 | 2320083 | 1239 | 0.949483 | 1 | NaN | 0 | 0 | 1 | 64 | |
| A0A3A9H9M7 | 2320083 | 1239 | 0.881429 | 1 | 20.6 | 49 | 1 | 2 | 78 | HAS ORTHOLOG WITH LARGER LLPS PROPENSITY |
| M1ZL23 | 1288971 | 1239 | 0.278258 | 0 | NaN | 0 | 0 | 0 | 52 | |
| A0A4Q1ZU28 | 2283630 | 1239 | 0.666683 | 1 | NaN | 0 | 0 | 1 | 45 | |
| B1C975 | 445971 | 1239 | 0.675635 | 1 | NaN | 0 | 0 | 1 | 39 | |
| A0A1I5UUE9 | 937334 | 1239 | 0.125848 | 0 | NaN | 0 | 0 | 0 | 45 | |
| Q04HQ6 | 203123 | 1239 | 0.983052 | 1 | 16.255 | 55 | 1 | 2 | 84 | |
| A0A1M5UH46 | 1123282 | 1239 | 1.18122 | 1 | NaN | 0 | 0 | 1 | 52 | |
| A0A252F3M8 | 1945634 | 1239 | 0.493249 | 0 | NaN | 0 | 0 | 0 | 59 | |
| C7GYI5 | 592031 | 1239 | 0.418221 | 0 | NaN | 0 | 0 | 0 | 49 | |
| A0A096BVW1 | 1401067 | 1239 | 0.664772 | 1 | NaN | 0 | 0 | 1 | 30 | |
| A0A1Q9JTH7 | 1261636 | 1239 | 0.0431093 | 0 | NaN | 0 | 0 | 0 | 46 | |
| U2JRQ6 | 1321784 | 1239 | 1.09021 | 1 | 13.666 | 31 | 1 | 2 | 52 | |
| A0A143ZVI4 | 1780381 | 1239 | -0.520334 | 0 | NaN | 0 | 0 | 0 | 47 | |
| A0A4R3MQE2 | 682400 | 1239 | 0.455694 | 0 | NaN | 0 | 0 | 0 | 15 | MOTIF MISSING, FRAGMENT? |
| A0A1T4PF66 | 1121911 | 1239 | 0.529106 | 1 | NaN | 0 | 0 | 1 | 41 | HAS ORTHOLOG WITH LARGER LLPS PROPENSITY |
| A0A1T4K798 | 1121911 | 1239 | 0.905262 | 1 | NaN | 0 | 0 | 1 | 34 | |
| F0T2X6 | 645991 | 1239 | 0.220545 | 0 | NaN | 0 | 0 | 0 | 33 | |
| A3DF53 | 203119 | 1239 | 0.608088 | 1 | NaN | 0 | 0 | 1 | 34 | |
| A0A0J8DBU2 | 1121307 | 1239 | 0.993245 | 1 | NaN | 0 | 0 | 1 | 51 | |
| A0A0J8DFM9 | 1121307 | 1239 | 0.625599 | 1 | NaN | 0 | 0 | 1 | 34 | HAS ORTHOLOG WITH LARGER LLPS PROPENSITY |
| A3DHF9 | 203119 | 1239 | -0.224237 | 0 | NaN | 0 | 0 | 0 | 37 | HAS ORTHOLOG WITH LARGER LLPS PROPENSITY |
| A0A0X1TJI9 | 1712675 | 1239 | 1.7699 | 1 | 10.762 | 59 | 1 | 2 | 87 | |
| A0A0X8G3H0 | 1712675 | 1239 | 0.335972 | 0 | NaN | 0 | 0 | 0 | 78 | HAS ORTHOLOG WITH LARGER LLPS PROPENSITY |
| A0A0C2UP49 | 1590651 | 1239 | 0.317561 | 0 | NaN | 0 | 0 | 0 | 47 | |
| A0A1H7YGW9 | 474960 | 1239 | 0.232094 | 0 | NaN | 0 | 0 | 0 | 49 | HAS ORTHOLOG WITH LARGER LLPS PROPENSITY |
| A0A1H7ZUU3 | 474960 | 1239 | 0.418809 | 0 | NaN | 0 | 0 | 0 | 49 | |
| B2A455 | 457570 | 1239 | 1.55695 | 1 | NaN | 0 | 0 | 1 | 46 | |
| R6RG83 | 1262886 | 1239 | -0.0585626 | 0 | NaN | 0 | 0 | 0 | 42 | |
| E0E595 | 596315 | 1239 | 0.616451 | 1 | NaN | 0 | 0 | 1 | 46 | |
| E0E1I1 | 596315 | 1239 | 0.0439097 | 0 | NaN | 0 | 0 | 0 | 31 | HAS ORTHOLOG WITH LARGER LLPS PROPENSITY |
| C6LLT2 | 478749 | 1239 | 0.811432 | 1 | NaN | 0 | 0 | 1 | 53 | |
| A0A078KJD7 | 29343 | 1239 | 0.289301 | 0 | NaN | 0 | 0 | 0 | 48 | |
| S0J676 | 1235790 | 1239 | 0.60255 | 1 | NaN | 0 | 0 | 1 | 45 | |
| A0A1M6HXV8 | 1122184 | 1239 | 0.175476 | 0 | NaN | 0 | 0 | 0 | 35 | |
| R6W1H8 | 1262957 | 1239 | 0.350085 | 0 | NaN | 0 | 0 | 0 | 42 | |
| R6M3F2 | 1263021 | 1239 | 1.50688 | 1 | 18.187 | 30 | 1 | 2 | 62 | |
| A0A1I0FAR6 | 1526 | 1239 | 0.263099 | 0 | NaN | 0 | 0 | 0 | 53 | |
| A0A2K9NZD6 | 1981510 | 1239 | 0.962747 | 1 | 17.633 | 39 | 1 | 2 | 67 | |
| A0A2K9P3S1 | 1981510 | 1239 | 0.670228 | 1 | NaN | 0 | 0 | 1 | 43 | HAS ORTHOLOG WITH LARGER LLPS PROPENSITY |
| R5VM87 | 1262777 | 1239 | 0.239886 | 0 | NaN | 0 | 0 | 0 | 46 | |
| H1PJB0 | 883109 | 1239 | 0.069001 | 0 | NaN | 0 | 0 | 0 | 47 | |
| U2CZK6 | 1507 | 1239 | -0.0959872 | 0 | NaN | 0 | 0 | 0 | 49 | |
| A1HSI9 | 401526 | 1239 | 0.20269 | 0 | NaN | 0 | 0 | 0 | 36 | |
| A1HR42 | 401526 | 1239 | 0.103779 | 0 | NaN | 0 | 0 | 0 | 31 | HAS ORTHOLOG WITH LARGER LLPS PROPENSITY |
| U2DDE1 | 1507 | 1239 | -0.244738 | 0 | NaN | 0 | 0 | 0 | 30 | HAS ORTHOLOG WITH LARGER LLPS PROPENSITY |
| R6SGE0 | 1262808 | 1239 | -0.0335694 | 0 | NaN | 0 | 0 | 0 | 47 | |
| A0A2I0N1C9 | 1658742 | 1239 | 1.17401 | 1 | NaN | 0 | 0 | 1 | 43 | |
| G9WKY9 | 796943 | 1239 | 1.13072 | 1 | NaN | 0 | 0 | 1 | 48 | |
| G9WL70 | 796943 | 1239 | 0.647718 | 1 | NaN | 0 | 0 | 1 | 22 | HAS ORTHOLOG WITH LARGER LLPS PROPENSITY |
| R5R0I9 | 1263008 | 1239 | 0.958745 | 1 | NaN | 0 | 0 | 1 | 44 | |
| R7NN10 | 1262890 | 1239 | 0.980896 | 1 | 28.973 | 45 | 1 | 2 | 68 | |
| D5WXD0 | 562970 | 1239 | 0.401771 | 0 | NaN | 0 | 0 | 0 | 53 | |
| F7V2Z0 | 1042156 | 1239 | 0.586819 | 1 | NaN | 0 | 0 | 1 | 48 | |
| F6B6R2 | 868595 | 1239 | 1.39374 | 1 | NaN | 0 | 0 | 1 | 39 | |
| A0A0B0HJ06 | 1548750 | 1239 | 1.17414 | 1 | NaN | 0 | 0 | 1 | 62 | |
| A0A239U257 | 158847 | 1239 | -0.0832455 | 0 | NaN | 0 | 0 | 0 | 45 | |
| F2BXW4 | 888062 | 1239 | 0.338945 | 0 | NaN | 0 | 0 | 0 | 38 | |
| F9MQX5 | 1000569 | 1239 | 0.47301 | 0 | NaN | 0 | 0 | 0 | 41 | |
| A0A1I3NFR9 | 1520815 | 1239 | 0.902861 | 1 | NaN | 0 | 0 | 1 | 48 | |
| R6I6X7 | 1262952 | 1239 | 0.012856 | 0 | NaN | 0 | 0 | 0 | 46 | |
| R6R1Y2 | 1262761 | 1239 | 0.647276 | 1 | NaN | 0 | 0 | 1 | 43 | |
| A0A0K8J2K5 | 1679721 | 1239 | 0.559963 | 1 | NaN | 0 | 0 | 1 | 46 | |
| R5ZLD1 | 1263028 | 1239 | -0.214044 | 0 | 16.71 | 32 | 1 | 1 | 62 | |
| R5B767 | 1262781 | 1239 | 0.275775 | 0 | NaN | 0 | 0 | 0 | 41 | |
| A0A0R1HN48 | 1423719 | 1239 | 0.663939 | 1 | 19.271 | 61 | 1 | 2 | 76 | |
| B0S2G8 | 334413 | 1239 | 1.1614 | 1 | 25.434 | 35 | 1 | 2 | 63 | |
| H1BPB1 | 457402 | 1239 | 0.110166 | 0 | NaN | 0 | 0 | 0 | 55 | |
| B0S470 | 334413 | 1239 | 0.385484 | 0 | NaN | 0 | 0 | 0 | 48 | HAS ORTHOLOG WITH LARGER LLPS PROPENSITY |
| F3ZYJ8 | 697281 | 1239 | 0.217556 | 0 | NaN | 0 | 0 | 0 | 51 | |
| F3ZWZ1 | 697281 | 1239 | -0.317823 | 0 | NaN | 0 | 0 | 0 | 36 | HAS ORTHOLOG WITH LARGER LLPS PROPENSITY |
| B0S487 | 334413 | 1239 | 0.498836 | 0 | NaN | 0 | 0 | 0 | 48 | HAS ORTHOLOG WITH LARGER LLPS PROPENSITY |
| B0S4C2 | 334413 | 1239 | 0.791535 | 1 | NaN | 0 | 0 | 1 | 36 | |
| A0A410PS30 | 2507160 | 1239 | 0.775085 | 1 | NaN | 0 | 0 | 1 | 44 | |
| A0A1M6M3R1 | 1120989 | 1239 | 0.203573 | 0 | NaN | 0 | 0 | 0 | 27 | MOTIF MISSING, FRAGMENT? |
| A0A1G9JEA3 | 1798184 | 1239 | 2.8682 | 1 | 22.393 | 70 | 1 | 2 | 100 | |
| A0A3P1S1N3 | 2491058 | 1239 | 0.622724 | 1 | NaN | 0 | 0 | 1 | 51 | |
| D5XDR1 | 635013 | 1239 | 0.545327 | 1 | NaN | 0 | 0 | 1 | 32 | |
| B9Y3H0 | 545696 | 1239 | 0.546078 | 1 | NaN | 0 | 0 | 1 | 57 | |
| A0A0M2NCL8 | 270498 | 1239 | 0.16373 | 0 | NaN | 0 | 0 | 0 | 49 | |
| A0A1B6BDQ2 | 1048380 | 1239 | 0.534905 | 1 | NaN | 0 | 0 | 1 | 38 | |
| A0A3P7PXA5 | 2173034 | 1239 | 0.79557 | 1 | NaN | 0 | 0 | 1 | 47 | |
| A0A133YGY4 | 1497955 | 1239 | 0.476293 | 0 | NaN | 0 | 0 | 0 | 60 | |
| V6Q3E5 | 1408226 | 1239 | 0.503965 | 1 | NaN | 0 | 0 | 1 | 66 | |
| R5AH99 | 1262770 | 1239 | 0.322789 | 0 | NaN | 0 | 0 | 0 | 39 | |
| A0A1G9Q5V4 | 321763 | 1239 | 0.140126 | 0 | NaN | 0 | 0 | 0 | 49 | |
| A0A1G9R8I9 | 321763 | 1239 | -0.0318378 | 0 | NaN | 0 | 0 | 0 | 48 | HAS ORTHOLOG WITH LARGER LLPS PROPENSITY |
| G9RVU9 | 665956 | 1239 | 0.283028 | 0 | NaN | 0 | 0 | 0 | 55 | |
| R7A8H1 | 1262805 | 1239 | 1.10913 | 1 | NaN | 0 | 0 | 1 | 47 | |
| R5VRL4 | 1262953 | 1239 | 0.0308087 | 0 | 19.02 | 47 | 1 | 1 | 77 | HAS ORTHOLOG WITH LARGER LLPS PROPENSITY |
| A0A425W3H2 | 2049045 | 1239 | 0.669917 | 1 | NaN | 0 | 0 | 1 | 46 | |
| R5VJF6 | 1262953 | 1239 | 0.163926 | 0 | 11.781 | 44 | 1 | 1 | 89 | |
| K0B4K0 | 1128398 | 1239 | 0.772243 | 1 | NaN | 0 | 0 | 1 | 40 | |
| A0A2L2XIB0 | 1898651 | 1239 | 0.388196 | 0 | NaN | 0 | 0 | 0 | 32 | HAS ORTHOLOG WITH LARGER LLPS PROPENSITY |
| A0A2L2X8Z9 | 1898651 | 1239 | 0.491567 | 0 | NaN | 0 | 0 | 0 | 35 | |
| A0A0R2FS85 | 1123500 | 1239 | 1.23171 | 1 | 12.145 | 58 | 1 | 2 | 82 | |
| R7GGC3 | 1262795 | 1239 | 0.338634 | 0 | 29.038 | 42 | 1 | 1 | 74 | |
| K4LED7 | 1089553 | 1239 | 0.216837 | 0 | NaN | 0 | 0 | 0 | 47 | |
| E7MR23 | 706433 | 1239 | 0.428724 | 0 | NaN | 0 | 0 | 0 | 57 | |
| A0A1M4T590 | 1120975 | 1239 | 1.23161 | 1 | NaN | 0 | 0 | 1 | 48 | |
| V2XNR2 | 592026 | 1239 | 1.04689 | 1 | NaN | 0 | 0 | 1 | 47 | |
| A0A0D0SB35 | 1392836 | 1239 | 0.391447 | 0 | NaN | 0 | 0 | 0 | 51 | |
| A0A0N8NST2 | 36849 | 1239 | 1.62876 | 1 | NaN | 0 | 0 | 1 | 42 | |
| A0A0P8WVA5 | 36849 | 1239 | 0.55075 | 1 | NaN | 0 | 0 | 1 | 32 | HAS ORTHOLOG WITH LARGER LLPS PROPENSITY |
| A0A150LKD9 | 301148 | 1239 | 0.405414 | 0 | NaN | 0 | 0 | 0 | 57 | |
| A0A1G6VZD6 | 2741 | 1239 | 0.00356113 | 0 | NaN | 0 | 0 | 0 | 43 | HAS ORTHOLOG WITH LARGER LLPS PROPENSITY |
| A0A1G7AEL3 | 2741 | 1239 | 0.862284 | 1 | NaN | 0 | 0 | 1 | 49 | |
| R6D7W9 | 1262963 | 1239 | 0.888404 | 1 | 26.233 | 43 | 1 | 2 | 71 | |
| R6TSJ9 | 1263015 | 1239 | 0.0117942 | 0 | NaN | 0 | 0 | 0 | 50 | |
| A0A0U1KIL9 | 1462524 | 1239 | 1.91295 | 1 | 19.07 | 35 | 1 | 2 | 61 | |
| E3DS65 | 572479 | 1239 | 1.55019 | 1 | 27.773 | 30 | 1 | 2 | 58 | |
| A0A3P2AI34 | 2491044 | 1239 | 0.636577 | 1 | NaN | 0 | 0 | 1 | 46 | |

| | | | | | | | | | | |
|---|---|---|---|---|---|---|---|---|---|---|
| D6GT00 | 546269 | 1239 | 1.03492 | 1 | NaN | 0 | 0 | 1 | 68 | |
| E7FWX8 | 525280 | 1239 | 0.184656 | 0 | NaN | 0 | 0 | 0 | 59 | |
| L7VKB8 | 1121335 | 1239 | 0.379538 | 0 | NaN | 0 | 0 | 0 | 49 | |
| A0A0R2DN02 | 1423744 | 1239 | 0.743002 | 1 | 20.291 | 32 | 1 | 2 | 67 | |
| A0A4Q7P0A5 | 1796618 | 1239 | 0.195977 | 0 | NaN | 0 | 0 | 0 | 4 | MOTIF MISSING, FRAGMENT? |
| A0A1M6B6J6 | 1122934 | 1239 | 1.00527 | 1 | NaN | 0 | 0 | 1 | 41 | |
| A0A1M6LNN8 | 1122934 | 1239 | 0.647505 | 1 | NaN | 0 | 0 | 1 | 44 | HAS ORTHOLOG WITH LARGER LLPS PROPENSITY |
| A0A1H3EPH8 | 1520816 | 1239 | 0.439734 | 0 | NaN | 0 | 0 | 0 | 62 | |
| B8D1C8 | 373903 | 1239 | 0.294904 | 0 | NaN | 0 | 0 | 0 | 33 | |
| C4GAT4 | 626523 | 1239 | 0.729068 | 1 | NaN | 0 | 0 | 1 | 40 | HAS ORTHOLOG WITH LARGER LLPS PROPENSITY |
| C4GC71 | 626523 | 1239 | 1.20503 | 1 | NaN | 0 | 0 | 1 | 48 | |
| E4Q4V6 | 632518 | 1239 | -0.249132 | 0 | NaN | 0 | 0 | 0 | 62 | |
| B1I6T0 | 477974 | 1239 | 0.928541 | 1 | NaN | 0 | 0 | 1 | 41 | |
| T0ND85 | 1354301 | 1239 | 1.08801 | 1 | NaN | 0 | 0 | 1 | 46 | |
| K0J855 | 698758 | 1239 | 0.996594 | 1 | 17.839 | 40 | 1 | 2 | 62 | |
| A0A448V2H2 | 54006 | 1239 | 1.21899 | 1 | NaN | 0 | 0 | 1 | 55 | |
| A0A0U1QL81 | 1069536 | 1239 | 1.11168 | 1 | 15.053 | 42 | 1 | 2 | 71 | |
| R6PGX1 | 1262888 | 1239 | 0.647505 | 1 | 12.947 | 30 | 1 | 2 | 51 | |
| A0A4V2ZP92 | 2528593 | 1239 | 0.599871 | 1 | NaN | 0 | 0 | 1 | 36 | |
| A0A1T4PN32 | 42322 | 1239 | 1.16491 | 1 | NaN | 0 | 0 | 1 | 43 | |
| A0A1T4P1L0 | 42322 | 1239 | 0.516527 | 1 | NaN | 0 | 0 | 1 | 36 | HAS ORTHOLOG WITH LARGER LLPS PROPENSITY |
| A0A417HJH0 | 2292269 | 1239 | -0.0127907 | 0 | NaN | 0 | 0 | 0 | 41 | |
| R5LYQ9 | 1263062 | 1239 | 0.820122 | 1 | NaN | 0 | 0 | 1 | 43 | |
| A0A1Y4M5T6 | 1965572 | 1239 | 0.662681 | 1 | NaN | 0 | 0 | 1 | 46 | |
| A0A1W1V2Y9 | 656914 | 1239 | 0.972989 | 1 | NaN | 0 | 0 | 1 | 39 | |
| R5HBE8 | 1263012 | 1239 | 1.12132 | 1 | NaN | 0 | 0 | 1 | 50 | |
| A0A1W1VQW5 | 656914 | 1239 | 0.873278 | 1 | NaN | 0 | 0 | 1 | 31 | HAS ORTHOLOG WITH LARGER LLPS PROPENSITY |
| A0A140L017 | 520762 | 1239 | 0.91593 | 1 | NaN | 0 | 0 | 1 | 35 | |
| B0TA59 | 498761 | 1239 | 2.20133 | 1 | 7.956 | 51 | 1 | 2 | 72 | |
| A0A1G8RAN8 | 1200749 | 1239 | 1.48993 | 1 | NaN | 0 | 0 | 1 | 44 | |
| A0A1U9K6Q2 | 1471761 | 1239 | 0.711671 | 1 | NaN | 0 | 0 | 1 | 37 | |
| B8I3W7 | 394503 | 1239 | 0.33852 | 0 | NaN | 0 | 0 | 0 | 42 | |
| A0A1G5RRY9 | 1120920 | 1239 | 0.240131 | 0 | NaN | 0 | 0 | 0 | 40 | |
| A0A1U9KB70 | 1471761 | 1239 | 0.0849444 | 0 | NaN | 0 | 0 | 0 | 39 | HAS ORTHOLOG WITH LARGER LLPS PROPENSITY |
| A0A143XW62 | 1780380 | 1239 | 0.341869 | 0 | NaN | 0 | 0 | 0 | 62 | |
| J4UBY6 | 936556 | 1239 | 0.590331 | 1 | NaN | 0 | 0 | 1 | 47 | |
| R5A556 | 1262769 | 1239 | 0.439685 | 0 | NaN | 0 | 0 | 0 | 60 | |
| A0A267MFW7 | 1478221 | 1239 | 1.54922 | 1 | NaN | 0 | 0 | 1 | 43 | |
| A0A1H3MBG9 | 159292 | 1239 | 0.558657 | 1 | NaN | 0 | 0 | 1 | 49 | |
| A0A3A9DZA7 | 2320089 | 1239 | 0.0399565 | 0 | 9.073 | 52 | 1 | 1 | 74 | |
| A0A2V3VUB7 | 1494959 | 1239 | 0.609574 | 1 | 16.28 | 42 | 1 | 2 | 61 | |
| A0A1T4VR86 | 39495 | 1239 | 1.03359 | 1 | NaN | 0 | 0 | 1 | 49 | |
| A0A151AKW6 | 1121305 | 1239 | 0.69019 | 1 | NaN | 0 | 0 | 1 | 52 | |
| R7I585 | 1262897 | 1239 | -0.238057 | 0 | NaN | 0 | 0 | 0 | 43 | |
| A0A1M6MPR0 | 1121331 | 1239 | 0.976991 | 1 | NaN | 0 | 0 | 1 | 49 | |
| F3BBR7 | 742723 | 1239 | 0.371027 | 0 | NaN | 0 | 0 | 0 | 43 | |
| A0A1M6S2I4 | 1121331 | 1239 | 0.315127 | 0 | NaN | 0 | 0 | 0 | 32 | HAS ORTHOLOG WITH LARGER LLPS PROPENSITY |
| Q8YAR8 | 169963 | 1239 | 2.02815 | 1 | 24.43 | 53 | 1 | 2 | 77 | |
| Q8Y4X1 | 169963 | 1239 | 0.71845 | 1 | NaN | 0 | 0 | 1 | 59 | HAS ORTHOLOG WITH LARGER LLPS PROPENSITY |
| P37455 | 224308 | 1239 | 2.42273 | 1 | 19.368 | 46 | 1 | 2 | 71 | |
| Q9K5N9 | 272558 | 1239 | 1.9761 | 1 | 13.16 | 40 | 1 | 2 | 67 | |
| Q8R6M2 | 273068 | 1239 | -0.130439 | 0 | NaN | 0 | 0 | 0 | 49 | HYPERTH |
| Q8XH44 | 195102 | 1239 | 1.62935 | 1 | NaN | 0 | 0 | 1 | 53 | |
| P66855 | 171101 | 1239 | 0.174348 | 0 | NaN | 0 | 0 | 0 | 55 | |
| D7CWQ9 | 649638 | 1297 | 1.05372 | 1 | NaN | 0 | 0 | 1 | 50 | |
| Q9RY51 | 243230 | 1297 | 1.25969 | 1 | NaN | 0 | 0 | 1 | 72 | |
| Q5SLP9 | 300852 | 1297 | 0.761821 | 1 | NaN | 0 | 0 | 1 | 39 | HYPERTH |
| A0A2P2EDN8 | 1445552 | 28211 | 0.499228 | 0 | NaN | 0 | 0 | 0 | 53 | |
| Q5NM94 | 264203 | 28211 | 3.29214 | 1 | 14.453 | 37 | 1 | 2 | 81 | |
| A0A0H5BEK0 | 1079 | 28211 | 1.53276 | 1 | NaN | 0 | 0 | 1 | 52 | |
| A0A2Z4UHZ3 | 2026785 | 28211 | 2.53915 | 1 | NaN | 0 | 0 | 1 | 47 | |
| A0A397QB03 | 933063 | 28211 | 2.06947 | 1 | NaN | 0 | 0 | 1 | 53 | |
| D5BQS9 | 488538 | 28211 | 2.18488 | 1 | NaN | 0 | 0 | 1 | 56 | |
| A0A061QEU3 | 1492281 | 28211 | 2.45099 | 1 | NaN | 0 | 0 | 1 | 53 | |
| A0A1G7NI54 | 1082479 | 28211 | 1.70083 | 1 | NaN | 0 | 0 | 1 | 48 | |
| A0A2W2BAD0 | 2219703 | 28211 | 2.22829 | 1 | NaN | 0 | 0 | 1 | 53 | |
| A0A1S8D9X7 | 207340 | 28211 | 4.05748 | 1 | NaN | 0 | 0 | 1 | 66 | |
| A0A2Z6HZP9 | 1885025 | 28211 | 3.59699 | 1 | NaN | 0 | 0 | 1 | 78 | |
| A0A077FMS8 | 1528098 | 28211 | 0.841603 | 1 | NaN | 0 | 0 | 1 | 42 | |
| A0A239PTK7 | 1519374 | 28211 | 1.75688 | 1 | NaN | 0 | 0 | 1 | 38 | |
| A0A1H8Y143 | 1380357 | 28211 | 2.56656 | 1 | NaN | 0 | 0 | 1 | 55 | |
| K2JQQ4 | 1207063 | 28211 | 3.28929 | 1 | NaN | 0 | 0 | 1 | 58 | |
| D5AUB1 | 272942 | 28211 | 3.37452 | 1 | 9.28 | 47 | 1 | 2 | 63 | |
| A0A2P2E8R9 | 1445552 | 28211 | 2.19364 | 1 | NaN | 0 | 0 | 1 | 56 | |
| A0A0C1QWE1 | 86105 | 28211 | 0.886983 | 1 | NaN | 0 | 0 | 1 | 50 | |
| W6K6N2 | 1288970 | 28211 | 3.30489 | 1 | 17.808 | 33 | 1 | 2 | 62 | |
| A0A0D8CEC3 | 1616823 | 28211 | 3.51518 | 1 | NaN | 0 | 0 | 1 | 64 | |
| Q6N604 | 258594 | 28211 | 2.42797 | 1 | NaN | 0 | 0 | 1 | 55 | |
| A0A420WKS8 | 568099 | 28211 | 2.7257 | 1 | NaN | 0 | 0 | 1 | 55 | |
| A0A1G7CQK7 | 637679 | 28211 | 3.33848 | 1 | 17.846 | 51 | 1 | 2 | 67 | |
| A0LDV7 | 156889 | 28211 | 2.11801 | 1 | 9.565 | 45 | 1 | 2 | 70 | |
| G2KQL9 | 856793 | 28211 | 1.81023 | 1 | NaN | 0 | 0 | 1 | 49 | |
| A0A1B1ADN2 | 1759059 | 28211 | 1.92549 | 1 | NaN | 0 | 0 | 1 | 48 | |
| A0A318MZ43 | 2070537 | 28211 | 0.667974 | 1 | NaN | 0 | 0 | 1 | 59 | |
| A0A2S9QCH8 | 346911 | 28211 | 1.41687 | 1 | NaN | 0 | 0 | 1 | 50 | |
| A0A366FSW9 | 1473586 | 28211 | 2.13568 | 1 | NaN | 0 | 0 | 1 | 59 | |
| A0A366EH65 | 1473586 | 28211 | 0.564456 | 1 | NaN | 0 | 0 | 1 | 47 | HAS ORTHOLOG WITH LARGER LLPS PROPENSITY |
| A0A2G4YV69 | 2043170 | 28211 | 2.8139 | 1 | NaN | 0 | 0 | 1 | 49 | |
| E0TIG3 | 314260 | 28211 | 2.23454 | 1 | NaN | 0 | 0 | 1 | 42 | |
| A0A143DFC1 | 1549855 | 28211 | 1.55958 | 1 | NaN | 0 | 0 | 1 | 66 | |
| J9DXI6 | 1220535 | 28211 | 1.98409 | 1 | NaN | 0 | 0 | 1 | 44 | |
| I3TKW7 | 1110502 | 28211 | 5.32348 | 1 | 11.699 | 73 | 1 | 2 | 101 | |
| A0A1Y5T5N0 | 745714 | 28211 | 3.34536 | 1 | NaN | 0 | 0 | 1 | 54 | |
| Q2GD12 | 222891 | 28211 | 0.370652 | 0 | NaN | 0 | 0 | 0 | 42 | |
| A0A1B2AGZ7 | 692370 | 28211 | 3.10282 | 1 | 7.891 | 47 | 1 | 2 | 59 | |
| A0A165YQ76 | 989403 | 28211 | 4.27589 | 1 | 14.179 | 64 | 1 | 2 | 77 | |
| A5G056 | 349163 | 28211 | 1.46859 | 1 | NaN | 0 | 0 | 1 | 49 | |
| A8ID59 | 438753 | 28211 | 4.80647 | 1 | 9.594 | 67 | 1 | 2 | 93 | |
| A0A3Q8T4R6 | 2486578 | 28211 | 0.956164 | 1 | NaN | 0 | 0 | 1 | 35 | |
| A0A3S4DSA3 | 2490941 | 28211 | 3.12035 | 1 | NaN | 0 | 0 | 1 | 62 | |
| C6XJ14 | 582402 | 28211 | 1.68159 | 1 | NaN | 0 | 0 | 1 | 46 | |
| B6ISP2 | 414684 | 28211 | 3.78289 | 1 | 11.25 | 62 | 1 | 2 | 76 | |
| B9KI69 | 320483 | 28211 | 0.382348 | 0 | NaN | 0 | 0 | 0 | 55 | |
| B8ET29 | 395965 | 28211 | 0.994601 | 1 | NaN | 0 | 0 | 1 | 56 | |
| A0A368A097 | 2079009 | 28211 | 1.29878 | 1 | NaN | 0 | 0 | 1 | 42 | |
| A0A317EHV3 | 2211142 | 28211 | 4.48613 | 1 | 9.37 | 44 | 1 | 2 | 67 | |
| A0A1Y2QVV4 | 1985171 | 28211 | 3.16349 | 1 | NaN | 0 | 0 | 1 | 62 | |
| A0A1M5GKK8 | 1122133 | 28211 | 1.30251 | 1 | NaN | 0 | 0 | 1 | 58 | |
| A0A1M4Y5J1 | 1122133 | 28211 | -0.000604412 | 0 | NaN | 0 | 0 | 0 | 41 | HAS ORTHOLOG WITH LARGER LLPS PROPENSITY |
| A0A3L7JB23 | 2448481 | 28211 | 2.93058 | 1 | NaN | 0 | 0 | 1 | 55 | |
| A0A255XNV0 | 2022747 | 28211 | 4.26102 | 1 | 9.122 | 60 | 1 | 2 | 79 | |
| C6XHB3 | 537021 | 28211 | 0.387265 | 0 | NaN | 0 | 0 | 0 | 51 | |
| D8JQM6 | 582899 | 28211 | 2.10349 | 1 | NaN | 0 | 0 | 1 | 61 | |
| F7XVZ1 | 696127 | 28211 | 0.39754 | 0 | NaN | 0 | 0 | 0 | 43 | |
| B2IFV1 | 395963 | 28211 | 1.61621 | 1 | NaN | 0 | 0 | 1 | 68 | |
| A0A3R9ZIV1 | 2492378 | 28211 | 0.859883 | 1 | NaN | 0 | 0 | 1 | 41 | |
| A0A1G7UJF9 | 83401 | 28211 | 4.4907 | 1 | 18.229 | 50 | 1 | 2 | 72 | |
| A5CE60 | 357244 | 28211 | 0.344662 | 0 | NaN | 0 | 0 | 0 | 43 | |
| A0A0M2RDE4 | 1549748 | 28211 | 3.12164 | 1 | NaN | 0 | 0 | 1 | 63 | |
| A0A1E2S2U4 | 1177755 | 28211 | 1.93356 | 1 | NaN | 0 | 0 | 1 | 48 | |
| Q0ANG1 | 394221 | 28211 | 2.5383 | 1 | NaN | 0 | 0 | 1 | 52 | |
| J9Z216 | 744985 | 28211 | 0.351278 | 0 | NaN | 0 | 0 | 0 | 43 | |
| Q4FLZ6 | 335992 | 28211 | 1.37659 | 1 | NaN | 0 | 0 | 1 | 39 | |
| A0A371WUQ3 | 2293567 | 28211 | 1.68218 | 1 | NaN | 0 | 0 | 1 | 62 | |
| Q9A894 | 190650 | 28211 | 2.25629 | 1 | NaN | 0 | 0 | 1 | 55 | |
| P56898 | 266834 | 28211 | 2.49147 | 1 | NaN | 0 | 0 | 1 | 62 | |
| Q9ZCC2 | 272947 | 28211 | 0.728398 | 1 | NaN | 0 | 0 | 1 | 40 | |
| M1MEH1 | 1208922 | 28216 | 0.38109 | 0 | NaN | 0 | 0 | 0 | 38 | |
| A0A142LMB2 | 1690485 | 28216 | 1.41281 | 1 | NaN | 0 | 0 | 1 | 56 | |

| | | | | | | | | | | |
|---|---|---|---|---|---|---|---|---|---|---|
| A0A0H4JBR5 | 1623450 | 28216 | 0.492906 | 0 | NaN | 0 | 0 | 0 | 52 | |
| Q3SLL6 | 292415 | 28216 | 1.28121 | 1 | NaN | 0 | 0 | 1 | 52 | |
| A0A433SCC3 | 1965230 | 28216 | 1.33789 | 1 | NaN | 0 | 0 | 1 | 59 | |
| A0A0D6EWQ0 | 1581557 | 28216 | 0.711017 | 1 | NaN | 0 | 0 | 1 | 57 | |
| E8UF32 | 937774 | 28216 | 1.88026 | 1 | NaN | 0 | 0 | 1 | 70 | |
| D5X2Y7 | 75379 | 28216 | 1.22011 | 1 | NaN | 0 | 0 | 1 | 60 | |
| E0TIQ6 | 871271 | 28216 | 1.63552 | 1 | NaN | 0 | 0 | 1 | 89 | |
| A0A0A1H9H4 | 1469502 | 28216 | 0.797742 | 1 | NaN | 0 | 0 | 1 | 53 | |
| A0A165FGM2 | 1452487 | 28216 | 0.909248 | 1 | NaN | 0 | 0 | 1 | 62 | |
| A0A149VSA8 | 1356306 | 28216 | 1.08383 | 1 | NaN | 0 | 0 | 1 | 47 | |
| A0A2I7N694 | 2052837 | 28216 | 0.937999 | 1 | NaN | 0 | 0 | 1 | 52 | |
| A4G9L8 | 204773 | 28216 | 0.745632 | 1 | NaN | 0 | 0 | 1 | 55 | |
| A0A0P0LWH6 | 1678129 | 28216 | 0.903057 | 1 | NaN | 0 | 0 | 1 | 57 | |
| A0A212TEE9 | 2049319 | 28216 | 0.589122 | 1 | NaN | 0 | 0 | 1 | 40 | |
| A0A011PF87 | 1454000 | 28216 | 1.31543 | 1 | NaN | 0 | 0 | 1 | 53 | |
| E7RYT1 | 887898 | 28216 | 3.17823 | 1 | NaN | 0 | 0 | 1 | 69 | |
| A6GPL2 | 391597 | 28216 | 1.60906 | 1 | NaN | 0 | 0 | 1 | 63 | |
| A0A345DA70 | 2268024 | 28216 | 0.403927 | 0 | NaN | 0 | 0 | 0 | 51 | |
| D9SK11 | 395494 | 28216 | 0.442854 | 0 | NaN | 0 | 0 | 0 | 56 | |
| A0A1I5CZX8 | 83765 | 28216 | 0.890528 | 1 | NaN | 0 | 0 | 1 | 53 | |
| A0A2Z6I720 | 2494234 | 28216 | 0.258035 | 0 | NaN | 0 | 0 | 0 | 64 | |
| A0A0L1KQ71 | 1090379 | 28216 | 1.31126 | 1 | NaN | 0 | 0 | 1 | 32 | |
| U5NBX9 | 946483 | 28216 | 0.382805 | 0 | NaN | 0 | 0 | 0 | 62 | |
| A0A0N1B841 | 1523428 | 28216 | 0.669542 | 1 | NaN | 0 | 0 | 1 | 51 | |
| P66846 | 257313 | 28216 | 0.889515 | 1 | NaN | 0 | 0 | 1 | 59 | |
| P66849 | 122586 | 28216 | 0.711066 | 1 | NaN | 0 | 0 | 1 | 70 | |
| Q82S98 | 228410 | 28216 | 0.853332 | 1 | NaN | 0 | 0 | 1 | 44 | |
| D1ANB3 | 526218 | 32066 | -0.136042 | 0 | NaN | 0 | 0 | 0 | 30 | HAS ORTHOLOG WITH LARGER LLPS PROPENSITY |
| A0A0E3UTZ2 | 187101 | 32066 | 1.00795 | 1 | NaN | 0 | 0 | 1 | 36 | |
| A0A0E3ZA78 | 187101 | 32066 | -0.181046 | 0 | NaN | 0 | 0 | 0 | 31 | HAS ORTHOLOG WITH LARGER LLPS PROPENSITY |
| E3H7R3 | 572544 | 32066 | 1.02848 | 1 | NaN | 0 | 0 | 1 | 51 | |
| R7LYF2 | 1262901 | 32066 | 0.413369 | 0 | NaN | 0 | 0 | 0 | 44 | |
| D1AJZ4 | 526218 | 32066 | 0.114953 | 0 | NaN | 0 | 0 | 0 | 41 | |
| C7NDG9 | 523794 | 32066 | 0.839904 | 1 | NaN | 0 | 0 | 1 | 50 | |
| Q8RE26 | 190304 | 32066 | 0.780133 | 1 | NaN | 0 | 0 | 1 | 53 | |
| A0A091FBC6 | 1499107 | 68525 | 0.78603 | 1 | NaN | 0 | 0 | 1 | 34 | |
| A6Q465 | 387092 | 68525 | 0.189638 | 0 | NaN | 0 | 0 | 0 | 47 | |
| A0A2L1GRE7 | 1986146 | 68525 | 2.37777 | 1 | NaN | 0 | 0 | 1 | 56 | |
| A0A0K1Q5A4 | 1391654 | 68525 | 4.18212 | 1 | NaN | 0 | 0 | 1 | 83 | |
| C8X289 | 485915 | 68525 | 1.24968 | 1 | 17.488 | 31 | 1 | 2 | 50 | |
| A0A292YD15 | 1936991 | 68525 | 0.26073 | 0 | NaN | 0 | 0 | 0 | 52 | |
| A0A1M7SZD8 | 1121455 | 68525 | 0.86101 | 1 | NaN | 0 | 0 | 1 | 64 | |
| A0A0K1PGB9 | 1391653 | 68525 | 6.07001 | 1 | 12.519 | 76 | 1 | 2 | 83 | |
| A0A1G5I4D1 | 419481 | 68525 | 3.02626 | 1 | 25.125 | 81 | 1 | 2 | 103 | |
| A0A127ALM2 | 1621989 | 68525 | 0.238612 | 0 | NaN | 0 | 0 | 0 | 27 | HAS ORTHOLOG WITH LARGER LLPS PROPENSITY |
| A0A127ANQ3 | 1621989 | 68525 | 0.318444 | 0 | NaN | 0 | 0 | 0 | 26 | |
| A0A2S9YBE1 | 215803 | 68525 | 5.78806 | 1 | 12.908 | 62 | 1 | 2 | 89 | |
| A0A1L3GPE4 | 1842532 | 68525 | 0.625567 | 1 | NaN | 0 | 0 | 1 | 45 | |
| A8ZRT7 | 96561 | 68525 | 1.37999 | 1 | NaN | 0 | 0 | 1 | 32 | |
| D6Z5C9 | 589865 | 68525 | 0.480818 | 0 | NaN | 0 | 0 | 0 | 32 | |
| A0A444IQ37 | 1859131 | 68525 | 2.88455 | 1 | 8.034 | 65 | 1 | 2 | 72 | |
| F2NJT4 | 880072 | 68525 | 0.286246 | 0 | NaN | 0 | 0 | 0 | 32 | |
| A0A0M9ECS0 | 1509431 | 68525 | 0.199995 | 0 | NaN | 0 | 0 | 0 | 45 | |
| Q6AK18 | 177439 | 68525 | 0.902796 | 1 | NaN | 0 | 0 | 1 | 39 | |
| A0A1I1X788 | 54 | 68525 | 5.79296 | 1 | 13.922 | 61 | 1 | 2 | 97 | |
| A0A0M9EAX6 | 1509431 | 68525 | 0.188593 | 0 | NaN | 0 | 0 | 0 | 39 | |
| A0A318CVA4 | 2026735 | 68525 | -0.132415 | 0 | NaN | 0 | 0 | 0 | 29 | |
| S7TLS7 | 1121405 | 68525 | 2.61647 | 1 | 15.376 | 60 | 1 | 2 | 87 | |
| A0A1J1E426 | 1725232 | 68525 | 0.835281 | 1 | NaN | 0 | 0 | 1 | 64 | |
| A9FZR5 | 448385 | 68525 | 4.92767 | 1 | 12.741 | 70 | 1 | 2 | 97 | |
| A0A0F6SHL5 | 927083 | 68525 | 5.96775 | 1 | 12.83 | 56 | 1 | 2 | 88 | |
| C7LVB0 | 525897 | 68525 | 1.13879 | 1 | NaN | 0 | 0 | 1 | 54 | |
| H8MMS9 | 1144275 | 68525 | 2.80411 | 1 | 12.23 | 45 | 1 | 2 | 61 | |
| A0A2M9DKA4 | 157842 | 68525 | 3.10973 | 1 | 20.825 | 77 | 1 | 2 | 93 | |
| A0A2Z4FPE4 | 1548548 | 68525 | 1.77036 | 1 | NaN | 0 | 0 | 1 | 45 | |
| B8FAA1 | 218208 | 68525 | 1.55685 | 1 | 15.489 | 46 | 1 | 2 | 60 | |
| J0UXX0 | 1177931 | 68525 | 2.83321 | 1 | 30.249 | 56 | 1 | 2 | 82 | |
| Q1MR30 | 363253 | 68525 | 0.588158 | 1 | NaN | 0 | 0 | 1 | 48 | |
| A0A1H0F191 | 206665 | 68525 | -0.0484183 | 0 | NaN | 0 | 0 | 0 | 43 | |
| D0LVJ8 | 502025 | 68525 | 4.92032 | 1 | 12.475 | 46 | 1 | 2 | 80 | |
| E1QIQ1 | 644282 | 68525 | 3.13612 | 1 | 21.248 | 33 | 1 | 2 | 53 | |
| I4C0I0 | 706587 | 68525 | 1.00097 | 1 | NaN | 0 | 0 | 1 | 48 | |
| F2LV66 | 760142 | 68525 | 0.349448 | 0 | NaN | 0 | 0 | 0 | 31 | HAS ORTHOLOG WITH LARGER LLPS PROPENSITY |
| F2LU75 | 760142 | 68525 | 0.426682 | 0 | NaN | 0 | 0 | 0 | 26 | |
| Q2LVL9 | 56780 | 68525 | 0.490505 | 0 | NaN | 0 | 0 | 0 | 36 | |
| C8PFI3 | 553220 | 68525 | 1.43483 | 1 | 19.121 | 35 | 1 | 2 | 56 | HAS ORTHOLOG WITH LARGER LLPS PROPENSITY |
| C8PLH8 | 553220 | 68525 | 2.94303 | 1 | 17.491 | 60 | 1 | 2 | 94 | |
| C0QIM1 | 177437 | 68525 | 3.62437 | 1 | 19.984 | 54 | 1 | 2 | 65 | |
| Q2IG69 | 290397 | 68525 | 3.14623 | 1 | NaN | 0 | 0 | 1 | 58 | |
| A0A099TYB0 | 216 | 68525 | 0.957846 | 1 | 16.976 | 59 | 1 | 2 | 99 | |
| Q747Z8 | 243231 | 68525 | 2.05876 | 1 | NaN | 0 | 0 | 1 | 41 | |
| Q74B90 | 243231 | 68525 | 0.871856 | 1 | NaN | 0 | 0 | 1 | 34 | HAS ORTHOLOG WITH LARGER LLPS PROPENSITY |
| A0A1I4T4Z9 | 39841 | 68525 | 0.397916 | 0 | NaN | 0 | 0 | 0 | 47 | |
| O69302 | 192222 | 68525 | 1.15957 | 1 | 19.443 | 57 | 1 | 2 | 81 | |
| O25841 | 85962 | 68525 | 0.20207 | 0 | 11.816 | 56 | 1 | 1 | 76 | |
| P59933 | 273121 | 68525 | 0.738118 | 1 | NaN | 0 | 0 | 1 | 59 | |
| A0A0S3QVU2 | 1298851 | 200783 | 0.340415 | 0 | NaN | 0 | 0 | 0 | 33 | HYPERTH |
| F0S465 | 868864 | 200783 | 0.179576 | 0 | NaN | 0 | 0 | 0 | 20 | HYPERTH MOTIF MISSING, FRAGMENT? |
| C1DTM9 | 204536 | 200783 | 0.593467 | 1 | NaN | 0 | 0 | 1 | 35 | |
| O66475 | 224324 | 200783 | 0.414218 | 0 | NaN | 0 | 0 | 0 | 45 | HYPERTH |
| A0A0G2ZH51 | 1330330 | 200918 | 0.398683 | 0 | NaN | 0 | 0 | 0 | 45 | |
| A0A0C7NPU9 | 1006576 | 200918 | 0.29381 | 0 | NaN | 0 | 0 | 0 | 72 | |
| Q9WZ73 | 243274 | 200918 | -0.789215 | 0 | NaN | 0 | 0 | 0 | 36 | HYPERTH |
| A0A383S6K9 | 119981 | 201174 | 1.09343 | 1 | 16.825 | 51 | 1 | 2 | 77 | |
| F8AYN8 | 656024 | 201174 | -0.56604 | 0 | NaN | 0 | 0 | 0 | 49 | |
| A0A132ML03 | 1469144 | 201174 | 1.55411 | 1 | NaN | 0 | 0 | 1 | 56 | |
| A0A239VAG6 | 1863 | 201174 | 2.19751 | 1 | 10.614 | 68 | 1 | 2 | 83 | |
| A0A346Y532 | 1608957 | 201174 | 0.689242 | 1 | NaN | 0 | 0 | 1 | 43 | HAS ORTHOLOG WITH LARGER LLPS PROPENSITY |
| A0A346Y5S5 | 1608957 | 201174 | 0.741908 | 1 | NaN | 0 | 0 | 1 | 40 | |
| D7GH49 | 754252 | 201174 | 1.75855 | 1 | 21.415 | 52 | 1 | 2 | 81 | |
| A0A1G9N6L8 | 380244 | 201174 | 1.88294 | 1 | NaN | 0 | 0 | 1 | 64 | |
| A0A0M2H3M8 | 92835 | 201174 | 0.95685 | 1 | NaN | 0 | 0 | 1 | 61 | HAS ORTHOLOG WITH LARGER LLPS PROPENSITY |
| A0A0M2GYF5 | 92835 | 201174 | 1.04259 | 1 | NaN | 0 | 0 | 1 | 94 | |
| D6ZB10 | 640132 | 201174 | 1.10323 | 1 | NaN | 0 | 0 | 1 | 59 | |
| A0A239IUZ1 | 1945885 | 201174 | 0.918772 | 1 | NaN | 0 | 0 | 1 | 56 | |
| A0A3A3YS05 | 2339232 | 201174 | 3.33992 | 1 | 11.114 | 55 | 1 | 2 | 81 | |
| C7R3A8 | 471856 | 201174 | 2.22396 | 1 | 18.113 | 49 | 1 | 2 | 81 | |
| A0A0M4MXW0 | 1528099 | 201174 | 1.15485 | 1 | NaN | 0 | 0 | 1 | 64 | |
| A0A1G6HDQ7 | 1577474 | 201174 | 2.62336 | 1 | 16.787 | 77 | 1 | 2 | 91 | |
| A0A1M7RKJ2 | 134849 | 201174 | 1.54803 | 1 | NaN | 0 | 0 | 1 | 62 | |
| A0A3Q8WUB9 | 1282737 | 201174 | 2.33019 | 1 | 9.711 | 39 | 1 | 2 | 70 | |
| A0A1M5IQR5 | 1206085 | 201174 | 1.43267 | 1 | NaN | 0 | 0 | 1 | 62 | |
| A0A2T0ZYT7 | 1629062 | 201174 | 1.2701 | 1 | NaN | 0 | 0 | 1 | 52 | |
| A0A075JDJ9 | 1274 | 201174 | 3.94493 | 1 | 17.197 | 87 | 1 | 2 | 112 | |
| A0A3N1GWX3 | 763993 | 201174 | 3.55171 | 1 | 20.016 | 77 | 1 | 2 | 87 | |
| A0A3D9V509 | 696763 | 201174 | 0.818799 | 1 | NaN | 0 | 0 | 1 | 55 | |
| A0A1L2ZNP0 | 556325 | 201174 | 1.97943 | 1 | 15.921 | 44 | 1 | 2 | 81 | |
| D3F3T6 | 469383 | 201174 | 0.655395 | 1 | NaN | 0 | 0 | 1 | 50 | |
| A0A1H0Z8H6 | 1881057 | 201174 | 4.16857 | 1 | 10.116 | 78 | 1 | 2 | 93 | |
| C7Q4G0 | 479433 | 201174 | 2.4453 | 1 | 19.78 | 44 | 1 | 2 | 68 | HAS ORTHOLOG WITH LARGER LLPS PROPENSITY |
| C7PVD3 | 479433 | 201174 | 4.95054 | 1 | 14.692 | 76 | 1 | 2 | 112 | |
| A0A060JHL8 | 529884 | 201174 | 0.0670081 | 0 | NaN | 0 | 0 | 0 | 64 | |
| A0A1A9GJ93 | 1300347 | 201174 | 2.42423 | 1 | 16.443 | 75 | 1 | 2 | 104 | |
| A0A329QRZ0 | 1981511 | 201174 | 0.495389 | 0 | NaN | 0 | 0 | 0 | 65 | |
| A0LWU4 | 351607 | 201174 | 0.225135 | 0 | NaN | 0 | 0 | 0 | 45 | |
| A0A1Y2MR47 | 2074 | 201174 | 3.215 | 1 | 11.895 | 42 | 1 | 2 | 79 | |
| Q6ABW8 | 281090 | 201174 | 0.60335 | 1 | NaN | 0 | 0 | 1 | 63 | |
| A0A3P1VMS4 | 2491050 | 201174 | 0.342669 | 0 | NaN | 0 | 0 | 0 | 70 | |
| X8BJS0 | 1299333 | 201174 | 1.73643 | 1 | NaN | 0 | 0 | 1 | 62 | |

| | | | | | | | | | | |
|---|---|---|---|---|---|---|---|---|---|---|
| Q47K95 | 269800 | 201174 | 2.73996 | 1 | 16.074 | 38 | 1 | 2 | 75 | |
| C7M1H0 | 525909 | 201174 | 0.144961 | 0 | NaN | 0 | 0 | 0 | 42 | |
| A0A3P3VUB7 | 2488791 | 201174 | 0.297812 | 0 | NaN | 0 | 0 | 0 | 56 | HAS ORTHOLOG WITH LARGER LLPS PROPENSITY |
| A0A3P3VW09 | 2488791 | 201174 | 1.73069 | 1 | NaN | 0 | 0 | 1 | 79 | |
| A4XDG7 | 369723 | 201174 | 1.567 | 1 | NaN | 0 | 0 | 1 | 62 | |
| E6S7C6 | 710696 | 201174 | 3.06285 | 1 | 15.765 | 97 | 1 | 2 | 110 | |
| B2GG11 | 378753 | 201174 | 1.56257 | 1 | NaN | 0 | 0 | 1 | 130 | |
| A0A255HB87 | 2016507 | 201174 | 3.33724 | 1 | 20.133 | 60 | 1 | 2 | 86 | |
| D5UP28 | 521096 | 201174 | 1.63992 | 1 | 14.247 | 35 | 1 | 2 | 70 | |
| A0A1J7BFQ6 | 1428644 | 201174 | 3.69451 | 1 | 13.068 | 78 | 1 | 2 | 98 | |
| A0A2X0VKA6 | 1658 | 201174 | 2.14315 | 1 | 23.651 | 49 | 1 | 2 | 83 | |
| A0A1T3NZ26 | 159449 | 201174 | 3.17199 | 1 | 19.407 | 78 | 1 | 2 | 93 | |
| A0A1T3NYA0 | 159449 | 201174 | 1.25816 | 1 | NaN | 0 | 0 | 1 | 64 | HAS ORTHOLOG WITH LARGER LLPS PROPENSITY |
| D2Q3W6 | 479435 | 201174 | 2.66123 | 1 | 9.964 | 59 | 1 | 2 | 85 | |
| A0A3N2BCG4 | 56055 | 201174 | 2.35258 | 1 | 18.213 | 39 | 1 | 2 | 70 | |
| A0A150H5L8 | 479117 | 201174 | 2.97742 | 1 | 19.569 | 68 | 1 | 2 | 94 | |
| A0A3N2DA89 | 120377 | 201174 | 2.84827 | 1 | 13.681 | 46 | 1 | 2 | 77 | |
| S2W6P3 | 883161 | 201174 | 1.04406 | 1 | NaN | 0 | 0 | 1 | 77 | |
| A0A087VSW7 | 1341694 | 201174 | 2.13702 | 1 | 14.672 | 55 | 1 | 2 | 90 | |
| A0A087VUZ9 | 1341694 | 201174 | 1.33333 | 1 | NaN | 0 | 0 | 1 | 73 | HAS ORTHOLOG WITH LARGER LLPS PROPENSITY |
| A0A073B3Y4 | 28042 | 201174 | 0.710609 | 1 | NaN | 0 | 0 | 1 | 49 | |
| A0A022LTY2 | 1292020 | 201174 | 2.81654 | 1 | 21.728 | 37 | 1 | 2 | 78 | |
| A0A419I132 | 2340915 | 201174 | 1.25185 | 1 | NaN | 0 | 0 | 1 | 60 | |
| Q1B002 | 266117 | 201174 | 1.84891 | 1 | NaN | 0 | 0 | 1 | 41 | |
| A0A3G8ZQ20 | 1902245 | 201174 | 1.61187 | 1 | NaN | 0 | 0 | 1 | 62 | |
| A0A249KIF8 | 1884904 | 201174 | 0.20764 | 0 | NaN | 0 | 0 | 0 | 58 | |
| A0A212THA9 | 592308 | 201174 | 2.10394 | 1 | 12.6 | 68 | 1 | 2 | 93 | |
| A0A411YFM3 | 1670831 | 201174 | 0.52806 | 1 | NaN | 0 | 0 | 1 | 45 | |
| A0A2N9JKW3 | 75385 | 201174 | 2.09774 | 1 | 16.747 | 44 | 1 | 2 | 69 | |
| D6YAJ5 | 469371 | 201174 | 1.18569 | 1 | NaN | 0 | 0 | 1 | 63 | |
| A0A0W1KKS1 | 59561 | 201174 | 1.66034 | 1 | 17.154 | 56 | 1 | 2 | 69 | |
| A0A1Q5PT88 | 52770 | 201174 | 3.03657 | 1 | 19.871 | 47 | 1 | 2 | 82 | |
| A0A2P4UMR4 | 1926885 | 201174 | 4.01991 | 1 | 16.003 | 72 | 1 | 2 | 85 | |
| A0A1B1NCZ9 | 1758689 | 201174 | 2.80251 | 1 | 22.108 | 73 | 1 | 2 | 98 | |
| D6ZJY0 | 548479 | 201174 | 1.25781 | 1 | NaN | 0 | 0 | 1 | 75 | |
| A0A095WU40 | 1219581 | 201174 | 1.44922 | 1 | 12.525 | 42 | 1 | 2 | 71 | |
| C8WA62 | 521095 | 201174 | 0.322331 | 0 | NaN | 0 | 0 | 0 | 35 | |
| C8W8M1 | 521095 | 201174 | 0.297779 | 0 | NaN | 0 | 0 | 0 | 45 | HAS ORTHOLOG WITH LARGER LLPS PROPENSITY |
| C5C7V0 | 465515 | 201174 | 3.33908 | 1 | 14.277 | 58 | 1 | 2 | 93 | |
| S3YJM5 | 1203568 | 201174 | 2.8901 | 1 | 18.577 | 52 | 1 | 2 | 82 | |
| F8B488 | 656024 | 201174 | -0.482664 | 0 | NaN | 0 | 0 | 0 | 55 | |
| H6RVY1 | 1146883 | 201174 | 2.06361 | 1 | NaN | 0 | 0 | 1 | 56 | |
| F8B065 | 656024 | 201174 | 2.45409 | 1 | 11.156 | 46 | 1 | 2 | 79 | |
| A0A3N6ZGY8 | 1538144 | 201174 | 2.06957 | 1 | 13.574 | 46 | 1 | 2 | 69 | |
| A0A255EF71 | 2016505 | 201174 | 2.45329 | 1 | 12.847 | 45 | 1 | 2 | 67 | |
| A0A1R4FAQ5 | 1434822 | 201174 | 2.58742 | 1 | 14.839 | 64 | 1 | 2 | 77 | |
| A0A239QRU7 | 1945888 | 201174 | 2.0328 | 1 | 17.551 | 41 | 1 | 2 | 72 | |
| A0A1H6ZYK6 | 1043493 | 201174 | 1.83241 | 1 | 12.679 | 36 | 1 | 2 | 63 | |
| W1TLM5 | 1403948 | 201174 | 1.34294 | 1 | 15.526 | 49 | 1 | 2 | 85 | |
| Q83N34 | 203267 | 201174 | 0.613233 | 1 | NaN | 0 | 0 | 1 | 81 | |
| Q9X8U3 | 100226 | 201174 | 3.86489 | 1 | 13.888 | 79 | 1 | 2 | 92 | |
| Q8NLG0 | 196627 | 201174 | 3.30591 | 1 | 26.697 | 95 | 1 | 2 | 117 | |
| P9WGD5 | 83332 | 201174 | 0.748605 | 1 | NaN | 0 | 0 | 1 | 57 | |
| A0A368KU82 | 1031537 | 203682 | 2.50467 | 1 | NaN | 0 | 0 | 1 | 66 | |
| A0A142XSZ2 | 1630693 | 203682 | 3.81768 | 1 | NaN | 0 | 0 | 1 | 59 | |
| A0A286RKX9 | 1331910 | 203682 | 0.748491 | 1 | NaN | 0 | 0 | 1 | 45 | |
| E8R1N7 | 575540 | 203682 | 1.48431 | 1 | NaN | 0 | 0 | 1 | 82 | |
| A0A142Y6P6 | 1632865 | 203682 | 0.791649 | 1 | NaN | 0 | 0 | 1 | 51 | |
| I0IET8 | 1142394 | 203682 | 3.03782 | 1 | NaN | 0 | 0 | 1 | 63 | |
| A0A1U9NHE5 | 1936003 | 203682 | 1.63518 | 1 | 22.968 | 33 | 1 | 2 | 50 | |
| A0A1W6LIZ2 | 1941349 | 203682 | 1.71473 | 1 | 25.75 | 45 | 1 | 2 | 63 | |
| A0A142WUX5 | 1632864 | 203682 | 2.72105 | 1 | NaN | 0 | 0 | 1 | 66 | |
| A0A1P8WK57 | 1891926 | 203682 | 1.43672 | 1 | NaN | 0 | 0 | 1 | 58 | |
| D5SQI8 | 521674 | 203682 | 2.3237 | 1 | NaN | 0 | 0 | 1 | 63 | |
| D2R6N7 | 530564 | 203682 | 1.72191 | 1 | NaN | 0 | 0 | 1 | 55 | |
| A0A3M8FSD8 | 2030824 | 203682 | 2.57382 | 1 | 11.393 | 33 | 1 | 2 | 46 | |
| A0A1E3X5M9 | 1872076 | 203682 | 0.590511 | 1 | NaN | 0 | 0 | 1 | 47 | |
| F0SN41 | 756272 | 203682 | 2.31717 | 1 | 17.49 | 39 | 1 | 2 | 68 | |
| A6CAL9 | 344747 | 203682 | 2.32668 | 1 | NaN | 0 | 0 | 1 | 56 | |
| A0A255R309 | 2023130 | 203682 | 0.878587 | 1 | NaN | 0 | 0 | 1 | 59 | |
| A0A1V6LZU5 | 392547 | 203682 | 0.65154 | 1 | NaN | 0 | 0 | 1 | 41 | |
| D6YW20 | 716544 | 204428 | 0.938603 | 1 | NaN | 0 | 0 | 1 | 40 | |
| F8L7J6 | 331113 | 204428 | 1.24434 | 1 | 9.915 | 41 | 1 | 2 | 66 | |
| O84048 | 272561 | 204428 | 1.09338 | 1 | NaN | 0 | 0 | 1 | 60 | |
| A4SCJ6 | 290318 | 109 | Excluded ortholog. | | | | | | | |
| D3EPD2 | 1453429 | 111 | Excluded ortholog. | | | | | | | |
| A0A2Z6E2H1 | 2010829 | 123 | Excluded ortholog. | | | | | | | |
| A0A1B1YSY7 | 1810504 | 123 | Excluded ortholog. | | | | | | | |
| A0A2X0WGW8 | 179995 | 123 | Excluded ortholog. | | | | | | | |
| A0A1H8VQC6 | 406100 | 123 | Excluded ortholog. | | | | | | | |
| A0A4Q5VPW9 | 1889775 | 123 | Excluded ortholog. | | | | | | | |
| A0A1H6F5D0 | 1899563 | 123 | Excluded ortholog. | | | | | | | |
| A0A1H6F8Q9 | 1899563 | 123 | Excluded ortholog. | | | | | | | |
| A0A1A8TLR0 | 295068 | 123 | Excluded ortholog. | | | | | | | |
| A0A1A8T993 | 295068 | 123 | Excluded ortholog. | | | | | | | |
| Q5ZWV8 | 272624 | 123 | Excluded ortholog. | | | | | | | |
| E1VAE8 | 768066 | 123 | Excluded ortholog. | | | | | | | |
| A0A1I0FP31 | 1123402 | 123 | Excluded ortholog. | | | | | | | |
| A5EUY7 | 246195 | 123 | Excluded ortholog. | | | | | | | |
| A0A1P8UGI6 | 1765967 | 123 | Excluded ortholog. | | | | | | | |
| S6GFH1 | 1330036 | 123 | Excluded ortholog. | | | | | | | |
| Q9RX92 | 243230 | 129 | Excluded ortholog. | | | | | | | |
| Q9RY80 | 243230 | 129 | Excluded ortholog. | | | | | | | |
| A0A0G2Z8X6 | 1330330 | 20091 | Excluded ortholog. | | | | | | | |
| A0A142XIH8 | 1630693 | 20368 | Excluded ortholog. | | | | | | | |
| A0A255R6T2 | 2023130 | 20368 | Excluded ortholog. | | | | | | | |
| D6YRN8 | 716544 | 20442 | Excluded ortholog. | | | | | | | |
| A0A2Z6I4G7 | 1885025 | 2821 | Excluded ortholog. | | | | | | | |
| D5AVD7 | 272942 | 2821 | Excluded ortholog. | | | | | | | |
| W6KQ01 | 1288970 | 2821 | Excluded ortholog. | | | | | | | |
| A0A420WD99 | 568099 | 2821 | Excluded ortholog. | | | | | | | |
| A0A318MVI2 | 2070537 | 2821 | Excluded ortholog. | | | | | | | |
| A0A366ES38 | 1473586 | 2821 | Excluded ortholog. | | | | | | | |
| A0A366FG47 | 1473586 | 2821 | Excluded ortholog. | | | | | | | |
| E0TAY4 | 314260 | 2821 | Excluded ortholog. | | | | | | | |
| I3TVB0 | 1110502 | 2821 | Excluded ortholog. | | | | | | | |
| A0A447IAU8 | 2490941 | 2821 | Excluded ortholog. | | | | | | | |
| A0A317ECA7 | 2211142 | 2821 | Excluded ortholog. | | | | | | | |
| A0A3L7J649 | 2448481 | 2821 | Excluded ortholog. | | | | | | | |
| A0A433SFE1 | 1965230 | 2821 | Excluded ortholog. | | | | | | | |
| A0A433SGA9 | 1965230 | 2821 | Excluded ortholog. | | | | | | | |
| E8UF33 | 937774 | 2821 | Excluded ortholog. | | | | | | | |
| A0A0A1H454 | 1469502 | 2821 | Excluded ortholog. | | | | | | | |
| A0A0A1H6Y8 | 1469502 | 2821 | Excluded ortholog. | | | | | | | |
| A0A0A1H3N2 | 1469502 | 2821 | Excluded ortholog. | | | | | | | |
| A0A0A1H1W5 | 1469502 | 2821 | Excluded ortholog. | | | | | | | |
| A0A0A1H2Y5 | 1469502 | 2821 | Excluded ortholog. | | | | | | | |
| A0A2I7N668 | 2052837 | 2821 | Excluded ortholog. | | | | | | | |
| A0A011Q4A3 | 1454000 | 2821 | Excluded ortholog. | | | | | | | |
| A6GUR1 | 391597 | 2821 | Excluded ortholog. | | | | | | | |
| D9SEK8 | 395494 | 2821 | Excluded ortholog. | | | | | | | |
| A0A0N0JSL7 | 1523428 | 2821 | Excluded ortholog. | | | | | | | |
| R7LTK7 | 1262901 | 3206 | Excluded ortholog. | | | | | | | |
| A0A0K1QED6 | 1391654 | 6852 | Excluded ortholog. | | | | | | | |
| A0A0N1J1J8 | 1509431 | 6852 | Excluded ortholog. | | | | | | | |
| A0A1J1DPC3 | 1725232 | 6852 | Excluded ortholog. | | | | | | | |
| A9G766 | 448385 | 6852 | Excluded ortholog. | | | | | | | |
| A0A0F6W6E1 | 927083 | 6852 | Excluded ortholog. | | | | | | | |
| A0A0F6YG51 | 927083 | 6852 | Excluded ortholog. | | | | | | | |

| | | | |
|---|---|---|---|
| A0A0F6SDT3 | 927083 | 6852 | Excluded ortholog. |
| A0A2Z4FN28 | 1548548 | 6852 | Excluded ortholog. |
| B8FNJ8 | 218208 | 6852 | Excluded ortholog. |
| A0A1T5HLH3 | 889453 | 97 | Excluded ortholog. |
| A0A2A3UF12 | 2021370 | 97 | Excluded ortholog. |
| A0A0C1L5P3 | 1463156 | 97 | Excluded ortholog. |
| A0A0C1KR02 | 1463156 | 97 | Excluded ortholog. |
| I4A267 | 867902 | 97 | Excluded ortholog. |
| U5Q7H7 | 1400053 | 97 | Excluded ortholog. |
| A0A0S7C0X3 | 1678841 | 97 | Excluded ortholog. |
| A0A0S7BQ47 | 1678841 | 97 | Excluded ortholog. |
| A0A0S7BX33 | 1678841 | 97 | Excluded ortholog. |
| F4KZC1 | 760192 | 97 | Excluded ortholog. |
| A0A0K8QUK7 | 1688776 | 97 | Excluded ortholog. |
| A0A1I2JES8 | 1003 | 97 | Excluded ortholog. |
| A0A1I2APS7 | 1003 | 97 | Excluded ortholog. |
| A0A1I1UF18 | 385682 | 97 | Excluded ortholog. |
| R7RV76 | 402612 | 97 | Excluded ortholog. |
| A0A142EQH3 | 1727163 | 97 | Excluded ortholog. |
| A0A0B8Y464 | 1122941 | 97 | Excluded ortholog. |
| A0A401U926 | 2482724 | 97 | Excluded ortholog. |
| G8R7E5 | 926562 | 97 | Excluded ortholog. |
| I2EVM7 | 929562 | 97 | Excluded ortholog. |
| A0A1H4BI92 | 908615 | 97 | Excluded ortholog. |
| A0A060RC04 | 1433126 | 97 | Excluded ortholog. |
| A0A2A3UFT5 | 2021370 | 97 | Excluded ortholog. |
| I4AL65 | 880071 | 97 | Excluded ortholog. |
| A0A346Y530 | 1608957 | 20117 | Excluded ortholog. |
| A0A346Y6N6 | 1608957 | 20117 | Excluded ortholog. |
| D7GCR2 | 754252 | 20117 | Excluded ortholog. |
| A0A0M2H287 | 92835 | 20117 | Excluded ortholog. |
| D6Z9K0 | 640132 | 20117 | Excluded ortholog. |
| A0A239IX48 | 1945885 | 20117 | Excluded ortholog. |
| A0A3A3ZDI3 | 2339232 | 20117 | Excluded ortholog. |
| A0A3A3Z1T3 | 2339232 | 20117 | Excluded ortholog. |
| A0A3A3YTY8 | 2339232 | 20117 | Excluded ortholog. |
| A0A239EH07 | 1945885 | 20117 | Excluded ortholog. |
| A0A239JP78 | 1945885 | 20117 | Excluded ortholog. |
| A0A0M4MKR0 | 1528099 | 20117 | Excluded ortholog. |
| A0A0M4MZU6 | 1528099 | 20117 | Excluded ortholog. |
| A0A0M4MLW5 | 1528099 | 20117 | Excluded ortholog. |
| A0A1G6H2Z0 | 1577474 | 20117 | Excluded ortholog. |
| A0A1M7RJJ8 | 134849 | 20117 | Excluded ortholog. |
| A0A1M7Q9Z5 | 134849 | 20117 | Excluded ortholog. |
| A0A3Q8WTS6 | 1282737 | 20117 | Excluded ortholog. |
| A0A3S8Z9T2 | 1282737 | 20117 | Excluded ortholog. |
| A0A1M5PMQ6 | 1206085 | 20117 | Excluded ortholog. |
| A0A2T0ZTN7 | 1629062 | 20117 | Excluded ortholog. |
| A0A2T0ZEM4 | 1629062 | 20117 | Excluded ortholog. |
| A0A2T0ZU92 | 1629062 | 20117 | Excluded ortholog. |
| A0A075JK41 | 1274 | 20117 | Excluded ortholog. |
| A0A1L2ZLU5 | 556325 | 20117 | Excluded ortholog. |
| A0A3D9V7I0 | 696763 | 20117 | Excluded ortholog. |
| A0A1H0XU41 | 1881057 | 20117 | Excluded ortholog. |
| A0A1H0XN94 | 1881057 | 20117 | Excluded ortholog. |
| C7Q5K4 | 479433 | 20117 | Excluded ortholog. |
| A0A060JLA2 | 529884 | 20117 | Excluded ortholog. |
| A0A1A9GQG8 | 1300347 | 20117 | Excluded ortholog. |
| A0A1A9GMG3 | 1300347 | 20117 | Excluded ortholog. |
| A0A1A9GKG3 | 1300347 | 20117 | Excluded ortholog. |
| A0A329QLM7 | 1981511 | 20117 | Excluded ortholog. |
| A0A329QRS9 | 1981511 | 20117 | Excluded ortholog. |
| A0A329QE27 | 1981511 | 20117 | Excluded ortholog. |
| A0LVD2 | 351607 | 20117 | Excluded ortholog. |
| A0A1Y2ML66 | 2074 | 20117 | Excluded ortholog. |
| A0A1Y2MLA3 | 2074 | 20117 | Excluded ortholog. |
| A0A1Y2MHE7 | 2074 | 20117 | Excluded ortholog. |
| A0A1Y2MLJ5 | 2074 | 20117 | Excluded ortholog. |
| X8BI06 | 1299333 | 20117 | Excluded ortholog. |
| X7ZHF1 | 1299333 | 20117 | Excluded ortholog. |
| C7M1D3 | 525909 | 20117 | Excluded ortholog. |
| A0A3P3VYR0 | 2488791 | 20117 | Excluded ortholog. |
| A0A132MWI0 | 1469144 | 20117 | Excluded ortholog. |
| E6S6A4 | 710696 | 20117 | Excluded ortholog. |
| A0A255H0N2 | 2016507 | 20117 | Excluded ortholog. |
| A0A255H2D0 | 2016507 | 20117 | Excluded ortholog. |
| A0A255H2P0 | 2016507 | 20117 | Excluded ortholog. |
| A0A1J7BF44 | 1428644 | 20117 | Excluded ortholog. |
| A0A1J7BCA8 | 1428644 | 20117 | Excluded ortholog. |
| A0A2X0TU18 | 1658 | 20117 | Excluded ortholog. |
| A0A1T3P1T5 | 159449 | 20117 | Excluded ortholog. |
| A0A1T3NYW9 | 159449 | 20117 | Excluded ortholog. |
| A0A1T3P216 | 159449 | 20117 | Excluded ortholog. |
| A0A1T3NJ33 | 159449 | 20117 | Excluded ortholog. |
| D2PTH5 | 479435 | 20117 | Excluded ortholog. |
| A0A3N2BF90 | 56055 | 20117 | Excluded ortholog. |
| A0A3N2DDF5 | 120377 | 20117 | Excluded ortholog. |
| A0A087VVB3 | 1341694 | 20117 | Excluded ortholog. |
| A0A073AZ78 | 28042 | 20117 | Excluded ortholog. |
| A0A022LPA2 | 1292020 | 20117 | Excluded ortholog. |
| A0A022LPM7 | 1292020 | 20117 | Excluded ortholog. |
| A0A419HHG3 | 2340915 | 20117 | Excluded ortholog. |
| A0A419I0Z7 | 2340915 | 20117 | Excluded ortholog. |
| A0A3G8ZV06 | 1902245 | 20117 | Excluded ortholog. |
| A0A3G8ZY59 | 1902245 | 20117 | Excluded ortholog. |
| A0A2N9JDE4 | 75385 | 20117 | Excluded ortholog. |
| D6Y4T4 | 469371 | 20117 | Excluded ortholog. |
| Q8NPF0 | 196627 | 20117 | Excluded ortholog. |
| A0A0W1KHB7 | 59561 | 20117 | Excluded ortholog. |
| A0A2P4URP4 | 1926885 | 20117 | Excluded ortholog. |
| A0A095WT15 | 1219581 | 20117 | Excluded ortholog. |
| C5CAV8 | 465515 | 20117 | Excluded ortholog. |
| F8B2W0 | 656024 | 20117 | Excluded ortholog. |
| F8B6R8 | 656024 | 20117 | Excluded ortholog. |
| A0A3N6WAQ6 | 1538144 | 20117 | Excluded ortholog. |
| A0A3N6W440 | 1538144 | 20117 | Excluded ortholog. |
| A0A3N6Z641 | 1538144 | 20117 | Excluded ortholog. |
| A0A255EGL7 | 2016505 | 20117 | Excluded ortholog. |
| A0A239QWP6 | 1945888 | 20117 | Excluded ortholog. |
| A0A1R4FWI9 | 1434822 | 20117 | Excluded ortholog. |
| A0A239QU82 | 1945888 | 20117 | Excluded ortholog. |
| A0A1R4F5T2 | 1434822 | 20117 | Excluded ortholog. |
| A0A1R4EWQ6 | 1434822 | 20117 | Excluded ortholog. |
| Q9KYI9 | 100226 | 20117 | Excluded ortholog. |
| P66856 | 203267 | 20117 | Excluded ortholog. |
| A0A0R2H3B9 | 1410657 | 123 | Excluded ortholog. |
| F9VI82 | 1029718 | 123 | Excluded ortholog. |
| F9VL12 | 1029718 | 123 | Excluded ortholog. |
| Q8XJT6 | 195102 | 123 | Excluded ortholog. |
| R6QE47 | 1262815 | 123 | Excluded ortholog. |
| A0A1E9AE52 | 1739304 | 123 | Excluded ortholog. |
| R5EL91 | 1263000 | 123 | Excluded ortholog. |
| Q2FWF7 | 93061 | 123 | Excluded ortholog. |
| A0A143WWB4 | 1780379 | 123 | Excluded ortholog. |
| A0A1V2YI65 | 1884656 | 123 | Excluded ortholog. |
| A0A1C0BVE8 | 1768196 | 123 | Excluded ortholog. |
| A0A1C0BTU4 | 1768196 | 123 | Excluded ortholog. |
| A0A1U7M557 | 1123403 | 123 | Excluded ortholog. |
| A0A136WD00 | 36847 | 123 | Excluded ortholog. |

| | | | |
|---|---|---|---|
| A0A087N112 | 1473546 | 123 | Excluded ortholog. |
| A0A1W1Z8Q2 | 371602 | 123 | Excluded ortholog. |
| K9E8I4 | 883081 | 123 | Excluded ortholog. |
| A0A0R3JXV0 | 908809 | 123 | Excluded ortholog. |
| A0A0R3JUI7 | 908809 | 123 | Excluded ortholog. |
| R6WJP8 | 1262797 | 123 | Excluded ortholog. |
| A0A1V4IYG7 | 1450648 | 123 | Excluded ortholog. |
| A0A1V4IGJ6 | 1450648 | 123 | Excluded ortholog. |
| A0A421BDS6 | 2315861 | 123 | Excluded ortholog. |
| A0A396QI02 | 2292273 | 123 | Excluded ortholog. |
| A0A1I4XZ97 | 398199 | 123 | Excluded ortholog. |
| A0A421BF91 | 2315861 | 123 | Excluded ortholog. |
| Q8R8F1 | 273068 | 123 | Excluded ortholog. |
| Q8RBI8 | 273068 | 123 | Excluded ortholog. |
| F2JIG6 | 642492 | 123 | Excluded ortholog. |
| H3NIX1 | 883113 | 123 | Excluded ortholog. |
| A0A1V4I8U7 | 29349 | 123 | Excluded ortholog. |
| A0A1S6ILM7 | 708126 | 123 | Excluded ortholog. |
| A0A1M5VUA2 | 1121316 | 123 | Excluded ortholog. |
| A0A1M5UVI1 | 1121316 | 123 | Excluded ortholog. |
| A0A095YRG8 | 1230734 | 123 | Excluded ortholog. |
| A0A2N6S620 | 2069309 | 123 | Excluded ortholog. |
| A0A0B4REG2 | 1526927 | 123 | Excluded ortholog. |
| A0A3A9IJN9 | 2320083 | 123 | Excluded ortholog. |
| A0A3A9HG68 | 2320083 | 123 | Excluded ortholog. |
| A0A3A9CQV5 | 2320083 | 123 | Excluded ortholog. |
| A0A1I5WFJ6 | 937334 | 123 | Excluded ortholog. |
| A0A143ZUS2 | 1780381 | 123 | Excluded ortholog. |
| A0A1Q9JV36 | 1261636 | 123 | Excluded ortholog. |
| A0A0X8G2G5 | 1712675 | 123 | Excluded ortholog. |
| A0A1M6GM39 | 1122184 | 123 | Excluded ortholog. |
| S0J792 | 1235790 | 123 | Excluded ortholog. |
| R6MUJ6 | 1263021 | 123 | Excluded ortholog. |
| R6QMS7 | 1262886 | 123 | Excluded ortholog. |
| R6MLJ5 | 1263021 | 123 | Excluded ortholog. |
| A0A2K9P6M4 | 1981510 | 123 | Excluded ortholog. |
| A0A2I0N053 | 1658742 | 123 | Excluded ortholog. |
| A0A410PU66 | 2507160 | 123 | Excluded ortholog. |
| H1BK61 | 457402 | 123 | Excluded ortholog. |
| B9YD18 | 545696 | 123 | Excluded ortholog. |
| A0A1B6BI81 | 1048380 | 123 | Excluded ortholog. |
| R7GJ76 | 1262795 | 123 | Excluded ortholog. |
| V2Z5V8 | 592026 | 123 | Excluded ortholog. |
| E7MQM1 | 706433 | 123 | Excluded ortholog. |
| A0A0P8Y6Z9 | 36849 | 123 | Excluded ortholog. |
| A0A0B4S1U2 | 33033 | 123 | Excluded ortholog. |
| A0A1G6Z5R6 | 2741 | 123 | Excluded ortholog. |
| R6C7H6 | 1262963 | 123 | Excluded ortholog. |
| A0A150M6U9 | 301148 | 123 | Excluded ortholog. |
| A0A3P2A9L6 | 2491044 | 123 | Excluded ortholog. |
| A0A3P2A9M9 | 2491044 | 123 | Excluded ortholog. |
| T0PJE5 | 1354301 | 123 | Excluded ortholog. |
| T0PNS3 | 1354301 | 123 | Excluded ortholog. |
| A0A417HP54 | 2292269 | 123 | Excluded ortholog. |
| A0A1Y4M2D0 | 1965572 | 123 | Excluded ortholog. |
| A0A140L3M2 | 520762 | 123 | Excluded ortholog. |
| B8I053 | 394503 | 123 | Excluded ortholog. |
| A0A1G5S531 | 1120920 | 123 | Excluded ortholog. |
| A0A143XWZ6 | 1780380 | 123 | Excluded ortholog. |
| A0A143XLA3 | 1780380 | 123 | Excluded ortholog. |
| A0A143XLP9 | 1780380 | 123 | Excluded ortholog. |
| A0A143XY36 | 1780380 | 123 | Excluded ortholog. |
| A0A267MIV8 | 1478221 | 123 | Excluded ortholog. |
| A0A3A9E627 | 2320089 | 123 | Excluded ortholog. |
| A0A3A9EDP4 | 2320089 | 123 | Excluded ortholog. |
| A0A3A9E3N5 | 2320089 | 123 | Excluded ortholog. |
| A0A3A9EEE5 | 2320089 | 123 | Excluded ortholog. |
| A0A3A9E170 | 2320089 | 123 | Excluded ortholog. |
| Q8Y4C7 | 169963 | 123 | Excluded ortholog. |
| A0A151ARE1 | 1121305 | 123 | Excluded ortholog. |
| A0A151ARY9 | 1121305 | 123 | Excluded ortholog. |
| A0A1M6NE77 | 1121331 | 123 | Excluded ortholog. |
| C0SPB6 | 224308 | 123 | Excluded ortholog. |
| Q8XNP5 | 195102 | 123 | Excluded ortholog. |
| A0A1H9M7H5 | 137733 | 123 | Excluded ortholog. |
| R5V5F3 | 1262953 | 123 | Excluded ortholog. |
| A0A1T4W7R7 | 39495 | 123 | Excluded ortholog. |
| A0A1T4V7N4 | 39495 | 123 | Excluded ortholog. |
| D5X2Q2 | 75379 | 2821 | Excluded ortholog. |
| A0A0F6W766 | 927083 | 6852 | Excluded ortholog. |
| A8ZYK1 | 96561 | 6852 | Excluded ortholog. |
| Q8NMV9 | 196627 | 20117 | Excluded ortholog. |
| S3YAW5 | 1203568 | 20117 | Excluded ortholog. |
| W1TM52 | 1403948 | 20117 | Excluded ortholog. |
| C7R1E8 | 471856 | 20117 | Excluded ortholog. |
| A0A132MZK2 | 1469144 | 20117 | Excluded ortholog. |
| A0A3P3VV67 | 2488791 | 20117 | Excluded ortholog. |
| A0A150H8K3 | 479117 | 20117 | Excluded ortholog. |
| A0A3N2DD55 | 120377 | 20117 | Excluded ortholog. |
| A0A1E7Z3B5 | 252393 | 123 | Excluded ortholog. |
| A0A348HEE9 | 33074 | 123 | Excluded ortholog. |
